# Sequence diversity in the 3’ untranslated region of alphavirus modulates IFIT2-dependent restriction in a cell type-dependent manner

**DOI:** 10.1101/2021.12.10.472177

**Authors:** Sarah E. Hickson, Eden Brekke, Johannes Schwerk, Indraneel Saluhke, Shivam Zaver, Joshua Woodward, Ram Savan, Jennifer L. Hyde

## Abstract

Alphaviruses (family *Togaviridae*) are a diverse group of positive-sense RNA (+ssRNA) viruses that are transmitted by arthropods and are the causative agent of several significant human and veterinary diseases. Interferon (IFN)-induced proteins with tetratricopeptide repeats (IFITs) are a family of RNA-binding IFN stimulated genes (ISGs) that are highly upregulated following viral infection, and have been identified as potential restrictors of alphaviruses. The mechanism by which IFIT1 restricts RNA viruses is dependent on self and non-self-discrimination of RNA, and alphaviruses evade this recognition via their 5’UTR. However, the role of IFIT2 during alphavirus replication and the mechanism of viral replication inhibition is unclear. In this study, we identify IFIT2 as a restriction factor for Venezuelan equine encephalitis virus (VEEV) and show that IFIT2 binds the 3’ untranslated region (3’UTR) of the virus. We investigated the potential role of variability in the 3’UTR of the virus affecting IFIT2 antiviral activity by studying infection with VEEV. Comparison of recombinant VEEV clones containing 3’UTR sequences derived from epizootic and enzootic isolates exhibited differential sensitivity to IFIT2 restriction *in vitro* infection studies, suggesting that the alphavirus 3’UTR sequence may function in part to evade IFIT2 restriction. *In vitro* binding assays demonstrate that IFIT2 binds to the VEEV 3’UTR, however in contrast to previous studies VEEV restriction did not appear to be dependent on the ability of IFIT2 to inhibit translation of viral RNA, suggesting a novel mechanism of IFIT2 restriction. Our study demonstrates that IFIT2 is a restriction factor for alphaviruses and variability in the 3’UTR of VEEV can modulate viral restriction by IFIT2. Ongoing studies are exploring the biological consequences of IFIT2-VEEV RNA interaction in viral pathogenesis and defining sequence and structural features of RNAs that regulate IFIT2 recognition.

## INTRODUCTION

Alphaviruses (family *Togaviridae*) are a diverse group of positive-sense RNA (+ssRNA) viruses that are transmitted by arthropods and are the causative agent of several significant human and veterinary diseases. Venezuelan equine encephalitis virus (VEEV) belongs to the VEE complex of alphaviruses and is responsible for periodic epizootic outbreaks of disease in equines in South, Central, and North America (reviewed in [1]). The VEE serocomplex consists of numerous different species which can be divided into six subtypes (I-VI). VEEV belongs to subtype I and can be further classified as epizootic (IAB and IC) or enzootic (ID and IE). During inter-epizootic periods VEEV is transmitted via a sylvatic cycle involving *Culex* spp. mosquitoes and rodents which serve as reservoirs for the virus. These enzootic subtypes (ID and IE) are considered equine avirulent and not associated with significant morbidity or mortality in these species. In contrast, epizootic variants (IAB and IC) are transmitted by several different species of mosquito and are associated with high titer viremia in equines which serve as amplification hosts during epizootic outbreaks. Epizootic subtypes are known to arise from mutation of sympatric enzootic viruses, and the epizootic phenotype has been shown to correlate with type-I interferon (IFN) resistance [2]. Genetic studies have previously mapped the key epizootic determinants to the E2 glycoprotein, which serves as the viral attachment protein [3–6]. Mutations in E2 sequences of epizootic subtypes have been shown to support higher levels of replication in horses and greater virulence, as well as adaptation to epizootic mosquito vectors [5, 7]. Although these mutations alone have been shown to be sufficient to confer epizootic phenotypes, epizootic subtypes also acquire mutations outside of this region, suggesting that additional genetic changes may contribute to the emergence of epizootic VEEV [8]. None of the studies have investigated the effect the sequence diversity in the 3’UTR of VEEV has on viral pathogenesis.

Alphaviruses are sensitive to type I IFN and IFN stimulated genes (ISGs) are known to be the effectors of alphavirus restriction. We have previously identified IFIT1 as a restriction factor for VEEV and other alphaviruses [9]. While IFIT1 recognizes non-self-viral RNA by preferentially binding to and restricting 7-methylguanisine (m7G) capped viral RNA (cap 0), alphaviruses are able to evade IFIT1 by encoding stable RNA structures within their 5’UTR [9–12]. However, the role of IFIT2 in alphavirus replication and pathogenesis has not yet been defined. Like IFIT1, IFIT2 also binds RNA [13] and has been shown to protect against lethal neuropathogenesis induced by diverse RNA viruses including flaviviruses, coronaviruses, and rhabdoviruses [14–20]. Despite this, the precise mechanisms by which IFIT2 inhibits replication and pathogenesis of viruses is unclear. Furthermore, IFIT2 has been implicated in diverse cellular processes, including RNA binding [13, 21], translational regulation [22, 23], cell death [24–28], cancer development [26, 29-33], and immune signaling [16, 34-36]. Although IFIT2 has predominantly been demonstrated to function as a restriction factor *in vitro* and *in vivo* in models of viral infection, Tran et al. recently demonstrated that recruitment of IFIT2 to influenza A virus mRNA in fact promotes translation and replication of viral mRNAs, implicating a proviral role for IFIT2 during influenza virus replication [37].

Here, we identify IFIT2 as a restriction factor for VEEV using a shRNA-based screening approach and show that IFIT2 also protects mice from VEEV pathogenesis *in vivo*. We show that IFIT2 binds AU-rich RNA located in the 3’ end of the viral genome, and that changes in the 3’UTR sequence alter the sensitivity of VEEV to IFIT2-dependent inhibition in a cell type-dependent manner. We further show that replication of viruses encoding 3’UTR sequences from epizootic and enzootic VEEV subtypes are differentially affected in an IFIT2-dependent and -independent manner, suggesting that IFIT2 recognition of viral RNA is likely modulated by RNA structure. Lastly, we show that the mechanism by which IFIT2 inhibits VEEV is independent of viral RNA translation and suggest possible mechanisms by which IFIT2 restricts alphavirus replication.

## MATERIALS AND METHODS

### Lentiviral transduction and flow cytometry

HeLa cells were transduced with lentiviruses encoding shRNA. The bicistronic vector (pGIPZ) co-expresses the shRNA and GFP, which are driven by IRES and cytomegalovirus immediate early promoters, respectively. Individual shRNA constructs (control (NSC, catalog # RHS4346), and human STAT2 and human IFIT1 (**Table S1**)) were packaged into lentiviral vectors following the manufacturer’s instruction (Open Biosystems). Two days post-transduction, cells were treated with 10 IU/mL of human IFN-β (PBL interferon source) for 6 hours and then infected with TC83 at a MOI of 1. Twenty-four hours post-infection cells were trypsinized, washed once in Hanks Balanced salt solution (HBSS), fixed in 1% paraformaldehyde for 10 minutes at room temperature, and then permeabilized in permeabilization buffer (HBSS, 10 mM HEPES pH 7.3, and 0.1% saponin). Cells were subsequently with incubated with antibodies specific for VEEV [38] in permeabilization buffer at room temperature for 20 minutes. After three washes, cells were incubated with species-specific Alexa Fluor 647-conjugated secondary antibodies (Molecular Probes) in permeabilization buffer. Cells were analyzed for GFP expression (shRNA expression) and viral antigen by flow cytometry on a BD FACS Array flow cytometer (BD Biosciences). Data were processed using FlowJo analysis software (Tree Star, Inc.). The initial preliminary shRNA screen was performed in duplicate two times independently, and the fold increase in VEEV positive cells was calculated by comparing data to a non-silencing control shRNA, and z-scores calculated. Based on this data a subset of genes was chosen for further validation based on the following criteria: i) hits with a z-score >2, ii) hits for which more than two shRNAs against the same gene exhibited 2-fold or greater VEEV positive cells relative to shNSC; iii). The validation screen was performed as described above using 10 IU/mL of IFN-β stimulation, and was performed in quadruplicate 4 times independently.

### Antibodies, and cell lines

Vero C1008 and Raw264.7 cells were obtained from ATCC. Primary murine embryonic fibroblasts (MEF) were generated from E15 to 16 WT and *Ifit2^-/-^* mouse embryos. All cell lines were maintained in DMEM supplemented with 10% defined heat-inactivated FBS (HyClone), L-glutamax, and nonessential amino acids. Mouse anti-VEEV E2 (clone 36.E5) and mouse anti-ISG54 (clone 6.H9) antibodies were produced and purified from a clonal hybridoma cell line and was a generous gift of Dr. Michael Diamond (Washington University School of Medicine, St Louis).

### Generation of Raw264.7 *Ifit2^-/-^* CRISPR cells

A doxycycline-inducible CRISPR/Cas9 expression vector (pSBtet-puro-Cas9-U6) was generated by cloning the Cas9-U6 portion of pX459 (Addgene #62988; [39] into pSBtet-pur (Addgene #60507; [40]). Cas9 was first cloned into pSBtet-pur using the following primers: Cas9.F: 5’-CATGAGACCGGTGCCACCATG-3’, Cas9.R: 5’-CATGAGGCGGCCGCCTACTTTTTCTTTTTTGCCTGGCCG, pSBtet-pur. F: 5’-CATGAG GCGGCCGCCTTCC-3’, pSBtet-pur. R: 5’-CATGAGACCGGTGGTGGCCGATATCTCAGAG. Cas9 was ligated into the pSBtet-pur backbone using 5’ AgeI and 3’ NotI restriction sites. The U6 promoter was then cloned into this new plasmid using the following primers: U6.F 5’-ACTACAGGTACC GAGGG-3’, U6.R 5’-TCAGTCCTAGGTCTAGAGC-3’, pSBtet-pur-Cas9.F 5’-TCAGTCCTAGGTCTAGAGC-3’, pSBtet-pur-Cas9.R 5’-ATGAAGGTACCACATTTGTAGAGGTTTTACTTGC-3’. U6 was ligated into pSBtet-pur-Cas9 using 5’ KpnI and 3’ AvrII restriction sites. As the new pSBtet-pur-Cas9-U6 plasmid contained an addition BbsI site, this was remove using site directed mutagenesis and the following primers: dBbsI.F 5’-TTGG GAAGAT AATAGCAG-3’, dBbsI.R 5’-CTGCTATTATCTTCCCAA-3’. Sequence-specific gRNA sequences were designed using the Broad Institute Genetic Perturbation Platform gRNA design tool to target mouse Ifit2 (accession # NM_008332). The following primers were used to generate Ifit2 gRNA oligonucleotides which were cloned into pSBtet-puro-Cas9-U6 as described previously [39]: Ifit2.g13.F: 5’-CACCGCACTGCAGAGGTCTAAATG-3’, Ifit2.g13.R: 5’-AAACCATTTAGACCTCTGCAGTGC-3’; Ifit2.g25.F: 5’-CACCGATCAGAAGTCTGGTCACCTG-3’, Ifit2.g25.R: 5’-AAACCAGGTGACCAGACTTCTGATC-3’; Ifit2.g38.F: 5’-CACCGCAGGATTCTCAATCCTGTAG-3’, Ifit2.g38.R: 5’-AAACCTACAGGATTGAGAATCCTGC-3’. Raw264.7 Ifit2 CRISPR cells were generated by electroporating low passage Raw264.7 cells with Ifit2 pSBtet-puro-Cas9-U6 using Amaxa Nucleofector II and Amaxa Cell Line Nucleofector Kit V (Lonza). Cells were selected with puromycin 3 days post-nucleofection, and Cas9/gRNA expression induced at 7 days post-nucelofection. Cells were treated for 7-14 days with doxycycline and KO efficiency of bulk cells validated using western blotting and detection with mouse anti-ISG54 (6.H9).

### Generation of full-length and recombinant viruses

Construction of the full length TC83 VEEV infectious clone has been described [6]. The following VEEV epizootic and enzootic 3’UTR sequences were introduced into the TC83 infectious clone: (IC) TGAACATAGCAGCAATTGGCAAGCTGCTTATATAGAACTTGCGGCGATTGGCATGCCGCTTTAAAATTTTATTTTATTTTCTTTTCTTTTCCGAATCGGATTTTGTTTTTAATATTTC; (ID) TGAACATAGCAGCAATTGGCAAGCTGCTTATATAGAACTCGCGGCGATTGGCATGCCGCTTTAAAATTTTATTTTATTTTCTTTTCTTTTCCGAATCGGATTTTGTTTTTAATATTTC; (IE) TAGAATTAGCAGCGATTGGCATGCTGCTTGTAAAGTTTTATTACAAATAACGTGCGGCAATTGGCGAGCCGCTTTAATTAGAATTTTATTTTCTTTTACCATAATCGGATTTTGTTTTTAATATTTC. Point mutations were introduced by site-directed mutagenesis using Phusion High-Fidelity PCR kit (NEB) using primers listed in **Table S1**.

Plasmids were linearized at MluI restriction sites located downstream of the poly(A) tail and genomic RNA was transcribed from the SP6 promoter in the presence of N7^m^G cap analog using the SP6 mMessage mMachine kit (Ambion). 1×10^7^ BHK21 cells were electroporated with approximately 2 μg of *in vitro* transcribed RNA using a GenePulser Xcell electroporator (Bio-Rad) to generate P0 virus stocks.

### Focus-forming assays

Focus-forming assays (FFAs) were performed in 96-well plates by adapting an assay developed for flaviviruses [41]. Vero E6 monolayers were infected with serial 10-fold dilutions of infectious samples for 1 hour at 37oC, then overlaid with 100 μl per well of medium (0.5x DMEM, 5% FBS) containing 1% carboxymethylcellulose, and incubated for 20 to 22 hours at 37oC with 5% CO_2_. Cells were then fixed by adding 100 μl per well of 2% paraformaldehyde directly onto the overlay at room temperature for 2 hours. After removal of overlay media and fixative, cells were washed 3x with PBS and incubated with antibodies specific for VEEV E2 glycoprotein (gift of Dr. Michael Diamond; see above) for 2 hours at room temperature in FFA permeabilization buffer (1x PBS, 0.1% saponin, and 0.1% BSA). Cells were washed 3x in ELISA wash buffer (1x PBS, 0.05% triton X-100), then incubated with species-specific HRP-conjugated secondary antibodies (Sigma and ThermoFisher) for 1 hour at room temperature in FFA permeabilization buffer. Monolayers were washed 3x with ELISA buffer and foci were developed by incubating in 50 μl/well of TrueBlue peroxidase substrate (KPL) for 5 to 10 minutes at room temperature, after which time cells were washed twice in water. Well images were captured using Immuno Capture software (Cell Technology Ltd.), and foci counted using BioSpot software (Cell Technology Ltd.). All samples were titered in duplicate and calculated titers averaged for each duplicate.

### Mouse experiments

All mouse experiments were performed in compliance and with the approval of the Washington University School of Medicine, and University of Washington Animal Studies Committees. C57BL/6 wild type mice were obtained from Jackson laboratories, and congenic *Ifit2^-/-^* mice were generated previously [42] (gift of Dr. Ganes Sen) [11] [11] [11]. Sixteen week-old male and female mice were used in all *in vivo* infection experiments. For subcutaneous infection, 10^2^ FFU of ZPC738 was diluted in phosphate buffered saline (PBS) in a total volume of 50μL, and delivered via injection into the rear footpad.

### Viral growth kinetic assays

MEF and Raw264.7 cells were maintained in DMEM supplemented with 10% defined heat-inactivated FBS, L-glutamine, and non-essential amino acids. Multistep viral growth curves were performed by infecting MEFs with WT or mutant VEEV TC83 viruses as indicated at a MOI of 0.01 in the presence or absence of IFN-β (PBL Interferon Source) pre-treatment. Cell supernatants were serially harvested at indicated time points post-infection by removing cell culture supernatant and replacing with fresh growth media, and viral titers determined by FFA. All experiments were performed three times independently in triplicate. Statistical analysis was performed by calculating area under the curve (AUC) and performing unpaired t-test on AUC values calculated for each experiment. P values are reported in each figure.

### Translation assays

Construction of the VEEV 3’UTR translation reporters was similar to that described previously [9, 43]. Mutations derived from epizootic and enzootic 3’UTR sequences were introduced by restriction digestion and ligation. The following primers were used to amplify 3’UTR sequences from infectious clones (described above): VEEV F: TAGTTAATTAAGATCAGCCGTAATTATTATAATTGG VEEV R: GCGGCCGCTCAAGAATTAATTCCCctcgaC. Fragments were amplified using Q5 high fidelity polymerase (NEB), purified using AMPure XP beads (Beckman Coulter), digested with PacI and NotI (NEB), and ligated into the VRLF vector which was similarly digested. Reporter RNAs were *in vitro* transcribed and co-transcriptionally capped using the mMESSAGE mMACHINE T7 kit (Ambion) according to the manufacturer’s instructions, purified using RNeasy Mini Kit (Qaigen). WT and *Ifit2^-/-^* MEF were treated with 0 or 10 IU/ml of IFN-β (PBL Interferon Source) for 6 hours. Reporter assays were performed as previously described [9, 44] using 2 μg (MEF) or 10 μg (Raw264.7) of RNA for each electroporation reaction and cells were plated in 96-well plates. Cells were collected at indicated time points by centrifugation at 1500 x g for 3 minutes, washed 1x in DPBS (Gibco), and lysed in 1x cell culture lysis (CCL) buffer (Promega). Firefly luciferase assays were performed on 20 μL of cell lysates using Luciferase Assay System (Promega) according to the manufactures instructions. Luciferase activity was measured on a BioTek Synergy microplate luminometer. Relative light unit values were normalized to protein concentration using a standard BCA Protein assay (Pierce).

### Generation of mouse recombinant IFIT2 expression vector

An expression vector encoding human IFIT2 (hIFIT2) (pET28a(+)-6xHis-TEV-IFIT2, Addgene; [45]) was modified to express mouse IFIT2 (mIFIT2). A synthetic gene fragment encoding mIFIT2 (accession number NM_008332) flanked by NheI (5’) and XhoI (3’) restriction sites was synthesized (IDT). The hIFIT2 sequence was removed from pET28a(+)-6xHis-TEV-IFIT2 by digestion with NheI and XhoI, and mIFIT2 ligated into the corresponding sites, to generate the new pET28a(+)-6xHis-TEV-mIFIT2 expression plasmid. Plasmid was transformed into 10-beta cells (NEB) and clone sequence verified by sanger sequencing.

### Recombinant Protein Expression and Purification

pET28a(+)-6xHis-TEV-mIFIT2 plasmid encoding mouse IFIT2 was transformed into Rosetta (DE3) pLysS chemically competent cells. Stationary phase cultures of the resulting transformed bacteria were used to inoculate 1.5 L of LB broth at a 1:100 dilution. Bacterial cultures were grown to OD600 0.4 at 37°C after which protein expression was induced by the addition of 0.5 mM isopropyl β-D-1-thiogalactopyranoside (IPTG) for 20 hours at 16°C. Bacteria were harvested by centrifugation at 7,000 x g for 30 minutes, and the cell pellets were resuspended in 30 mL of Buffer A [20 mM HEPES, pH 7.3, 1.5 M NaCl, 2 mM MgCl2, 25 mM Imidazole, 10% Glycerol, supplemented with 5 mM BME, 1 mM PMSF, and 0.2 mg/mL Lysozyme]. Resuspended cells were incubated on ice for 20 minutes, lysed by sonication, and then cleared by centrifugation at 40,000 x g for 30 minutes at 4°C. Clarified lysate was bound to 0.5 mL of HisPur Ni-NTA Resin (Thermo Scientific). The resin was then washed with 100 mL of buffer A and bound proteins were eluted in Buffer B [20 mM HEPES, pH 7.3, 150 mM NaCl, 2 mM MgCl2, 300 mM Imidazole, 10% Glycerol supplemented with 5 mM BME]. The purity of each fraction was determined by SDS-PAGE analysis followed by Coomassie staining. Fractions containing IFIT2 protein were pooled, supplemented with 0.4 mg mL-1 TEV protease, and dialyzed overnight at 4°C in buffer C [20 mM HEPES, pH 7.3, 150 mM NaCl, 2 mM MgCl2, 10% Glycerol supplemented with 1 mM BME]. The sample was then passed through 0.5 mL of Ni-NTA resin to remove TEV protease, hexahistidine peptide, and uncut IFIT2. The resulting flow through fraction was analyzed for purity by SDS-PAGE analysis followed by Coomassie staining. The sample was then concentrated, snap frozen, and stored at −80°C until use.

### DRaCALA Assay

For DRaCALA assays comparing binding of rIFIT2 and BSA to the 3’UTR, the 3’ 200 nucleotides of VEEV TC83 was amplified using primers listed in **Table S1**, and using the infectious clone cDNA for TC83. For DRaCALA assays comparing binding of rIFIT2 to viral and synthetic RNAs with different AU content, 101-106 nucleotide fragments were amplified using the primers listed in Table S1 using the TC83 infectious clone as template. Forward primers contained the sequence for the bacteriophage class III T7 promoter sequences. PCR products were purified and *in vitro* transcribed using T7 HiScribe kit (NEB). Transcribed RNA products were run on a 5% acrylamide TBE-urea gel and visualized with 0.02% methylene blue in 1x TBE. RNAs were excised from gels and transferred into a 0.6ml microfuge tube with a hole in the bottom (made by a sterile 22.5G needle). The 0.6ml tube was placed in a 2ml microfuge tube and centrifuged at 10,000xg for 5 min to crush gel pieces. RNA was eluted by adding 3x volume excess of RNA elution buffer [10mM Tris-HCL (pH7.5), 1mM EDTA, 300mM NaCl, 0.1% SDS]. Samples were then frozen at −80°C for 15 min then heated to 95°C for 5 min and placed on a rotator at room temperature overnight. Gel fragments were removed by centrifuging samples though a cellulose acetate filter (Thermo Scientific) and at 10,000 x g for 3 min at room temperature. RNA was then purified by acid phenol:chloroform extraction. *In vitro* transcribed RNA (5’-ppp) was dephosphorylated and 5’ end labeled with P^32^ as follows: 25 pmol of RNA was dephosphorylated using 50U of Antarctic Phosphatase (NEB) according to the manufacturer’s protocol. Dephosphorylated RNA was then 5’ end-labeled with 20U T4 PNK (NEB) and 30μCi [g32-P]ATP (Perkin-Elmer). Reactions were incubated for 1.5 hrs at 37°C. Unincorporated nucleotides were removed using Micro Bio-Spin 6 columns (Bio-rad) according to the manufacturers protocol, and samples then purified using Mag-Bind Total Pure NGS beads (Omega Bio-Tek). Prior to performing binding assays the RNA was folded by heating at 95°C for 2 min and slow cooled at room temperature. Binding reactions were performed as follows: 100 fmol of RNA was combined with serial 2-fold dilutions of rIFIT2 and incubated at 37°C for 15 min in binding buffer [50mM HEPES, 100mM KCl] supplemented with 5mM MgCl2, 5% glycerol, 500ng of yeast tRNA and 1mM DTT. 1 μl of each reaction was spotted in duplicate onto a 0.45μM nitrocellulose membrane (Amersham) and allowed to dry at room temperature for 20 minutes. RNA was then visualized by exposing the nitrocellulose to a phosphor screen and the resulting signal was measured on a Sapphire imager (Azure Biosystems).

## RESULTS

### A shRNA screen identifies IFIT2 as a restriction factor for Venezuelan equine encephalitis virus

Previously we identified IFIT1 as a restriction factor for replication of VEEV [9]. To identify other ISGs that restrict VEEV replication, we used a shRNA based approach to knock down expression of approximately 250 IFN- and virus-induced genes (Table S3; Table S4). HeLa cells were transduced with lentivirus expressing a bicistronic reporter consisting of GFP and shRNA targeting individual host genes, or a non-silencing control shRNA (shNSC). 48 hours post-transduction, cells were stimulated with human IFN-β to activate ISG expression, then infected with VEEV TC83 (MOI 1). At 24 hours post-infection (hpi) cells were fixed and analyzed by flow cytometry to quantify the number of VEEV positive cells, as determined by staining for VEEV E2 protein. Knock down of VEEV restriction factors was anticipated to lead to an increase in the number of VEEV E2-positive cells of the transduced population (shRNA-expressing; GFP positive) which are positive for. As expected, we observed that knock down of genes involved in antiviral responses to alphavirus infection (PKR; [46–48]) and IFN signaling (IRF1, IRF9, STAT2; [49–51]) led to a significant increase in the number of virus positive cells relative to a non-silencing control shRNA (shNSC). Due to the high experimental variability observed with lentivirus transduction, we chose a subset of genes for additional validation using a secondary screen.

These targets were chosen based on several criteria, including those with z-scores >2-fold, genes for which several individual shRNA sequences were detected as positive hits, genes encoding proteins predicted to be involved in RNA interactions, and targets which have not previously been explored in the context of alphavirus replication. In total we selected 148 unique shRNAs targeting 53 genes to perform the validation screen. As a positive control, we also included a shRNA against STAT2. The validation screen was performed as described for the primary screen, and fold increase in VEEV E2-positive cells (Q2) of the transduced population (Q2+Q3) relative to shNSC was calculated (Figure 1A and B; Table S5). Of the 53 genes, knock down of 24 of these resulted in a >3-fold increase in VEEV positive cells. Knock down of 49 of these genes resulted in a >2-fold increase in VEEV positive cells, however only 32 of these genes exhibited a >2-fold increase for multiple unique shRNAs (Figure 1A). Notably, we identified several genes and pathways which have previously been implicated in restricting the replication of other alphaviruses, including SNX5 [52] and the ISG15 pathway (HERC6, UBE2L6; [53–55]). We also identified several IFIT proteins (IFIT1 [9], IFIT2, and IFIT3) as restriction factors for VEEV.

**Figure 1.**
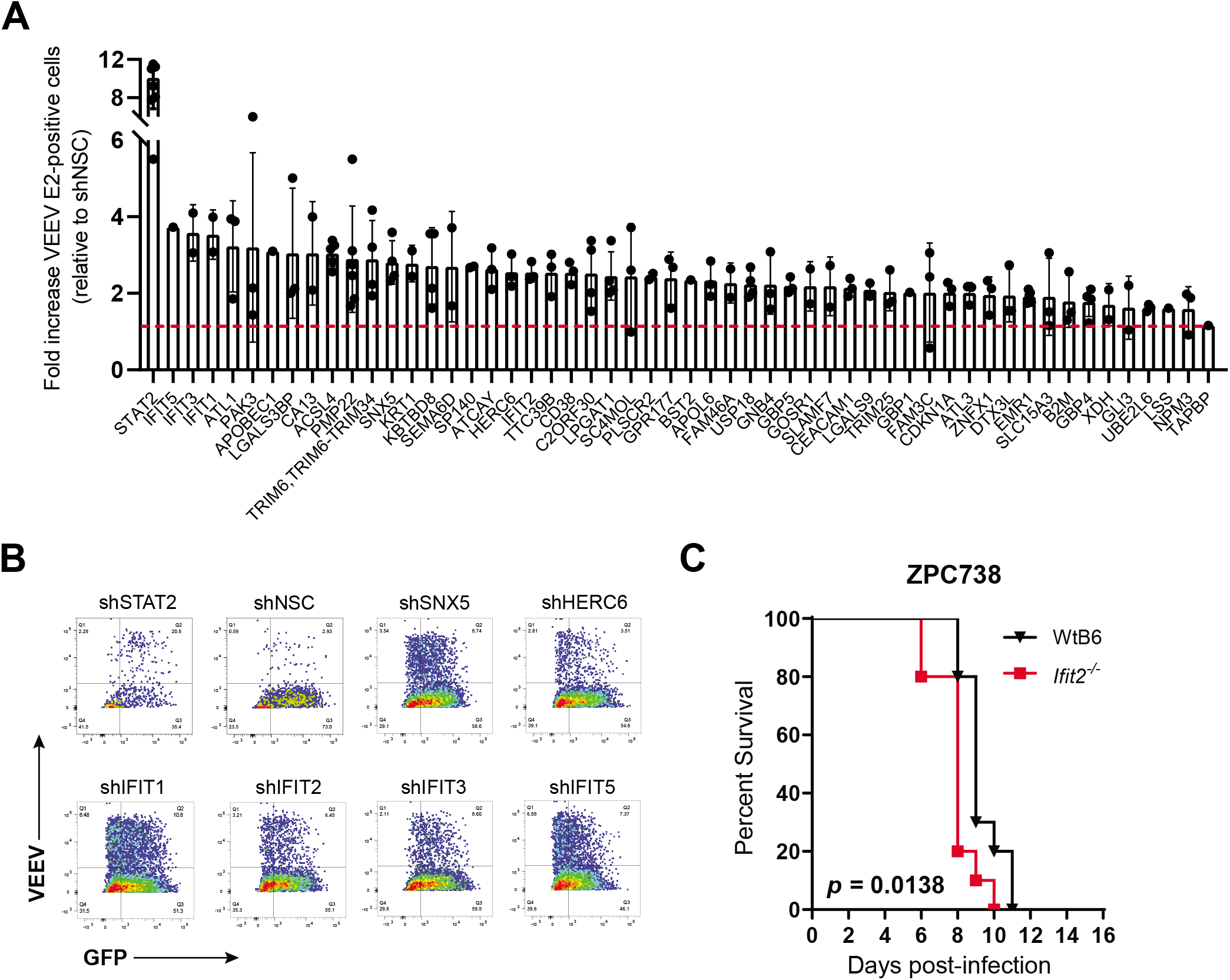
IFIT2 inhibits VEEV replication *in vitro* and pathogenesis *in vivo*. HeLa cells were transduced with lentiviruses encoding gene specific or non-silencing control shRNAs. Cells were treated with 10 IU/mL of IFN-β for 6 hours, infected with VEEV TC83 at a MOI of 1, and fixed and immunolabled for VEEV E2 and analyzed by flow cytometry 24 hours post-infection (hpi). Lentivirus (shRNA) positive cells are indicated by GFP, and E2-labled cells were counterlabeled with Alexaflour 647 secondary antibody. Summary of shRNA validation screen (53 genes) results expressed as the fold increase in VEEV positive cells relative to the shNSC control **(A)**. Each individual point represents a unique shRNA for the indicated gene. Mean fold change of all shRNAs against each target is represented by bars. **(B)** Representative flow plots of positive hits of known alphavirus restriction factors and IFN regulatory molecules detected in the validation screen. **(C)** 16-week-old WT (n=10) and *Ifit2^-/-^* (n=10) mice were infected with 10^2^ ffu of VEEV ZPC738 and monitored for survival (P = 0.0138). Statistical significance was calculated using Log-rank test.

We have previously identified IFIT1 as a restriction factor for VEEV and other alphaviruses [9]. As IFIT2 has previously been shown to inhibit replication of several other virus families and protect from lethal neuropathogenesis *in vivo* [14–19], and to be upregulated in the brains of mice following neurovirulent SINV infection [56], we decided to further explore the role of IFIT2 in restriction of VEEV replication. To validate our *in vitro* findings in an *in vivo* model of pathogenesis, we infected wild type (WT) and *Ifit2* knockout (*Ifit2^-/-^*) mice with VEEV ZPC738 and monitored animals for survival (Figure 1C). We observed a modest but significant decrease in the average survival time of *Ifit2^-/-^* mice infected with ZPC738 relative to WT mice (p = 0.0138), indicating that in addition to inhibition of replication *in vitro*, IFIT2 plays some role in VEEV restriction *in vivo*.

### The VEEV 3’UTR is a target of IFIT2

Like IFIT1, IFIT2 also possesses RNA binding activity [13], and has been proposed to exert its antiviral activity via a mechanism dependent on this property. Studies from our lab and others have shown that IFIT1 binds and restricts RNAs containing a 5’ 7-methylguanisine cap (m7G; cap 0), but not 2’-O-methylated cap structures (cap 1) [9, 11, 12, 57, 58]. In contrast, the RNA targets of IFIT2 are poorly defined. Previous studies have demonstrated binding specificity of human IFIT2 for poly (AU) RNA as well as RNAs containing AU-rich elements (AREs), but not polyA, polyU, or GC-rich RNA [13]. However, whether any AU-rich sequence is sufficient for recognition and binding by IFIT2, or whether other RNA motifs or surrounding sequences influence binding specificity is unknown. Analogous to cellular mRNAs, many +ssRNA viruses also contain AU-rich regulatory elements in the 3’ end of their genomes. As such we speculated that the alphavirus 3’ terminus which contains AU-rich sequences is a likely target of IFIT2. To explore this hypothesis, we first performed a sliding window analysis of the VEEV TC83 genome to identify regions with AU-rich sequences (Figure 2A). We used a Z-score cutoff of 2.32 (representing the 99^th^ percentile) to identify regions containing significantly high AU content (average = 50.2%) and observed multiple regions in the genome with high AU content, including several genes in open reading frame 1 (ORF1) as well as the 3’ UTR. As we predicted, the AU content of the 3’UTR, particularly the very 3’ end, was highest amongst all the regions identified (Z-score 2.5-4.2; Figure 2B). This is consistent with previous studies demonstrating IFIT2 binding specificity for AU-rich sequences [13] and proposed interactions with AU-rich elements (AREs) in cellular RNAs [23].

**Figure 2.**
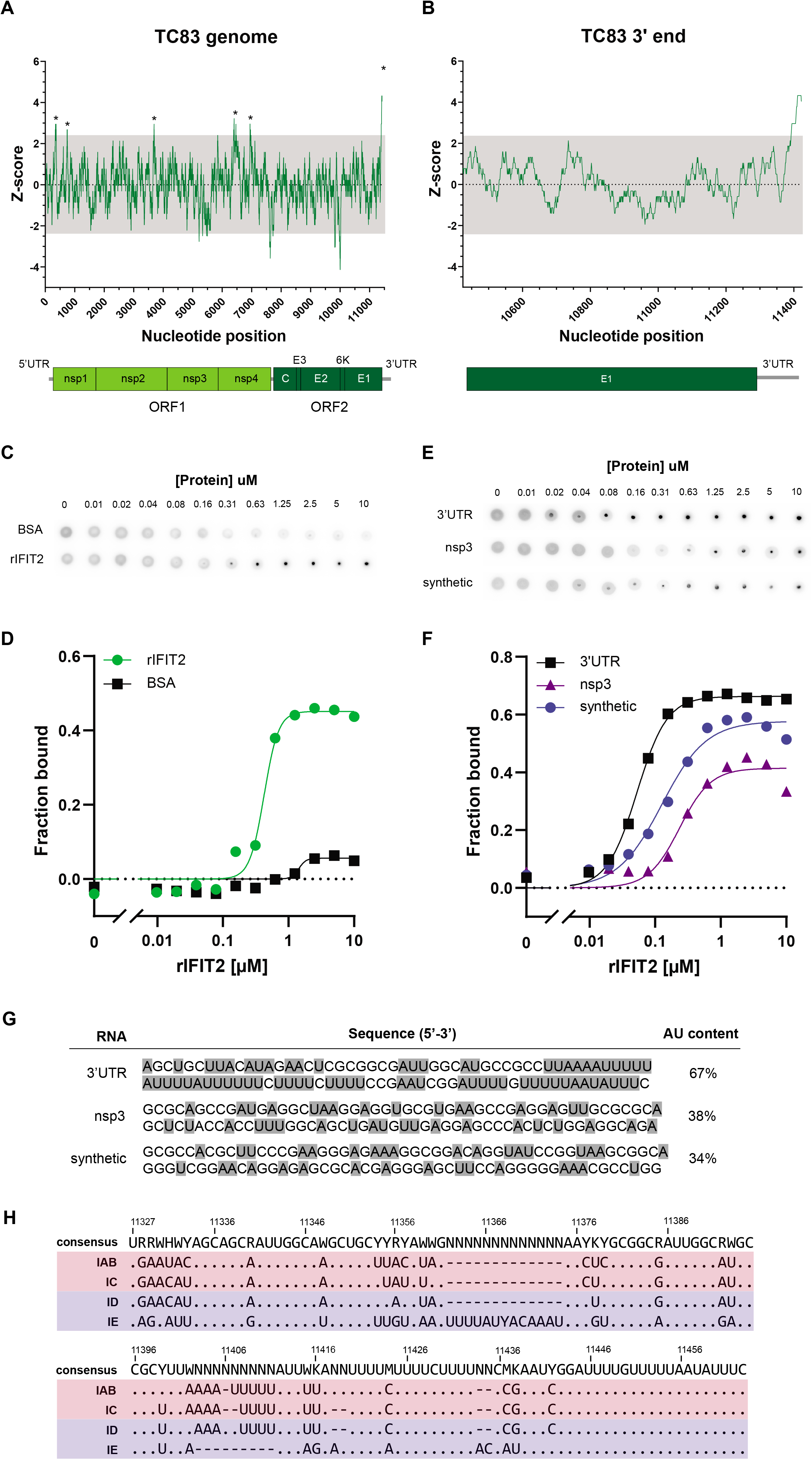
IFIT2 binds the VEEV 3’UTR. The VEEV TC83 genome nucleotide content was analyzed using sliding window analysis (window = 50 nucleotide, step = 1 nucleotide), and the Z-score of the AU content graphed **(A)**. A Z-score cutoff of 2.32635 (representing the 99^th^ percentile) was used to determine significance, and regions with Z-scores > 2.32635 are indicated by asterisk (*). Z-scores below this cutoff are shaded in grey. A schematic of the VEEV genome is depicted below. Gene boundaries correspond with the approximate nucleotide number represented in the Z-score analysis. 5’ and 3’UTRs are indicated by grey lines, and open reading frames and poly A tail are annotated. **(B)** Z-score analysis of the AU-content of the 3’ end of the VEEV TC83 genome comprising part of E1 and the full length 3’UTR, with the corresponding genomic region depicted below. **(C, D)** Determination of IFIT2 binding specificity using DRaCALA. P32 5’ end-labeled RNA corresponding to the 3’ terminal 200 nucleotide was incubated with serial 2-fold dilutions of rIFIT2 (green) or bovine serum albumin (BSA; black) for 15 min at 37°C, then spotted onto nitrocellulose and visualized after exposure to phosphor screen. A representative blot of a single replicate from one experiment is shown in **(C)**. The fraction of bound RNA was calculated and graphed in relation to protein concentration **(D)**. **(E, F)** Determination of IFIT2 binding specificity to different RNAs using DRaCALA. Binding of rIFIT2 to 100bp RNAs corresponding to the VEEV 3’UTR (black), VEEV nsp3 (purple), and a synthetic RNA (blue) was performed as described for (C) and (D). A representative blot of a single replicate from one experiment is shown in **(E)**. The fraction of bound RNA was calculated and graphed in relation to protein concentration **(F)**. (G) Sequence of 3’, nsp3, and synthetic RNAs assayed in E and F. AU content (%) for each RNA is indicated and A/U nucleotides highlighted in grey. **(H)** Sequence alignment of VEEV 3’UTR consensus sequences from IAB, IC, ID, and IE subtypes (see also **Supplementary Figure 1**). Full-length and partial VEEV genomes were aligned using Geneious Prime (MAFFT), and the 3’UTR region of the alignment extracted (including the ORF2 stop codon). Consensus sequences were generated from individual IAB, IC, ID, and IE alignments, and an alignment of these consensus sequences was performed. The ncbi accession number for each subtype analysis is listed in **Supplementary Figure 1**, and strains were grouped by subtype as indicated on the left of the alignment. Consensus sequence is shown at the top of the alignment in IUPAC-IUB notation (R=purine, Y=pyrimidine, K=keto (G/U), W=weak interaction (2-hydrogen bonds; A/U), H=not G). Identical sequences are represented as periods (.) and sequence gaps are represented as dashes (-).

To determine whether IFIT2 indeed binds the VEEV 3’UTR, we generated recombinant mouse IFIT2 (rIFIT2) and performed **<u>D</u>**ifferential **<u>Ra</u>**dial **<u>C</u>**apillary **<u>A</u>**ction of **<u>L</u>**igand **<u>A</u>**ssay (DRaCALA) [59] to measure interaction of IFIT2 with VEEV TC83 3’UTR RNA (Figure 2C). Analogous to filter binding assays, DRaCALA relies on the propensity of nitrocellulose to bind proteins but not small molecules such as RNA. When protein-ligand mixtures are applied to dry nitrocellulose membranes, protein and protein-bound ligands are immobilized at the site of application, whereas unbound ligand diffuses freely through the membrane via capillary action. We *in vitro* transcribed and P^32^ end-labeled RNA corresponding to the last 200 nucleotides of the TC83 genome that constitutes the 3’UTR and 79 nucleotides of upstream sequence and incubated radiolabeled RNA with serial dilutions of rIFIT2 or bovine serum albumin (BSA) as a negative control. Protein-RNA mixtures were then applied to nitrocellulose and the fraction of bound RNA quantified (Figure 2D) [59]. We chose to examine ligand binding using an RNA consisting of additional sequence upstream of the 3’UTR as the nucleotides at the start of the 3’UTR are predicted by RNAfold [60] to participate in base-pairing and therefore would be disrupted by examining binding to the 3’UTR sequence alone. In these assays we observed significantly higher binding of rIFIT2 to TC83 3’ RNA (K_d_ = 0.424 ± 0.051) as compared to BSA which exhibited minimal binding to RNA (K_d_ = 1.433 ± 0.028). Notably, the affinity of rIFIT2 for the VEEV 3’UTR is approximately 10-fold lower than that which we previously observed for IFIT1 and the VEEV 5’UTR [9], suggesting significant differences in affinity of these proteins for their respective RNAs which likely impacts their biological activities.

To validate whether the AU content of the 3’ RNA is important for determining specificity of IFIT2 binding to RNA, we performed additional DRaCALA assays comparing binding of rIFIT2 to 3’ RNA, and two additional RNAs with lower AU content (Figure 2E-G). The first of these corresponds to a region in nsp3 (38% AU), and second a synthetic RNA derived from the plasmid (non-viral sequence) encoding the TC83 infectious clone (34% AU). In these experiments, we synthesized, labeled, and purified ~100 nucleotide long RNAs as described above. Shorter RNAs were used for this experiment, as longer RNAs tended to have a higher AU content that were not significantly different from the 200 bp RNA fragment encoding the VEEV 3’ (Figure 2C and D). In these experiments we observed significantly higher binding of rIFIT2 to TC83 3’ RNA (K_d_ = 0.053 ± 0.030) as compared to either nsp3 RNA (K_d_ = 0.236 ± 0.069), or synthetic RNA (K_d_ = 0.126 ± 0.050). We observed increased binding of rIFIT2 to the 100 bp 3’ RNA as compared to the 200 bp RNA, as observed by an 8-fold decrease in the measured K_d_ of the shorter RNA. This may be explained by the slightly lower AU content of the 200 bp RNA (62%) as compared to the 100 bp RNA (66%), although we cannot rule out that additional structures in the 200 bp RNA may modulate IFIT2 binding. Notably, we observed decreased binding of rIFIT2 to both the nsp3 and synthetic RNAs, which exhibited a 4.5-fold and 2.4-fold increase in K_d_ relative to 3’ RNA respectively (compare blue and purple lines in Figure 2F). Interestingly, we observed that the binding affinity of IFIT2 for the nsp3 RNA was lowest, despite the fact that the AU content of this RNA is slighter higher than the synthetic RNA. At present it is unknown what the exact determinants of IFIT2-RNA binding are (e.g. AU track length, RNA structure) and we are currently exploring this further. Overall, our binding data shows that IFIT2 preferentially binds AU-rich RNA, and suggests that other factors such as exact primary sequence and secondary structure may also modulate the specificity of binding.

### Changes in the VEEV 3’UTR sequence alter restriction of VEEV replication by IFIT2 *in vitro*

To determine whether 3’UTR sequences are conserved across different VEEV species, we compared full-length and partial 3’UTR sequences from 136 strains representing four major VEEV subtypes (IAB, IC, ID, and IE) (Figure 2E; Figure S1). As seen with TC83, we observed several stretches of polyAU sequence throughout the 3’UTR, as is also seen in many +ssRNA viruses. 3’UTR sequences from IE subtypes were considerably different from IAB, IC, and ID sequences, consistent with the fact that IE subtypes are more evolutionarily divergent from these other subtypes [61, 62]. However, notably we observed that the 3’UTR sequences of different VEEV strains contained multiple nucleotide variations. Although we did not identify a single mutation or groups of mutations that were exclusively found in viruses belonging to a single subtype (except for the more divergent IE viruses), (Figure 2E, bottom sequence), several mutations appeared more frequently in viruses belonging to a given subtype. Therefore, we speculated that 3’UTR nucleotide variations present in individual subtypes may lead to alteration in the biological functions of viral 3’UTR RNA, including susceptibility to IFIT2 binding and inhibition of replication.

In order to test our hypothesis, we introduced select mutations from IC, ID, and IE subtypes into the 3’UTR of the TC83 infectious clone (Figure 3A; Table S6). VEEV subtypes exhibit genetic diversity and can be divided into different lineages, with epizootic strains being further classified into clades [1]. We chose several different 3’UTR sequences in order to capture the sequence variation observed, particularly within the ID subtype which encompasses several major lineages and several epizootic clades (Figure S1; Table S6). We compared replication kinetics of 3’UTR chimeras in primary murine embryonic fibroblasts (MEF) derived from WT and *Ifit2^-/-^* mice (Figure 3B-E). MEF were mock or IFN-β pre-treated for 12 hours to induce IFIT2 expression and infected at a MOI of 0.01. Cell culture supernatant was harvested at 1, 6, 12, 24, 36, and 48 hours post-infection (hpi) and infectious virus quantified using focus forming assay (FFA). When we compared replication of WT and mutant viruses in IFN-β pre-treated samples, we observed that replication of TC83 and TC83/IC-3’UTR viruses (Figure 3B and C) was consistently higher in *Ifit2^-/-^* cells as compared to WT (>1 log) though not statistically significantly (IAB WT vs KO, p=0.3553), although differences in replication of IC in WT vs KO cells was approaching significance (0.0845). In contrast, we observed identical replication kinetics of TC83/ID-3’UTR and TC83/IE-3’UTR mutant viruses in IFN-stimulated WT and *Ifit2^-/-^* cells (Figure 3D and E), suggesting that these viruses are resistant to the antiviral activities of IFIT2. When comparing replication of all viruses in KO cells pre-treated with IFN, no biological or statistical significance was observed (Figure 3G; compare red shaded lines). When we compared replication of all four viruses in WT IFN pre-treated cells (Figure 3G; compare black and grey shaded lines), we observed significant differences between epizootic and enzootic 3’UTR chimeras (IAB vs ID, p=0.0061; IAB vs IE, p=0.0019; IC vs IE, p=0.0184) except for IC vs ID, although this difference was approaching statistical significance (IC vs ID, p=0.0577). Notably, replication kinetics and viral titers in mock-treated samples were near identical for all viruses (compare solid lines in Figure 3B-E), indicating that: i) this phenotype is IFN-dependent and; ii) that the observed IFIT2 phenotype is not due to replication defects or advantages caused by introduction of these mutations. Therefore, we conclude that the differential replication observed between IAB and IC versus ID and IE mutants can be attributed to the specific activities of IFIT2. Of note, TC83/IC-3’UTR and TC83/ID-3’UTR mutant viruses differ only by a single nucleotide (U-to-C mutation at nucleotide 11366; compare Figure 3C and D), suggesting that the differences in IFIT2-mediated restriction could potentially be driven by changes in 3’UTR RNA structure and not primary sequence. This is supported by findings that the primary sequence of the IFIT2 ligand (AU-rich RNA) is somewhat broadly defined. Although higher overall AU content clearly coincides with greater IFIT2 binding (Figure 2F and G), IFIT2 is still able to bind RNAs with variable AU content and length of AU-rich tracts. As such we hypothesize that it would be unlikely that a single nucleotide change would significantly alter the linear IFIT2 recognition motif sufficiently to result in a significant change in IFIT2 binding or IFIT2-dependent viral replication, particularly given that the position of the nucleotide change in IC versus ID does not disrupt the AU-rich tracts that lie downstream of this nucleotide (Figure 3A). This is further supported by our previous observations that single nucleotide mutations in the alphavirus 5’UTR is sufficient to alter RNA secondary structure in a manner which prevents binding of RNA by IFIT1 [9].

**Figure 3.**
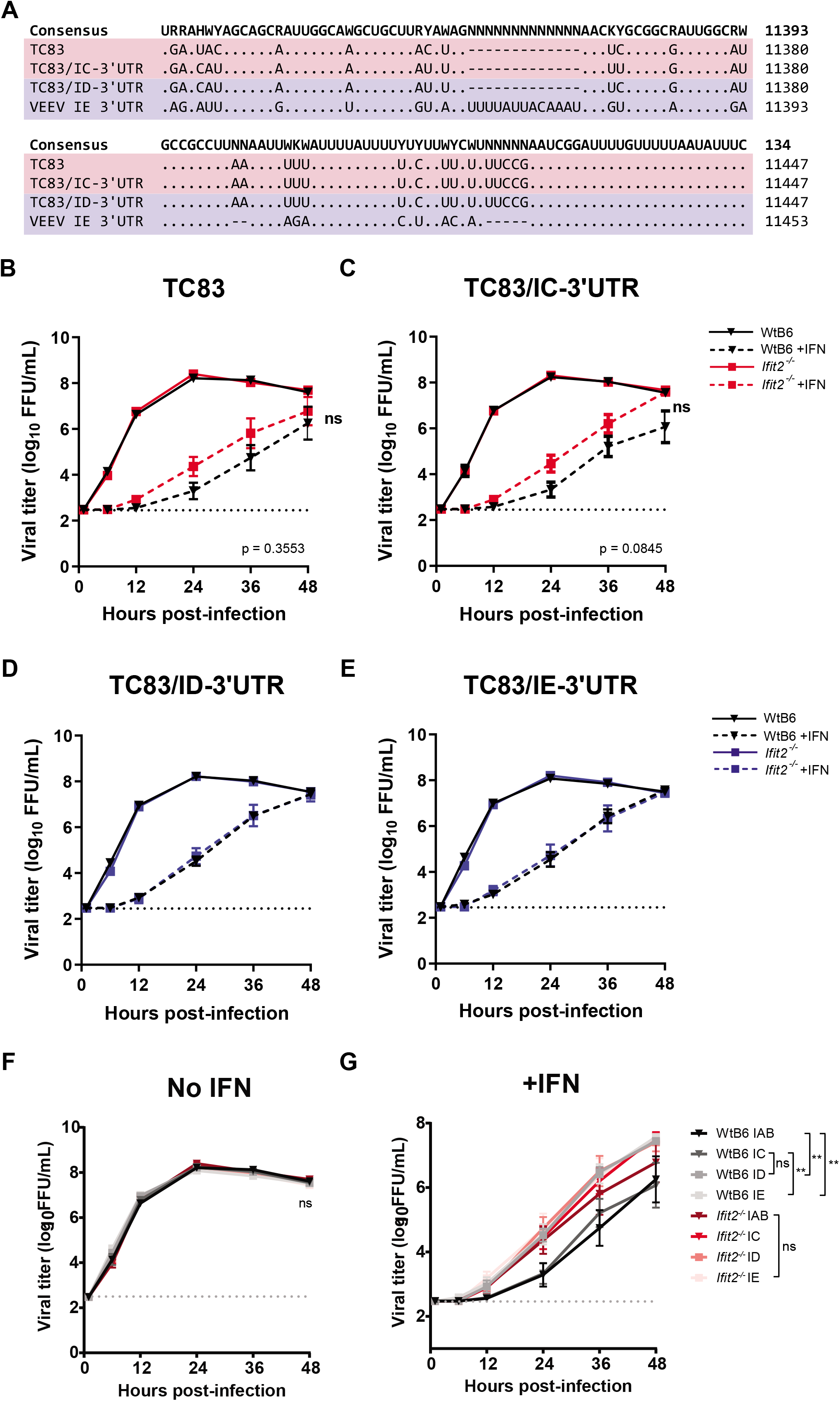
IFIT2 differentially inhibits replication of VEEV mutants encoding distinct 3’UTR sequences. **(A)** Sequence alignment of 3’UTR sequences from TC83, IC (TC83/IC-3’UTR), ID (TC83/ID-3’UTR), and IE (TC83/IE-3’UTR) 3’UTR chimeras. Viruses encoding epizootic 3’UTRs (IAB and IC) are shaded in red, and viruses encoding enzootic 3’UTRs (ID and IE) are shaded in blue. Consensus sequence is shown at the top of the alignment in IUPAC-IUB notation (R=purine, Y=pyrimidine, K=keto (G/U), W=weak interaction (2-hydrogen bonds; A/U), H=not G). Nucleotide numbering is indicated on the right side. Identical sequences are represented as periods (.) and sequence gaps are represented as dashes (-). **(B-G)** Growth kinetics of VEEV 3’UTR mutants in WT and *Ifit2^-/-^* primary MEF. WT (black) and *Ifit2^-/-^* (red or blue) cells were mock (solid lines) of IFN-β pre-treated (dotted lines) for 12 hours, then infected with indicated viruses at a MOI of 0.01. Cell culture supernatant was serially harvested at 1, 6, 12, 24, 36, and 48 hpi and infectious virus titered using focus forming assay (FFA). Red lines indicate mutant viruses encoding epizootic 3’UTR sequences, and blue indicate mutant viruses encoding enzootic 3’UTR sequences. Infections with each virus were performed simultaneously and displayed individually for ease of viewing. Each experiment was performed in triplicate three times independently and the mean and SEM graphed. Statistical analysis was performed by calculating the area under the curve (AUC) for each replicate and experiment, and AUC values from WT and KO cells analyzed by unpaired t-test. **(F, G)** Data from B-E plotted by treatment (no IFN, +IFN). Statistical significance comparing all viruses in WT cells +IFN and KO cells +IFN was calculated on the AUC by one-way ANOVA with multiple comparisons and is indicated on the right. **, *P* ≤ 0.01; ns, not significant.

### IFIT2 modulates VEEV replication in a cell type-dependent manner

We wanted to further explore the role of IFIT2 inhibition of VEEV replication in more physiologically relevant cell types. Myeloid cells including macrophages are early targets of VEEV infection *in vivo* [44, 63, 64]. In contrast to non-myeloid cells (including MEF) which fail to secrete IFN-α/β following alphavirus infection due to viral-mediate transcriptional and translational inhibition [65, 66], macrophages are significant producers of type-I IFN *in vitro* and *in vivo* [51]. To determine the role of IFIT2 in inhibition of VEEV replication in macrophages, we generated *Ifit2* CRISPR Raw264.7 macrophages using a doxycycline-inducible CRISPR/Cas9 all-in-one expression vector encoding Cas9 and individual *Ifit2*-specific gRNAs (g13, g25, g38). We verified *Ifit2* gene KO of bulk cells by western blotting (Figure 4A), and then compared replication kinetics of 3’UTR mutants in WT (Raw264.7-Empty) and *Ifit2^-/-^* CRISPR KO (Raw264.7-*Ifit2*.g13) cells (Figure 4B-E). As macrophages basally express IFIT2 (Figure S2) and secrete IFN-α/β following VEEV infection [51], cells were not pre-treated with IFN-β prior to infection to induce ISG expression. Cells were infected at a MOI of 0.01, and cell culture supernatant was harvested at 1, 6, 12, 24, 36, and 48 hpi and infectious virus quantified by FFA.

**Figure 4.**
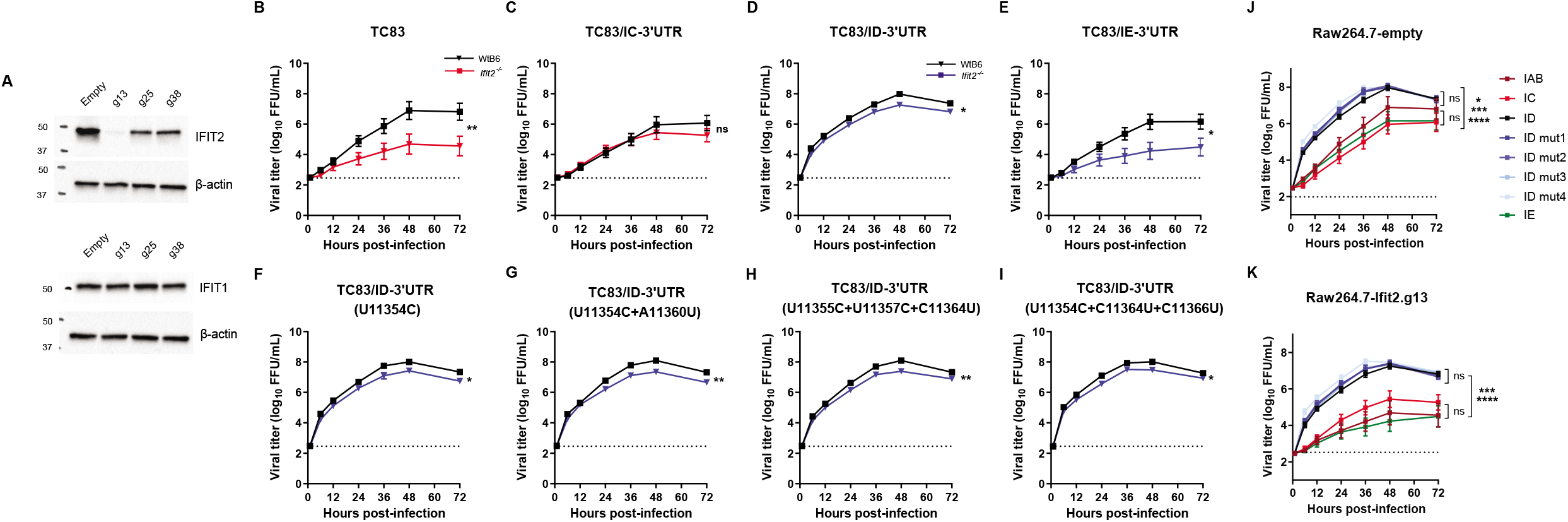
IFIT2 differentially inhibits replication of VEEV 3’UTR mutants in a cell type-dependent manner. Validation of *Ifit2* CRISPR KO macrophages (Raw264.7); **(A)** generated using CRISPR. Cell lysates were electrophoresed, blotted onto nitrocellulose, and probed using monoclonal IFIT2-specific antibodies, as well as IFIT1 antibodies to determine specificity of gene KO for IFIT2. Growth kinetics of 3’UTR mutants in WT (empty; black) and Ifit2 KO (g13; red/blue) Raw264.7 **(B-I)**. Red lines indicate mutant viruses encoding epizootic 3’UTR sequences, and blue lines indicate mutant viruses encoding enzootic 3’UTR sequences. Each experiment was performed in triplicate three times independently and the mean and SEM graphed. Statistical analysis was performed by calculating the area under the curve (AUC) for each replicate and experiment, and AUC values from WT and KO cells analyzed by unpaired t-test. **(J,K)** Data from B-I plotted by cell line (Raw264.7-empty, Raw264.7-g13). Statistical significance comparing all viruses in WT or KO cells was calculated on the AUC by one-way ANOVA with multiple comparisons. Comparison of all non-ID viruses to all ID viruses was significant. No significant difference was observed between any of the ID viruses. *, *P* ≤ 0.05; **, *P* ≤ 0.01; ***, *P* ≤ 0.001; ****, *P* ≤ 0.0001; ns, not significant.

In stark contrast to our observations of virus replication in primary MEF, where *Ifit2* KO resulted in increased virus replication (Figure 3), we observed a decrease in VEEV replication in *Ifit2^-/-^* versus WT macrophages (Figure 4B-E), suggesting that IFIT2 functions in a cell type-dependent manner to modulate VEEV replication. However, similar to MEF, we observed that changes in the VEEV 3’UTR sequence also impact IFIT2-dependency of virus replication in macrophages. In WT cells we observed similar viral titers over the course of the infection with TC83, TC83/IC-3’UTR, and TC83/IE-3’UTR viruses (Figure 4B, C, E; black line). However, in contrast to TC83/IC-3’UTR which replicated similarly in both WT and *Ifit2* KO cells, TC83 and TC83/IE-3’UTR mutants exhibited significantly less replication in KO cells relative to WT (*P* = 0.0283 and *P* = 0.0382 respectively; compare black and red lines in Figure 4B, C, E). We also observed a modest (2-5 fold) but significant decrease in replication of TC83/ID-3’UTR virus in *Ifit2* KO vs WT cells (*P* = 0.0103). Strikingly, we observed that TC83/ID-3’UTR, which differs from IC by only a single nucleotide, replicated to significantly higher titers than TC83/IC-3’UTR in both WT (20-100 fold increase; *P* < 0.0001) and *Ifit2* KO macrophages (20-80 fold increase; *P* = 0.004) (Figure 4D), suggesting that changes in the virus 3’UTR not only affect VEEV IFIT2-dependency, but also impact VEEV replication in an IFIT2-independent manner. This effect was also cell type-dependent, and possibly IFN-dependent, as we did not observe any difference in replication of IC versus ID mutants in WT mock-treated MEF or in IFIT2 deficient cells following IFN pre-treatment (Figure 3).

To determine whether other SNPs identified in subtype ID 3’UTR sequences from different VEEV lineages (Supplemental Figure 1) similarly affected virus replication in macrophages, we also constructed additional TC83/ID-3’UTR mutants (U11354C; U11355C+A11360U; U11355C+U11357C+C11364U; U11354C+C11364U+U11366U; Table S6) and compared replication of these viruses in WT and *Ifit2* KO Raw264.7 cells simultaneously (Figure 4F-I). Remarkably, all mutants exhibited near-identical replication to each other as well as the parent ID virus, suggesting that the macrophage-specific function of the 3’UTR is conserved amongst different ID subtypes, despite the presence of these nucleotide variations. Collectively, this data suggests that: i) IFIT2 modulates VEEV replication in a cell type-dependent manner; ii) changes in the VEEV 3’UTR alter the IFIT2-dependency of virus replication; iii) mutations in the virus 3’UTR (IC vs ID) also affects replication of VEEV independent of IFIT2, in a cell type manner; iv) while subtype ID viruses contain numerous SNPs, the 3’UTR-dependent replication phenotype of these viruses is conserved.

### IFIT2 modulates VEEV replication via a translation-independent mechanism

Several biological functions have been ascribed to IFIT2, including translational control, cancer development, and apoptosis. Multiple studies have implicated translational control to be the major mechanism by which IFIT2 controls viral replication, which can occur through sequestration of host translation initiation factors [67], direct binding of viral RNA [13, 37], or indirectly through interaction with IFIT1 [21, 68]. To determine whether the 3’UTR-dependent replication phenotypes observed in fibroblasts and macrophages could be explained by differences in translation, we compared translation of viral reporter RNAs (VEEV TC83/FLuc/rep; [9, 43]) encoding mutant 3’UTRs in WT and *Ifit2^-/-^* primary MEF and Raw264.7 cells (Figure 5). Translation reporters consist of the firefly luciferase gene (FLuc) fused in-frame downstream of a truncated VEEV nsP3 sequence flanked at either end by the VEEV 5’- and 3’UTRs. We chose to use reporter RNAs to assess translation as this allows us to uncouple effects of IFIT2 on viral RNA translation versus RNA replication and transcription. WT and *Ifit2^-/-^* cells were mock-or pre-treated with IFN-β for 12 hours, then electroporated with *in vitro* transcribed m7G-capped reporter RNA. Cell lysates were harvested at 30, 60, 120, and 240 minutes post electroporation and translation measured by quantifying FLuc activity and normalizing to total protein. As expected, IFN pre-treatment of MEF significantly inhibited translation of all reporter RNAs in both WT and *Ifit2^-/-^* cells (compare Figure 5A and B to Figure 5C and D), demonstrating the contribution of other ISGs (including IFIT1) to translation inhibition of viral RNA. However, we observed no significant difference in translation of reporter RNAs encoding WT or mutant 3’UTR sequences. Interestingly, translation of all reporter RNAs appeared to trend slightly higher or lower in WT and *Ifit2^-/-^* cells under conditions of mock- or IFN pre-treatment respectively, although this was not statistically significant.

**Figure 5.**
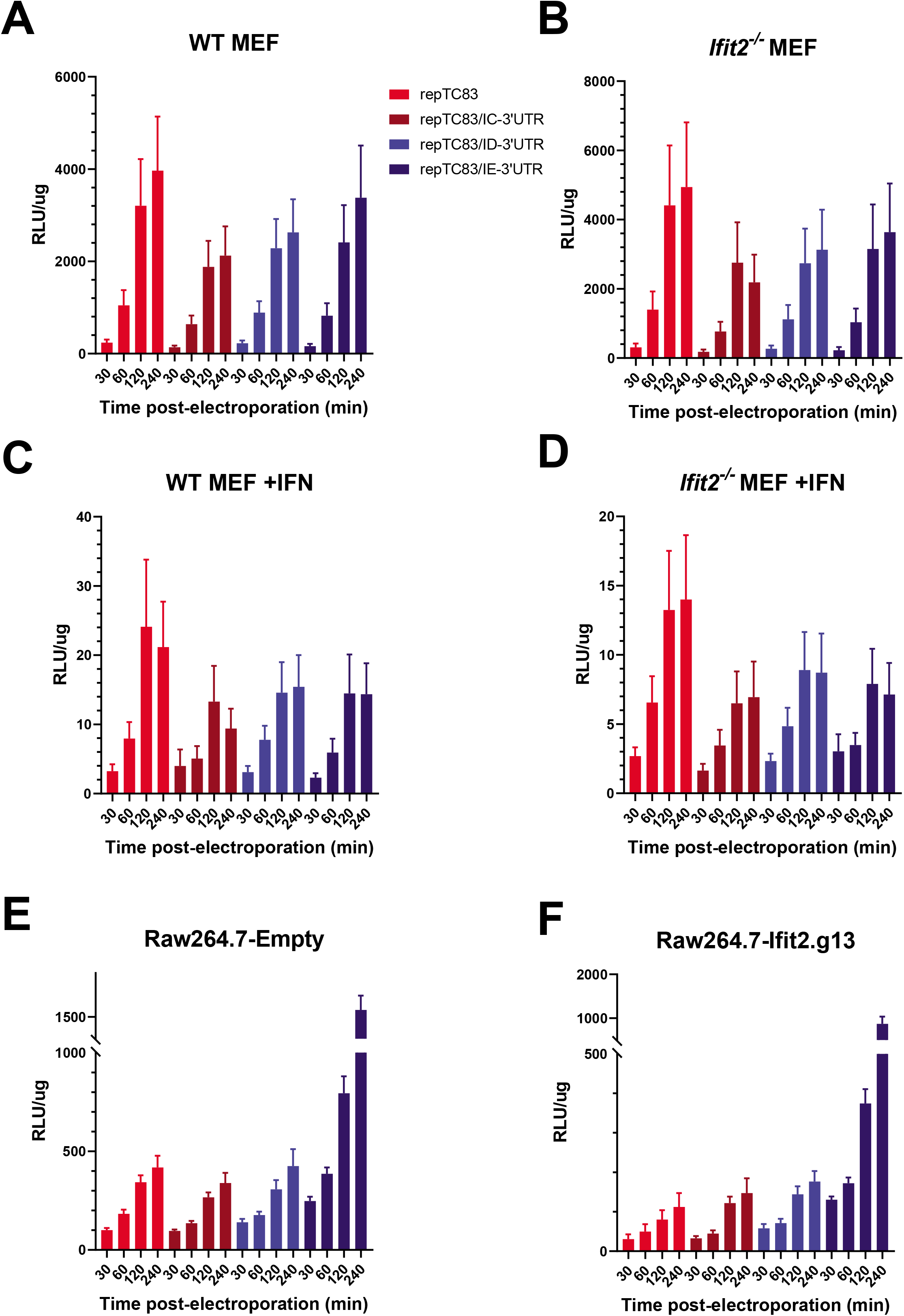
IFIT2-dependent inhibition of 3’UTR mutants is translation independent. WT and *Ifit2^-/-^* primary MEF were mock **(A, B)** or IFN-β pre-treated **(C, D)**, then electroporated with 2ug of viral reporter RNAs encoding TC83, IC, ID, or IE 3’UTR sequences. Cell lysates were harvested at 30, 60, 120, and 240 minutes post-electroporation, luciferase activity measured, and normalized to total protein. Measurements are given as relative light units (RLU)/ug of protein. Each data point represents three independent experiments. **(E and F)** WT (Empty) and *Ifit2^-/-^* CRISPR (g13) Raw264.7 cells were electroporated with 10μg viral reporter RNAs and lysates harvested and analyzed for luciferase activity as for MEF.

As we observed contrasting replication phenotypes in fibroblasts and macrophages, we also performed translation reporter assays in WT and *Ifit2* CRISPR KO Raw264.7 (Figure 5E and F), to determine whether these differences could be attributed to differential IFIT2-dependent translation in these cell types. Assays were performed as for MEF (described above), however unlike MEF, Raw264.7 cells were not pre-treated with IFN-β as these cells basally express IFIT2 (Figure S2). When we compared translation of reporter RNAs in these cells we observed a modest (~2-4 fold) but significant decrease in translation of all reporter RNAs in WT vs KO cells, consistent with regulation role for IFIT2 in global translation regulation. However, we observed no difference in translation between reporter RNAs in either WT or KO cells, with the exception of repTC83/IE-3’UTR, suggesting that 3’UTR-dependent differences in translation do not account for our observed replication phenotypes in either fibroblasts or macrophages. Interestingly, we observed up to a 5-fold (WT) or 7.7-fold (*Ifit2^-/-^*) increase in translation of repTC83/IE-3’UTR relative to other reporters, specifically in macrophages but not fibroblasts. Of note, we observed no difference in translation of either IC or ID in either cell type, despite the fact that the single nucleotide difference between these RNAs accounts for significant differences in replication in both cell lines (Figure 3 and 4). Collectively, our translation reporter data suggests that IFIT2 plays a modest role in global regulation of cellular translation in fibroblasts and macrophages, consistent with previous reports of the role of IFIT2 in translation inhibition and regulation [22, 67]. However, the mechanism by which IFIT2 inhibits (MEF) or promotes (Raw264.7) replication of 3’UTR mutants appears to be independent of viral RNA translation. Given that IFIT2 has been shown to bind to and regulate translation of cellular mRNAs [23, 37], it is possible that IFIT2-dependent translation inhibition of a subset of cellular transcripts could explain these observations. Further studies are necessary to elucidate the molecular mechanism by which IFIT2 regulates replication of VEEV and how the viral 3’UTR contributes to replication, innate immune evasion, and pathogenesis.

## DISCUSSION

In this study, we used a shRNA screening approach to identify VEEV restriction factors and identified IFIT2 as a novel viral restriction factor of VEEV replication and pathogenesis. We identified that IFIT2 specifically binds to the 3’UTR of the VEEV genome, which contains AU rich elements, a broadly defined motif abundant in both cellular and +ssRNA viral RNA. This is consistent with previous studies of human IFIT2 which has been shown to bind AU rich RNA sequences [13] as well as AREs found in mouse transcripts [23]. When we compared 3’UTR sequences from different strains of VEEV, we observed tremendous diversity in their sequences. We also noted that the 3’UTRs of viruses belonging to different subtypes (IAB, IC, ID, and IE) of VEEV were distinct, and speculated that these sequence differences may confer functional changes in 3’UTR RNA that affect replication and innate immune evasion. To test this, we compared replication of IAB, IC, ID, and IE 3’UTR mutants in WT and *Ifit2^-/-^* primary fibroblasts, as well as macrophages which are early targets of VEEV infection *in vivo* and important producers of type I IFN [51].

We made several striking observations in these cell lines. Firstly, we observed opposing IFIT2-dependent phenotypes, with an antiviral role in fibroblasts (Figure 3) and a proviral role in macrophages (Figure 4). Although the majority of viral pathogenesis studies to date have predominantly shown an antiviral role for IFIT2, a recent study by Tran et al suggests that influenza virus usurps IFIT2 to promote translation of viral mRNAs *in vitro* indicating a possible proviral role [37]. While our observations, along with others indicate cell-type specificity, our mouse infection studies with VEEV ZPC738 shows that IFIT2 predominantly plays an antiviral role during VEEV pathogenesis (Figure 1C). As TC83 is highly attenuated in C57BL/6J mice compared to VEEV ZPC738, such direct comparisons might not be possible at this time. TC83 infection causes only modest weight loss in WT C57BL/6J mice, and only minor or modest clinical manifestations, thus this model is not sufficiently robust to distinguish and dissect modest in vivo phenotypes. As such, we are generating 3’UTR chimeras on in other VEEV backgrounds (ZPC738) which will be used to investigate this hypothesis in following studies. We also cannot exclude the possibility that other VEEV sequences outside of the 3’UTR may contribute to IFIT2-dependent phenotypes. Nonetheless, our data shows that the biological function of IFIT2 during virus replication differs depending on the cell type, and likely the virus as well.

Secondly, we observed that changes in the VEEV 3’UTR had a significant effect on virus replication in an IFIT2-dependent manner. While the effect of coding sequence mutations has previously been studied in the context of epizootic VEEV emergence, the role of sequence variability in the 3’UTR has not been investigated. Despite observing opposing IFIT2 phenotypes in fibroblasts and macrophages, changes in the VEEV 3’UTR affected the dependency of VEEV replication on IFIT2 in both cell types. Interestingly, the effect of some 3’UTR mutations on IFIT2-dependent phenotypes differed between these cells. For example, in MEF we observed an IFIT2-dependent replication phenotype with TC83/IC-3’UTR (Figure 3C), but in macrophages we observed no significant difference in replication of this mutant in WT vs KO cells (compare with Figure 4C). These data may suggest that other cellular factors which are differentially expressed in these cell types also impacts IFIT2. Other IFIT proteins have been demonstrated to form hetero-oligomers to modulate their functions. Recent studies have shown that IFIT2 and IFIT3 interact with IFIT1 leading to enhanced cap binding activities of this protein [21, 68]. We propose other host factors which interact with IFIT2 alter the biological properties of IFIT2, and/or IFIT2 competes with other RNA-binding proteins for binding to the VEEV 3’UTR and the outcome of this competition is influenced by the underlying sequence and secondary structure of the viral RNA.

We were particularly intrigued by differences in replication of IC and ID 3’UTR mutant viruses. In MEF we observed that the ID mutant was resistant to IFIT2-mediated inhibition (compare Figure 3C and 3D), and in macrophages observed a significant increase in replication (~20-200 fold) relative to the IC mutant (compare Figure 4C and 4D). These mutants differ by only a single nucleotide and are derived from equine virulent (epizootic; IC) and equine avirulent (enzootic; ID) subtypes. Epizootic (IC) VEEV subtypes arise in nature via de novo mutagenesis of enzootic subtypes (ID) which continually circulate in a sylvatic transmission cycle. With the exception of a IE epizootic outbreak in Mexico in 1996 [69, 70], IC subtypes are thought to be the predominant epizootic subtype and emerge in nature via mutation of ID enzootic viruses [71–75]. Previous studies have shown that acquisition of mutations in the viral attachment protein (E2) are essential for conferring this epizootic phenotype, which is characterized by increased type I IFN resistance [7, 8, 72, 73], as well as high titer viremia and pathogenesis in equines which serve as amplification hosts during epizootic episodes [7]. While these mutations are critical for emergence of epizootic VEEV, comparative analysis of epizootic and enzootic sequences reveal that epizootic viruses also acquire other mutations elsewhere in the genome, including the 3’UTR (Figure S1). Significantly, a high proportion of these mutations are synonymous (unpublished data). While many of these mutations can likely be explained by divergent evolution due to geological constraints, we speculate that some mutations may confer biological properties (e.g. type I IFN resistance) that, although not essential, may aid in emergence of epizootic subtypes. Thus we speculate that acquisition of mutations that alter the function of underlying diversity in the sequences of 3’UTR may contribute to emergence of epizootic VEEV by allowing the virus to overcome the host innate immune response, or by altering virus replication properties in specific intrahost niches and cell types (e.g. macrophages). Our observations that the 3’UTRs of epizootic and enzootic viruses alter the replication properties and IFIT2-dependency of VEEV lends support to this hypothesis.

Multiple studies have demonstrated a role for IFIT2 in regulation of viral and cellular translation, either through sequestration of translation factors or direct binding to viral RNA. We observed a small decrease in translation of all reporter RNAs in *Ifit2* KO vs WT cells, consistent with a role for IFIT2 in global RNA translation. However, no significant differences were observed between mutant 3’UTR reporter RNAs in either WT or KO cells. Interestingly, we observed a significant increase in translation of IE 3’UTR reporter RNA relative to the other VEEV 3’UTRs, specifically in macrophages. These data would suggest that mutations in the IE 3’UTR enhances viral RNA translation specifically in macrophages. This again bolsters our hypothesis that cell-type specific host factors expressed in macrophages are critical for determining VEEV replication, however how this enhanced translation would impact replication and pathogenesis of IE subtypes *in vivo* is at present unclear. Nonetheless, these data show that our observed IFIT2-dependent replication phenotypes of 3’UTR mutants cannot be explained by differences in translation alone.

In addition to demonstrating that the VEEV 3’UTR modulates IFIT2-dependent activities, we also observed IFIT2-independent roles for the 3’UTR in VEEV replication and RNA translation. Moreover, these effects were cell type-dependent, suggesting that the 3’UTR plays a crucial role in regulating replication, and possibly immune evasion, specifically in macrophages. Recent studies have demonstrated that macrophages are important early targets of VEEV infection *in vivo* and upon initial infection are capable of secreting significant amounts of type-I IFN [51]. This is particularly relevant as early and robust production of IFN in the periphery has been shown to limit replication and dissemination to the central nervous system of other neurotropic alphaviruses [76]. Thus, we speculate that changes in the VEEV 3’UTR that alter replication and innate immune responses in macrophages may have profound impacts of viral pathogenesis *in vivo* and is a focus of our future studies.

In summary, we identified a role for IFIT2 in restricting VEEV replication and pathogenesis *in vitro in vivo*. We demonstrated that IFIT2 targets VEEV by binding to the 3’UTR and show that changes in the VEEV 3’UTR sequence modulate the ability of IFIT2 to inhibit VEEV replication in a cell type-dependent manner. In contrast to previous studies, our data suggests that IFIT2 affects replication of VEEV 3’UTR mutants via a mechanism independent of translation. We also demonstrated an IFIT2-independent role for the VEEV 3’UTR in replication in macrophages. These findings have broad implications for the role of the alphavirus 3’UTR in the replication and pathogenesis of VEEV, as well as for the emergence of epizootic VEEV subtypes.

## SUPPLEMENTARY FIGURES

**Figure S1. Consensus sequence alignment of full and partial VEEV 3’UTR sequences.** Full-length and partial VEEV genomes were aligned using Geneious Prime (MAFFT), and the 3’UTR region of the alignment extracted (including the ORF2 stop codon). The NCBI accession number for each virus is listed next to the corresponding sequence, and strains were grouped by subtype as indicated on the left of the alignment. Consensus sequence is shown at the top of the alignment. Identical sequences are represented as periods (.) and sequence gaps are represented as dashes (-).

**Figure S2. Basal and induced expression of IFIT2 in Raw264.7 macrophages.** Raw264.7 macrophages were mock treated or stimulated with 100ug of high molecular weight (HMW) poly I:C for 24 hours. Cells were then lysed in RIPA buffer and analyzed by western blotting using antigen-specific antibodies against IFIT2 or β-actin.

**Table S1.**
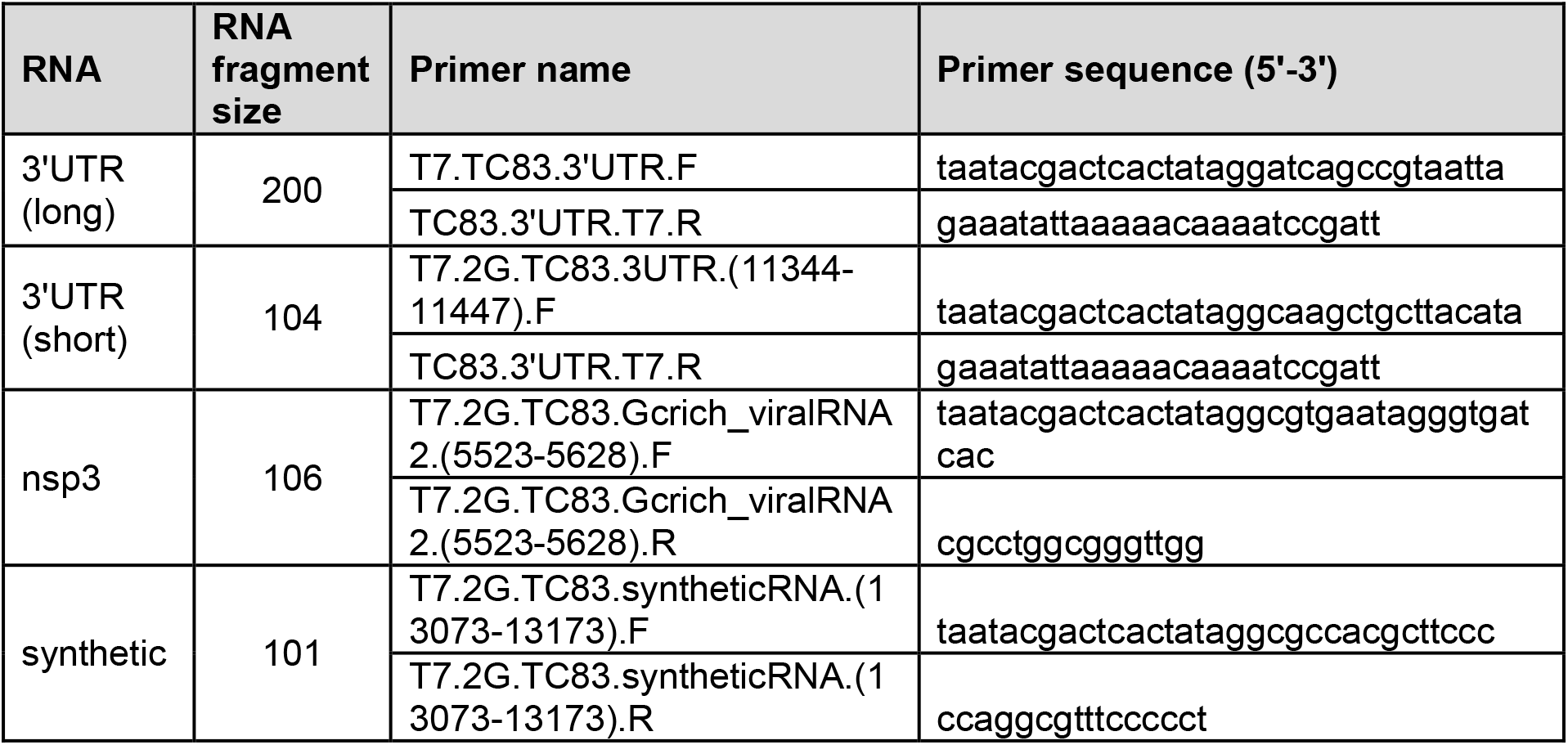
Primer sequences for RNA fragment transcription template generation.

**Table S2.**
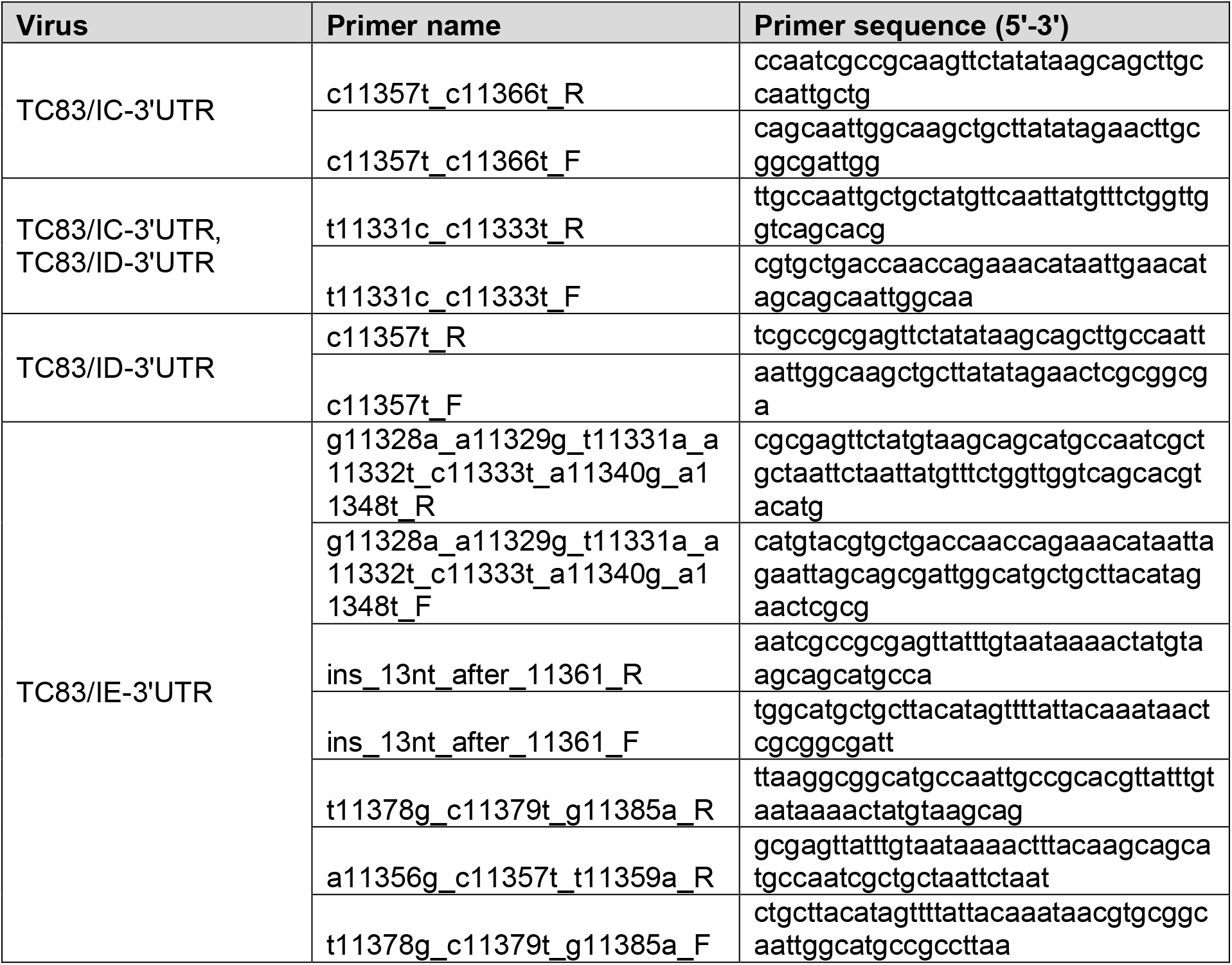
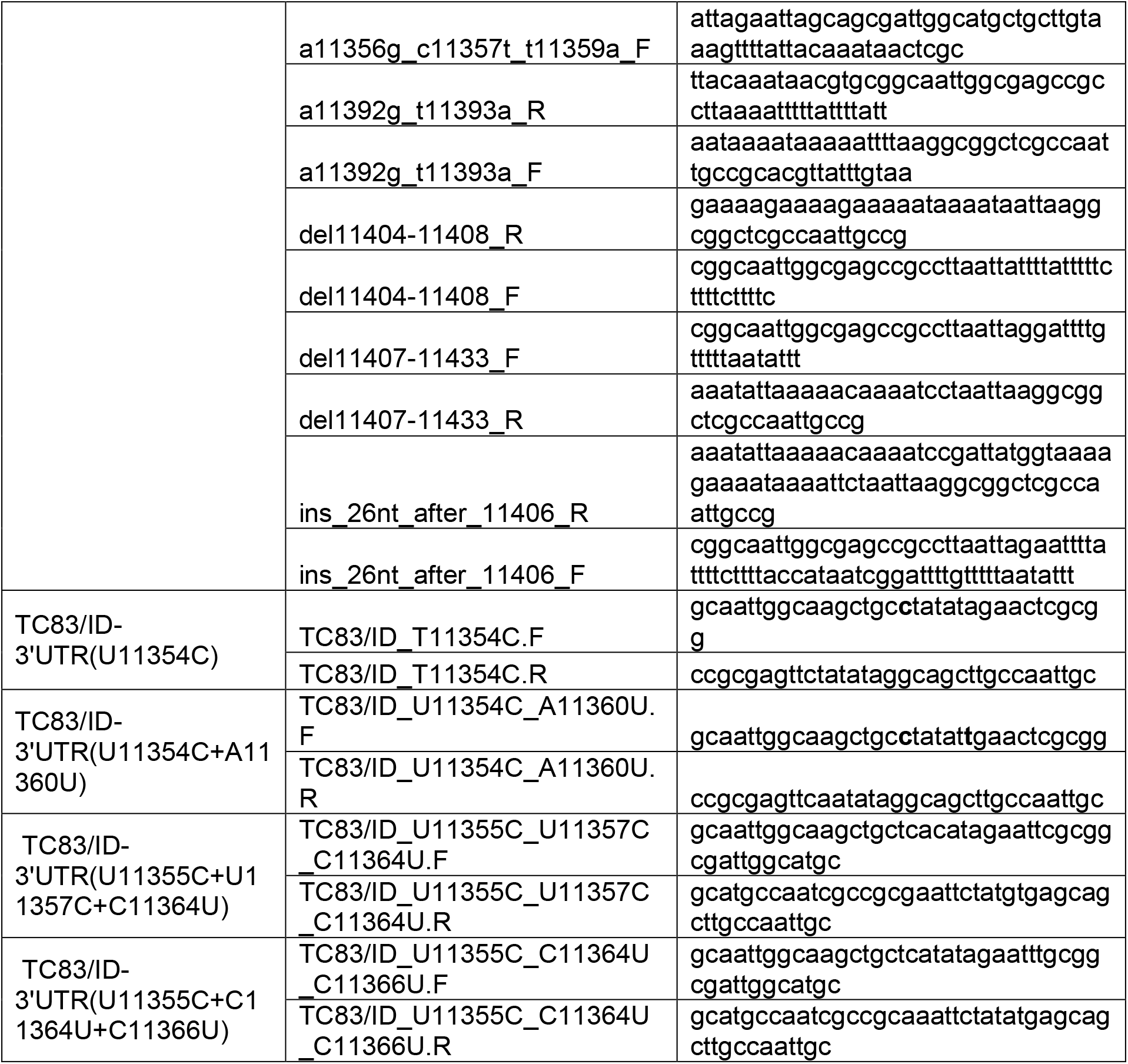
Primer sequences for mutagenesis of 3’UTR.

**Table S3. Target genes and corresponding shRNA sequences used in HeLa screen.**

**Table S4. Primary ISG screen data.**

**Table S5. Summary of 3’UTR mutants and corresponding VEEV strains, epizootic clades, and lineages which are represented by each mutant.**

## ACKNOWLEDMENTS

We thank the members of the Ram laboratory for helpful scientific discussions and technical assistance. We thank Dr. Michael Diamond for generously providing reagents for this work, and we thank Jason Smith for scientific discussion.

The authors declare that they have no conflict of interest.

