## Supplementary figures and images for "Sequence diversity in the 3’ untranslated region of alphavirus modulates IFIT2-dependent restriction in a cell type-dependent manner"

### Figure S1

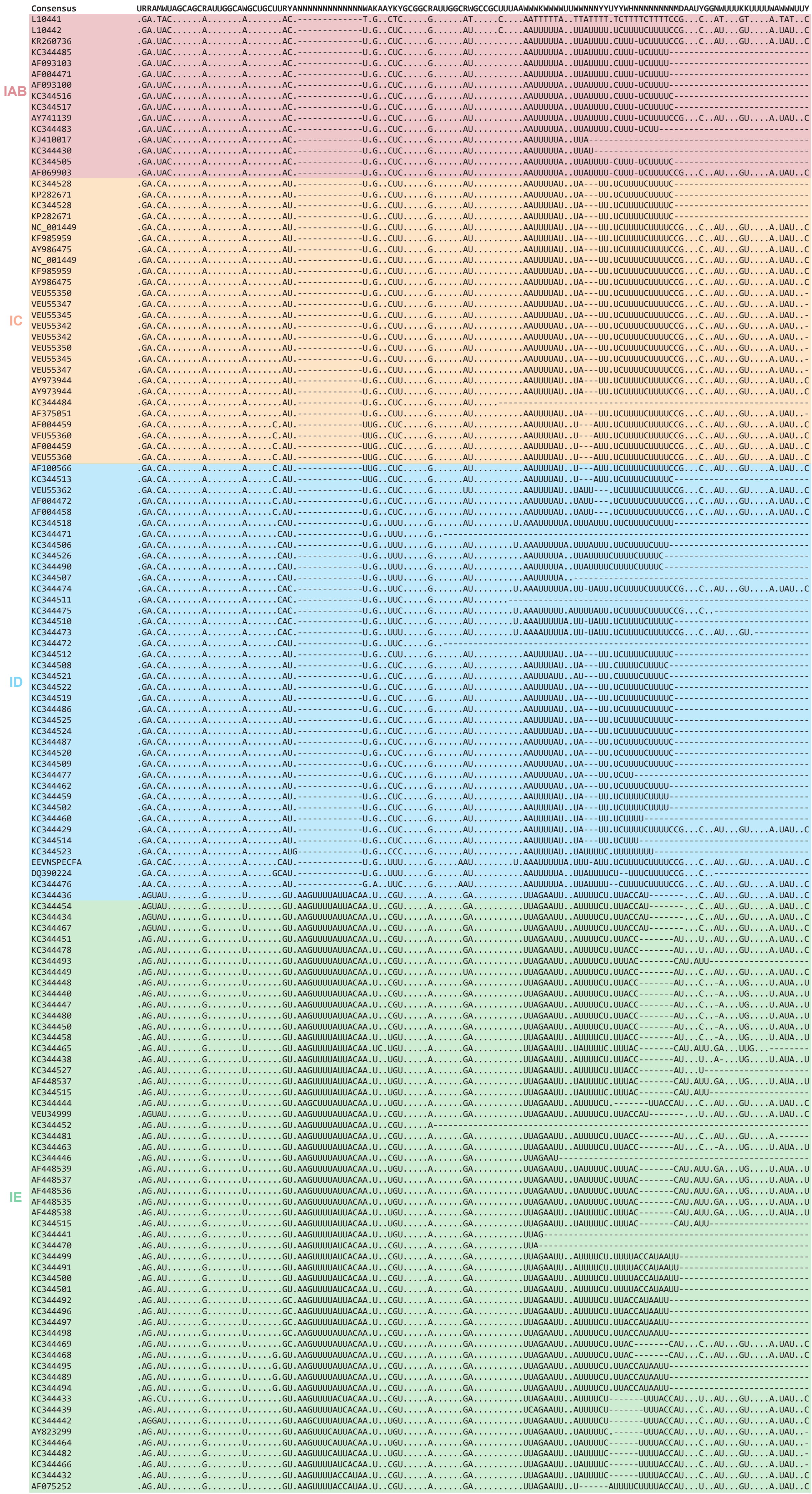


**Figure S1**

### Figure S2

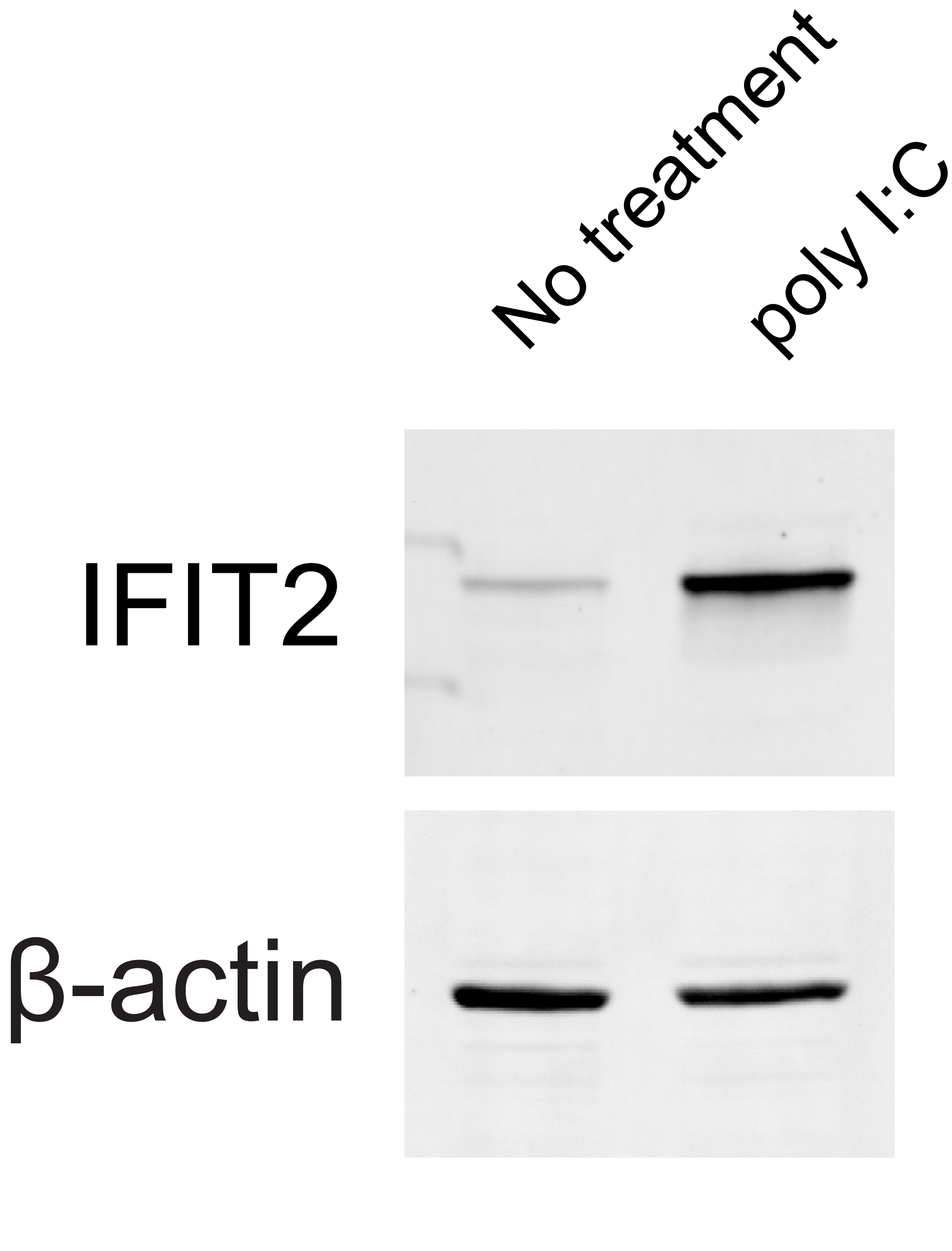


Figure S2
