## Supplementary material for "Sequence diversity in the 3’ untranslated region of alphavirus modulates IFIT2-dependent restriction in a cell type-dependent manner": Table S3

| Accession List | Gene Symbol | ShRNA Sequence |
|---|---|---|
| NM_152574,AK093459,AK298581,AK298998,AK301427,AK310482,BC038592,BC042417,BC142985 | TTC39B | CAACTTATGTGTTCTTGAA |
| NM_001223,NM_033292,NM_033293,NM_033294,AK223503,AK290114,AK290122,AK301037,AK313516,AY660536,BC041689,BC062327,CR592117,M87507,U13697,U13698,U13699,X65019 | CASP1 | GGCAGAGATTTATCCAATA |
| NM_006019,NM_006053,AF025374,BC018133,BC032465,CR594613,CR599870,U45285 | TCIRG1 | CTCTGAGGCAGGAGAGGAA |
| NM_000864,BC007720 | HTR1D | GTGAGAAACTGTTTGATTA |
| NM_001039916,NM_001039917,NM_001039918,NM_001039919,NM_001039920,NM_001135734,NM_133476,NM_175557,NM_001034830,NM_001034831,NM_133429,AK095734,BC053361,U80738,AK031726,AK156026,BC038489,BC046410,BC064013,AB019281,AF216804,AF216805,AF216806,AF216807 | ZNF384, Zfp384, Zfp384 | CCTCCCAGATAACAGAGAA |
| NM_001565,BC010954,DQ892786,DQ896033,X02530 | CXCL10 | GCCATCAAGAATTTACTGA |
| NM_005465,NM_181690,NM_011785,NM_031575,AF085234,AF124141,AF135794,AJ245709,AL117525,AM392791,AM392849,AY005799,BC020479,BC121154,AF124142,BC066861 | AKT3, Akt3, Akt3 | CTATGAAGATTCTGAAGAA |
| NM_004048,AB021288,AF072097,AK315776,AY007153,AY187687,BC032589,BC064910,CR457066,CR590254,CR592576,CR592679,CR593405,CR593761,CR593874,CR594444,CR596238,CR596347,CR596444,CR596717,CR597808,CR599099,CR599510,CR603043,CR603887,CR604555,CR604705,CR604949,CR605353,CR606667,CR607236,CR608139,CR608197,CR610094,CR612132,CR613616,CR613782,CR614402,CR614556,CR615574,CR615578,CR615729,CR616171,CR616437,CR619071,CR619828,CR620994,CR621079,CR623073,CR624595,CR625152,CR626821,DQ839493,DQ884406,DQ890551,DQ893711,S54761,S82300,V00567,X07621,BU658737 | B2M | GCTATCCAGCGTACTCCAA |
| NM_002201,NM_001008510,XM_001064354,AK307769,BC016341,BT006952,CR456942,X89773 | ISG20, Isg20 | CTGCACAAGAGCATCCAGA |
| NM_001974,AK131562,AK290401,AK291518,AK313495,BC059395,X81479 | EMR1 | CGTAGTTTCTCTGAAGAAT |
| NM_014705,AB018259,AK091557,AY233380,BC117688,BC117689 | DOCK4 | GGAATGTCATGATTCCAAT |
| NM_003113,AF056322,CR616734,L79988 | HMG1L6, SP100 | GGACAAGGCCCATTATGAA |
| NM_017878 | HRASLS2 | GGGAAAGGCAATAAATCCA |
| NM_001025079,NM_001777,NM_198793,AK289813,BC010016,BC012884,BC037306,BT006907,CR591693,CR591788,CR592842,CR601569,CR615658,CR621937,CR623329,CR624381,DQ890756,DQ891811,DQ893920,X69398,Z25521 | CD47 | GTCCCAGGTGAATATTCAT |
| NM_004060,NM_199246,AK297190,AK312913,BC000196,BC007093,BT007134,BX538035,CR619041,D78341,DQ893765,EU176176,L49504,U47413,U53328,X77794 | CCNG1 | CACAGAAGTGTGTAGAGTT |
| NM_002717,AK303981,AK314823,AY189686,BC041071,M64929,BX440554 | PPP2R2A | GCCAGTCCACGAAGAATAT |
| NM_007315,NM_139266,AK096686,AK225853,AK292604,AK315002,BC002704,BT007241,CR601119,CR618286,CR749636,EU831980,EU832073,M97935,M97936,BX400908 | STAT1 | GCCCTAAAGGAACTGGATA |
| NM_001127397,NM_001127398,NM_015701,AK001913,AK315477,AY358717,AY453410,BC013129,BC022228,CR597633,CR601764,CR604548,CR608194,CR613668,CR616366,DQ894762 | C2orf30 | CAAGAACCTTCTATTTGAA |
| NM_001562,AF077611,AF380360,AY044641,AY266351,BC007007,BC007461,BC015863,CR541973,CR542001,D49950,U90434 | IL18 | GACATGATAATAAGATGCA |
| NM_001775,AK297592,AK303169,AK313268,BC007964,CR606718,CR610722,CR626710,D84276,D84277,DQ893184,EU176777,M34461,CD688069 | CD38 | CTTTCAAGGGTGCATTTAT |
| NM_022831,XM_001725626,XM_001726789,XM_930476,XR_015691,XR_017594,XR_038764,NM_181732,XM_001473773,XM_905339,NM_001127600,AK022868,AL833718,BC015535,BC043142,AK027971,AK076031,AK081494,AK085039,AK143716,AK149788,AK150639,AK150702,AK151783,AK152961,AK153448,AK158103,AK159318,AK167520,AK172451,BC057183,BC086763 | AIDA, LOC646890, LOC653631, Aida, LOC631071, LOC682999 | GATACAGGTTTATTGTAGA |
| NM_001547,AK300431,AK312831,BC005987,BC032839,EB387611 | IFIT2 | CACTTTATAGAGGGTGTAA |
| NM_020119,NM_024625,AK023350,AK292811,BC002935,BC025308,BC033105,BC038809,BC040956,BX571742,BX647138,BX647974 | ZC3HAV1 | CACTTGTTAACGATTCTTT |
| NM_000416,AF056979,AK222803,AK297052,AK312674,BC005333,BT006814,CR601864,CR624388,CR625186,EF535103,J03143 | IFNGR1 | GAAGTGAGATCCAGTATAA |
| NM_014795,AB011141,AB056507,AB193095,AB193096,AK308124,AK308445,BC037975,BC127101,BC127102 | ZEB2 | GCAAGGCCTTCAAATATAA |
| NM_001031683,NM_001549,AF026939,AF083470,AK290427,AK297137,AK312675,BC001383,BC004977,BT007284,CR618361,CR626380,U52513 | IFIT3 | GCTACTGCAACCTTCAGAA |
| NM_001547,AK300431,BC032839 | IFIT2 | CAGCCAAATCCTTCATGTA |
| NM_020140,NM_152788,AF145204,AF164792,AK294994,BC142669 | ANKS1B | CTGATAAAGCCTAAATAGA |
| NM_001136540,NM_001136541,NM_003661,NM_030882,NM_145343,NM_145637,AB208990,AF019225,AF305224,AF305225,AF305428,AF305429,AF323540,AF324223,AF324224,AK056938,AK300454,AK303919,AK303978,AK313752,BC004395,BC017331,BC112943,BC113867,BC114475,BC127186,BC141823,BC142720,BC143038,BC143039,CR593100,CR626091,DQ893431,DQ896744,BG419709,DA375861,DC340792,DC367266 | APOL1, APOL2 | CCCAGAGAGCAGTATCTTT |
| NM_001647,AK294523,AK312090,BC007402,BT019860,BT019861,CR456838,CR541773,DQ892668,DQ896124,J02611,BU195270,DA892855 | APOD | CAGGCCAACTACTCACTAA |
| NM_033109,AJ458465,AK307589,AY027528,BC009057,BC053660,CR625599 | PNPT1 | CCTGATGTCCTAGCAATTA |
| NM_006084,BC035716,CR596057,CR602478,CR603020,CR609755,CR610511,CR619285,M87503,BI497082 | IRF9 | CTGTCCTCTTTGTGATAAT |
| NM_001127397,NM_015701,AK001913,AK315477,AY453410,BC013129,CR604548,CR608194,CR613668,CR616366,CR623209,DQ894762 | C2orf30 | GAAGATTTGCAATCAACTA |
| AF061738 | LAP3 | GTGTCTGCTGCAAAGTTAA |
| NM_021100,AF097025,AJ010952,AK001265,AK001470,AK056242,AK091049,AK223239,AK295215,AK297969,AK301969,AK302023,AK312293,BC065560,CR594451,CR610756 | NFS1 | CCTCTTATGTGCTTAGAGC |
| NM_001127397,NM_001127398,NM_015701,AF131743,AF131849,AK001913,AK315477,AY358717,AY453410,BC013129,BC022228,CR597633,CR601764,CR604548,CR608194,CR613668,CR616366,CR623209,DQ894762 | C2orf30 | CCTCACTCCTGTCAATATA |
| NM_005419,AK094039,BC051284,BX640607,CR621797 | STAT2 | CCAAGTCTGTGGAACTTAA |
| NM_022831,XM_001725626,XM_001726789,XM_930476,XR_015691,XR_017594,XR_038764,AK022868,AK289865,AL833718,BC015535,BC043142,BC067805 | AIDA, LOC646890, LOC653631 | CTGGGCCAATTGTAATAGA |
| NM_001142315,NM_001142316,NM_005574,AF257211,BC034041,BC035607,BC042426,BC073973,CR604507,CR614368,CR625714,X61118 | LMO2 | GCCCTTGAGGGAGAACTTA |
| NM_000600,BC015511,BT019748,BT019749,CR450296,CR590965,CR626263,DQ891463,DQ894639,M14584,M18403,M29150,M54894,S56892,X04403,X04430,X04602 | IL6 | CAAAGAATCTAGATGCAAT |
| NM_001165,NM_182962,BC037420,L49432,U37546,U45878 | BIRC3 | CAACACGTTTGAACTGAAA |
| NM_005204,AK290320,AK313295,BC104833,BC113566,CR542284,CR619928,D14497,Z14138 | MAP3K8 | GTGTCAAGACAGTAATCAA |
| NM_000168,AK299299,AK308429,BC113616,BC117168,M57609,AA330409 | GLI3 | CCTCCACATCCCTACATTA |
| U36500 | SP140 | GTCAGTTTCTTGTTTCTCT |
| NM_015459,AK023383,AK097588,AK301910,AL117600,BC077727,CR936784 | ATL3 | CACTGATACCGTATGTATT |
| NM_001003818,NM_001003819,NM_058166,AB039903,AF220030,AK023210,AK027664,AK290172,AK293295,AK298301,BC047564,BC065575,BC136871,CR624250,CR749260,CX760618 | TRIM6, TRIM6-TRIM34 | CTACAAAGCTGAGAAGTAT |
| NM_020830,AB037856,AK022888,AK023415,BC040525,BC065934 | WDFY1 | CTCATAAATTGCTTTCCTA |
| NM_020140,NM_152788,AF145204,AF164792,AK294994,BC142669 | ANKS1B | CCTTCACTTTAGTGGTCTA |

| | | |
|---|---|---|
| NM_004458,NM_022977,AB061713,AB061714,AF030555,AK292070,AK294915,AK307566,BC034959,DQ890835,DQ893990,Y12777 | ACSL4 | GACAATAAGGCTATCAATA |
| NM_015474,AB013847,AB208944,AF228421,AK027811,AK304187,AK311150,AK315169,AL050267,BC036450 | SAMHD1 | CTGATTCGAGTATATTGTA |
| NM_033064,AB058775,AK027889,AK092309,AK125457,BC008736,BC026217 | ATCAY | CTCCGACCTTATGTCCAAA |
| NM_001042500,NM_004084,NM_005217,BC027917,BC093791,BC112188,BC119706,M21130,M21131,M26602,X13621,X52053 | DEFA1, DEFA3, LOC728358 | CTTTGAGAGCTACAGGGAA |
| NM_000463,NM_001072,NM_007120,NM_019075,NM_019076,NM_019077,NM_019078,NM_019093,NM_021027,NM_205862,AF030310,AF056188,AF462267,AF462268,AK025403,AK290834,AK313488,AK313510,AK313623,AY435136,AY435137,AY435139,AY435141,AY435143,AY435144,BC011409,BC019861,BC020971,BC043491,BC058844,BC069210,BC121036,BC128414,BC128415,BC131623,BC139784,BC141470,BC146418,BC153172,BC156848,BC166641,J04093,M57899,M57951,S55985,U89507,U89508 | UGT1A1, UGT1A10, UGT1A3, UGT1A4, UGT1A5, UGT1A6, UGT1A7, UGT1A8, UGT1A9 | GAGTTAAGAAAGCCCACAA |
| NM_006985,NM_178541,NR_003610,XR_016512,XR_016550,XR_041580,AF132984,AK124516,AK131084,AK131085,AK160377,AK302708,BC039707,BC063633,BC146422,CR619663,BE253379 | LOC339047, LOC399491, LOC642778, LOC642799, NPIP, PDXDC2 | GCCGTAGACAGGAAGGAAT |
| NM_003895,NM_203446,AB020717,AF009039,AF009040,BC098395 | SYNJ1 | CCCTGATTGATATAGATAT |
| NM_002468,AK097983,AK124685,AL832906,BC013589,BX537602,CR619528,U70451 | MYD88 | CTAACCATGTCCCTGAACA |
| NM_024911,AK026744,AK074583,AK074984,AY359035,BC007211,BC110826,BC137109,BC137113,BX648748,CR591378,CR602492,DQ323735 | GPR177 | CAAGTAAGATTTACTGTAT |
| XM_001723375,XM_001724221,AK294177,AK308629 | LOC100132540, LOC100132620, LOC23117, NPIP | CAGAGGTTCCGTATTTGTA |
| NM_004120 | GBP2 | CAATTCAACTCATGCTTAT |
| NM_004048,AB021288,AK026463,AY007153,BC032589,BC064910,CR590254,CR591572,CR592576,CR592679,CR593405,CR593761,CR593874,CR594444,CR596238,CR596347,CR596444,CR596717,CR597808,CR599099,CR599510,CR599875,CR603043,CR603887,CR604110,CR604555,CR604705,CR604949,CR605353,CR606667,CR607236,CR608139,CR608197,CR610094,CR610959,CR612132,CR612519,CR613616,CR613782,CR613955,CR614402,CR614556,CR615574,CR615578,CR615729,CR616171,CR616437,CR618556,CR619071,CR619828,CR620994,CR621079,CR623073,CR624595,CR625152,CR626821 | B2M | GTGCATAAGTTAACTTCCA |
| NM_003335,BC006378,BT007026,DQ893307,DQ893707,L13852 | UBA7 | GCCAGATTATCCCAGCCAT |
| NM_002468,AK097983,AK124685,AL832906,BC013589,BX537602,CR619528,U70451 | MYD88 | GACCCTAAATCCAATAGAA |
| NM_002163,AK313524,BC126247,M91196,BG107396,BI835080 | IRF8 | CAGATTGACAGTAGCATGT |
| NM_001548,AK308688,AK314588,BC007091,BT006667,CR591778,CR593386,CR601585,CR617896,CR621649,CR623965,DQ893534,DQ893755,X03557 | IFIT1 | GTCAATGCAATTATCCATT |
| NM_001032394,NM_001032395,NM_020455,NM_198569,AB183546,AB183547,AB183548,AB183549,AK075087,AK092519,AL080079,BC075798,BX648315 | GPR126 | CTTAGACATTCATAGTAGA |
| NM_001141969,NM_001141970,NM_001350,NR_024517,AF006041,AF015956,AF039136,AF050179,AF097742,AK223404,AK292187,AK303767,AK303854,AK313034,BC000220,BC073776,BC109073,BC109074,CR457085,EU446949 | DAXX | GGCCATTAGGAAACAGCTA |
| NM_000161,NM_001024024,NM_001024070,NM_001024071,BC025415,U19523,U66095,U66097 | GCH1 | CGTATAGATGGTATAGGTA |
| NM_002985,AF043341,AF266753,AK312212,BC008600,DQ230537,DQ891490,DQ894681,M21121 | CCL5 | GGGAGTACATCAACTCTTT |
| NM_000566,AK291451,AK291502,BC032634,BC110416,BC152383,CR608315,CR622515,DQ786309,DQ890842,DQ893997,L03418,X14355,X14356 | FCGR1A, FCGR1B | CTCTGAATACCAAATACTA |
| NM_001135865,NM_006985,NM_130464,NM_178541,NR_002555,NR_002603,NR_003610,XM_001132754,XM_001717235,XM_001717595,XM_001717652,XM_001717676,XM_001720898,XM_001720909,XM_001720910,XM_001723013,XM_001723016,XM_001723375,XM_001723497,XM_001724221,XM_001732844,XM_001732845,XR_015786,XR_015889,XR_016512,XR_016550,XR_041580,AB209632,AF132984,AF229069,AK124516,AK128772,AK131084,AK131085,AK160377,AK294452,AK295170,AK296301,AK296338,AK303264,AK303500,AK303572,AK304162,BC008178,BC010188,BC036263,BC039707,BC061522,BC063633,BC094882,BC146422,BC157836,CR619663,BM557638,BQ721066 | LOC100132247, LOC100132540, LOC100132620, LOC100133970, LOC23117, LOC339047, LOC388237, LOC399491, LOC440345, LOC440353, LOC613037, LOC642778, LOC642799, LOC728734, LOC728741, LOC728888, LOC729602, LOC729978, LOC730153, NPIP, PDXDC2 | CTCACAGCTGAAACTTTAA |
| NM_001647,AK294523,AK312090,BC007402,BT019860,BT019861,CR456838,CR541773,DQ892668,DQ896124,J02611,BU195270,DA892855 | APOD | GGAAAGATCAAAGTGTTAA |
| NM_001548,AK092813,AK308688,AK314588,BC007091,BT006667,CR591778,CR593386,CR601585,CR617896,CR621649,CR623965,DQ893534,DQ893755,X03557 | IFIT1 | GCATCATTAACAAGGGATA |
| NM_021105,AB006746,AF098642,AK313377,BC017901,BC021100,BC032718,CR593697,CR607459,CR623341,DQ890879,DQ894033 | PLSCR1 | CTGTCCACCTGGATTAGAA |
| NM_001024912,NM_001712,AL833584,X14831,X16354 | CEACAM1 | CCCTTCAGCGATTATTTAA |
| NM_017912,AF336798,AK000644,AK097168,AK225506,AK295832,AL833664,AY653201,AY653202,AY653203,BC035775,BC042047,BX647121 | HERC6 | CCATCATTGGAAGATTTAA |
| NM_005465,NM_181690,AF085234,AF124141,AF135794,AJ245709,AK308052,AL117525,AM392791,AM392849,AY005799,BC020479,BC121154,AW960221 | AKT3 | GATATAAAGAGAAACCTCA |
| NM_004079,AK024855 | CTSS | CAGGCAGCAGGAACTTTAA |
| NM_020414,AB209382,AF134475,AF145022,AF214731,AK025162,AK296726,AK297232,AK304504,AL136886,AM393538,BC008847,BC009406,BC096826,BX537533,CR596324,CR615921,CR619609,CR624535,DQ893523,DQ896510 | DDX24 | GCCAGGTTTACAGGAATTA |
| NM_005858,NM_019774,NM_053855,BC037270,Y11997,AB028920,AK004801,AK089092,BC042602,BC060170,BC060209,BC078448,BC125521,BC137746,U01914 | AKAP8, Akap8, Akap8 | CAGCTGGCAAGGTTATGAA |
| NM_000304,NM_153321,NM_153322,AK300690,BC019040,BC091499,CR592810,CR597127,CR597311,CR598725,CR602788,CR602976,CR603134,CR604376,CR604692,CR604834,CR608530,CR610349,CR611493,CR611595,CR612069,CR615858,CR618437,CR618628,CR618649,CR622251,CR624540,D11428,L03203,M94048,X65968 | PMP22 | CAGTCCACCTCATTTAGAA |
| NM_080745,NM_182985,AF302088,AF302089,AK226115,AK292252,AL360161,AY305385,BC024199,BC033314,BC047945,DQ232883,EU446573,EU832108,EU832202 | TRIM69 | CAGAGAGCTTATTTCCAGA |
| NM_030882,NM_145637,AB208990,AF305225,AF305429,AF324223,AF324224,AK056938,AK303978,BC004395,CR593100,DQ893431,DQ896744 | APOL2 | GGAACACACCCAATGTTCT |
| NM_014795,AB011141,AB056507,AB193095,AB193096,AK308445,BC127101,BC127102 | ZEB2 | GAGACAGATCAGTAATATA |
| NM_022091,AK023189,BC050681,BC125211,BC125212,BC130038 | ASCC3 | ATATTTGACTATTAAAGGA |
| NM_001012509,NM_016180,AF172849,BC064405,BU166522 | SLC45A2 | GTCTTTACTTCACGGGATA |
| NM_016376,AB033081,AB037360,AK025483,AK057047,AK225170,AK292930,BC060812,BC148355,BC152991,CR933717 | ANKFY1 | CCCACCGCCTGCTTTCTTA |
| NM_014705,AB018259,AK091557,AY233380,BC117688,BC117689 | DOCK4 | CTCAGTATTTGCAGATATA |
| NM_052941,AF288814,AK131094,AK312417,AL832576,BC008421,BC017889,BC050625,BC070055 | GBP4 | CCTACAAATGACAAGCAAT |
| NM_017554,AB033094,AK098816,AK304269,AY134858,BX648758,DQ063584,DQ063585 | PARP14 | CTTCCTTAGTTATTGACTA |
| NM_052941,AF288814,AK131094,AK304245,AK312417,AL832576,BC050625,BC070055 | GBP4 | CAGTTAAATAAAGAGATTA |
| NM_001135651,NM_002759,AK290655,AK313818,BC007769,BC022314,BC040851,BC057805,BC093676,BC101475,CR600308,M35663,M85294 | EIF2AK2 | GTTATTAAACGTGTTAAAT |
| NM_003150,NM_139276,NM_213662,AK024535,BC008044,BC067119,BC107775 | STAT3 | AAGTTCATGGCCTTAGGTA |
| NM_032036,AF208232,AF238978,BC022800,BC032626,BN000226,BE737110 | IFI27L2 | CTCTCCACATCATCCAACA |

| | | |
|---|---|---|
| NM_004688,AB451305,AB451436,AK291548,BC001268,BC021987,CR604696,U32849 | NMI | CATTCAAATGGATGAAGAA |
| NM_007315,NM_139266,AK096686,AK225853,AK292604,AK315002,BC002704,BT007241,CR601119,CR618286,CR749636,EU831980,EU832073,M97935,M97936,BX400908 | STAT1 | CTGAAGTATCTGTATCCAA |
| NM_015668,AK097399,AK292559,AL117544,AY009106,BC036665,BC050053,BC092411 | RGS22 | GCCTCAGTTCTGTGAGTTT |
| NM_021964,AF039019,AF432210,AJ236885,AK314203,AL833616,BC050260,BC128582,L04282 | ZNF148 | CTGTGCATAGTAGTACTAA |
| NM_006084,AK295467,BC035716,CR596057,CR602478,CR603020,CR609755,CR610511,CR619285,EU446692,M87503 | IRF9 | CTGCTCACCTTCATCTACA |
| NM_022750,AL136766,AL137255,BC044660,CR602813 | PARP12 | CTGGTCTATGGCACAACTA |
| NM_005746,AK292851,BC020691,BC072439,BC106046,CR593609,CR606862,U02020 | NAMPT | CATCTTCCAATAGAAATAA |
| NM_006121,AK313986,BC063697,M10938 | KRT1 | CAAATCAAGTCACTCAACA |
| NM_017523,NM_199139,AK056908,AK290276,AK292710,AK292848,BC032776,BC058017,BC073156,BX648588,BX649188,EF028165,EF028166,X99699 | XAF1 | CTATGACATTCTGAGGAGA |
| NM_001127713,NM_015915,NM_181598,AF131801,AF444143,AK223436,AK290185,AK312518,AL833591,AY032844,BC010708,CR457153,CR599776 | ATL1 | CTGTGCACTTGGCATATA |
| NM_001040020,NM_014888,BC024200,BC068526,D87120 | FAM3C | CTATTCGGATAATTTCTTA |
| NM_002164,AK313259,AY221100,BC027882,CR591703,CR613807,CR614485,CR616634,CR621546,M34455,X17668,CD690354 | INDO | GTAAGGTCTTGCCAAGAAA |
| NM_018284,AK309096,AL136680,BC063819,CR936755,CR936758 | GBP3 | CTAGGTAAGTGGTTGACAT |
| NM_002308,NM_009587,NR_024043,AB003517,AB005894,AB006782,AB008492,AK097892,AK126017,AK223232,AK296031,BC034392,BC105942,BC105944,BC110340,CR597107,CR598297,CR602208,CR604285,CR608609,CR614567,CR616420,CR622120,CR626506,Z49107 | LGALS9 | CTGACCCATGTGCAGACAT |
| NM_001562,AF077611,AF380360,AK311301,AY044641,AY266351,BC007007,BC007461,CR541973,CR542001,D49950,U90434 | IL18 | GCAATGAAATTTATTGACA |
| NM_021105,AB006746,AF098642,AK300181,AK313377,BC021100,BC032718,BC070251,CR593697,CR607459,CR623341,DQ890879,DQ894033 | PLSCR1 | CTTTAGACCTTGATGTTAA |
| NM_198584,AK095314 | CA13 | CTATTTACACTGTAATGAA |
| AJ344101 | TXNDC17 | CTCAAAGGAACGTGATTCT |
| NM_198584,AK095314 | CA13 | CATTGTTTGACATATATTA |
| NM_014398,AB013924,AJ005766,BC032940 | LAMP3 | CTGACTTTAGTCTTAATAA |
| NM_001030287,NM_001040619,NM_001674,NM_004024,AB066566,AB078026,AB078027,AB209032,AK294296,AK312998,AY313926,AY313927,BC006322,BT006996,CR450334,CR600751,CR602765,CR612905,CR614684,CR614862,CR620886,CR626784,DQ892723,DQ895972,L19871,AU135799,DA691940,DB197400 | ATF3 | CGACGAGAAAGAAATAAGA |
| NM_014547,AF177171,AF237631,AK310344,AK312569,AL137543,BX647277,CR611644 | TMOD3 | CAGAGCAGCTAATGCTATA |
| NM_001035267,NM_021104,AF026844,AK023340,BC014383,BC015984,BC017820,BC032611,BC062462,BC064907,BC066328,BC070252,BC070253,BC070254,BC070255,BC070483,BC082759,BC105279,BC105286,BC105296,BC105794,BC105799,BC142666,BC142726,CR599786,CR605341,CR609127 | CTTNBP2NL, RPL41 | CGAGTGTAACAACCATATA |
| NM_017414,AF176642,AJ243526,AK313385,AL136690,AM393395,BC014896,BT006835,CR457216,CR604420,CT841507,CU013072,CU013360,BI752849 | USP18 | GACTTCACCAGGATATTGA |
| NM_021629,AK001890,AK022599,CR591019 | GNB4 | CTGATAGCTTCATGTTTAA |
| NM_080745,NM_182985,AF302088,AF302089,AK226115,AK292252,BC033314,BC047945,DQ232883 | TRIM69 | ACAGTAATGAGTCATAATA |
| NM_021181,AK301438,AL713801,AL833025,AL834424,BC027867 | SLAMF7 | ATGTTTGGCAGATACTATA |
| NM_000577,NM_173841,NM_173842,NM_173843,AF043143,AK290898,AK290926,BC009745,BC068441,CR605915,CR616671,CR619093,CR625646,DQ892566,DQ895781,M55646,X52015,X53296,X84348,BI766516 | IL1RN | CAAATGTCAATTTAGAAGA |
| NM_004458,NM_022977,AB061713,AF030555,BC034959,CR603171 | ACSL4 | CAACTAAAGTTGTACTTTA |
| NM_174975,AK131358,AK298079,AY158086,AY240872,BC069641,BC101001,BC101002,BC101003,BC101004 | SEC14L3 | CTCATCACACCAAGTGGAA |
| NM_004217,AB011450,AF004022,AF015254,AK297976,BC000442,BC009751,BC013300,BC080581,CR591471,CR600836,CR603941,CR605805,CR617702 | AURKB | ATCCCTAACTGTTCCCTTA |
| NM_002360,AK056767,AL831913,BC012777,BC094871,BM981655 | MAFK, TMEM184A | CAGTTCCATTTCATTATTT |
| NM_022347,AK289793,BC096704,BC098170,BC098309,BC098348 | IFRG15 | ACCAGATGGACCTTTGAGA |
| NM_001775,AK297592,AK303169,AK313268,BC007964,CR606718,CR610722,D84276,D84277,DQ893184,EU176777,M34461 | CD38 | CTGTTTCAGTATTCTGGAA |
| NM_001077269,NM_003387,AK097901,AL832275,BC045584,BG820214,BM475558 | WIPF1 | GAATTTGCCCTTCCTAGAA |
| NM_000146,XR_017149,XR_019548,XR_037197,AK026534,AK130191,AK130205,AK131048,AK131050,AK131053,AK307065,AK311773,AY207005,AY466472,BC002991,BC004245,BC008439,BC008441,BC013928,BC016346,BC016354,BC016715,BC018990,BC021670,BC050625,BC058820,BC062708,BC067772,BC105971,BX571748,CR456715,CR596451,DQ892980,DQ896228,EU831925,EU832020,M10119,M11147,Y09188 | FTL, LOC392437, SEC62 | CTCCCAGATTCGTCAGAAT |
| NM_001025079,NM_001777,NM_198793,AK289813,BC010016,BC012884,BC037306,BT006907,CR591788,CR592842,CR601569,CR615658,CR621937,CR624381,DQ890756,DQ891811,DQ893920,X69398,Z25521,BI753986,CB995865 | CD47 | CTCAGCTACTATTTAATAA |
| NM_002984,AY766446,AY766448,BC027961,BC107433,J04130,M23502,M25316,M57503,X16166,X53683 | CCL4 | GTGTCATTTCCATTATTTA |
| NM_018381,AK002148,AK096142,AK098062,AL137694,BC010847,BC026180,CR590846,CR592012,CR592313,CR594187 | C19orf66 | CAACTTTCTTCAGAGTAAT |
| NM_006877,BC008281,CR594139,CR610393,EU831974,EU832067,M24470 | GMPR | CAATGGGTATTCAGAACAT |
| NM_019111,AF522250,AK297032,AK313123,BC032350,BC071659,CR457013,CR590143,CR594558,CR613451,CR622870,DQ891595,DQ894789,J00194,K01171,M60334,V00523,BM849755 | HLA-DQA1, HLA-DRA | CAATGTACCTCCAGAGGTA |
| NM_024586,NM_148904,NM_148905,NM_148906,NM_148907,NM_148908,NM_148909,AB208851,AF392445,AK022554,AK027535,AK027707,AK056617,AK091703,AK124037,AK128043,AK315167,AY178997,BC025978,CR457288,CR609523 | OSBPL9 | CAGCTCGGGAAATAGTCTA |
| NM_005384,AK313970,BC008197,CR542049,CR599259,CR608714,CR614563,DQ891105,S79880,U26173,X64318 | NFIL3 | CACACAAGCTCCGGATCAA |
| NM_000386,AK223403,AK301948,AK312896,BC003616,BT007018,X92106 | BLMH | CAACCAGAAGGATGAATGA |
| NM_002535,NM_016817,AK292796,AK292906,BC049215,M87284,M87434,BQ213828 | OAS2 | CTTCAACGCTCTGAGCTTA |
| NM_001974,AK131562,AK290401,AK291518,AK313495,BC059395,X81479 | EMR1 | CAGAGCATGCAACTTGTAA |
| NM_017654,AB095925,AF445355,AK122951,AK125101,AK125131,BC132773,BC132775,BC150249,BX647072 | SAMD9 | GTGATTATCCTAAATTGTA |
| NM_000062,NM_001032295,AB209826,AK293054,AK303840,BC011171,CR590941,CR612675,CR614379,CR619147,M13656,M13690,X07577 | SERPING1 | AAGTTCACCAGACTCTATA |
| NM_005533,AK025406,BC001356,DQ892598,DQ895826,U72882 | IFI35 | CTCTGAGAGTCTCTCCGTA |
| NM_024586,NM_148904,NM_148905,NM_148906,NM_148907,NM_148908,NM_148909,AB208851,AF392445,AK022554,AK027535,AK027707,AK056617,AK074392,AK091703,AK124037,AK128043,AY178997,BC006160,BC025978,CR607539,CR609523 | OSBPL9 | CTGTTCATTCCAATCTTCT |
| NM_002443,AK312108,BC005257,BT006816,CR457015,DQ892498,DQ895711,M15885,S67815,U78976 | MSMB | CTACGAAACAGAAATTTCA |

| | | |
|---|---|---|
| NM_032206,AF389420,AK025362,AK074182,AK090439,AK097030,BC156512,EF452236 | NLRC5 | CAACATGACTCAGCTCTAT |
| AK021862,BC098263,BC098294,BC098363 | FAM106A | CAATTCAATGGTGTTTAGA |
| NM_020897,AB040968,AK055840,BC000066,BC028024,CR625647,BF110802 | HCN3, PKLR | CTCTCCTACTCTACAGCAT |
| NM_002535,NM_016817,AK292796,AK292906,BC049215,M87284,M87434 | OAS2 | GTGTTCCATAACTCACTTA |
| X14831,X16354 | CEACAM1 | GGCAGTAATGCTTCTCCTA |
| NM_022147,AJ251832,BC013161 | RTP4 | GTACATAATGGATTTAGTA |
| NM_018284,AK001823,AK309096,AL136680,BC017992,BC140837,CR936755,CR936758,EU832447,EU832454,EU832530,EU832536 | GBP3 | CTAGTGCTGACCTATATCA |
| NM_001572,NM_004029,NM_004031,AF076494,AK303752,U53830,U53831,U53832,U73036 | IRF7 | CGCCTAGAACCCAGTCTAA |
| NM_021100,AF097025,AK001265,AK001470,AK056242,AK223239,AK295215,AK297969,AK301969,AK302023,AK312293,BC018471,BC065560,CR594451,CR610756 | NFS1 | CTCTCTATATGGATGTGCA |
| NM_021105,AB006746,AF098642,BC021100,BC032718,BC070251,CR607459 | PLSCR1 | ATATGCATATTATCTTTAA |
| NM_015459,AK023383,AK097588,AK301910,AL117600,BC077727,CR936784,BX642720 | ATL3 | CTTCTTCTCATATCGGTAA |
| NM_004509,NM_004510,NM_080424,AF280094,AF280095,AK026488,AK301097,AL832300,BC012447,BC019059,CR625933,L22343 | SP110 | GGCTCTATGCCAGAGATAA |
| NM_006573,AF116456,AF132600,AF134715,AF136293,AF186114,AK309629,AY129225,AY302751,AY358881,BC020674,CR541818,DQ891413,EU176634,BI818459 | TNFSF13B | CTTACTTCTTGCCTTAAGA |
| NM_001008540,NM_003467,AF147204,AK129916,AK296674,BC020968,CR594428,CR594588,CR596547,CR598681,CR598713,CR601301,CR604071,CR605131,CR610268,CR614199,CR614594,CR614663,CR619476,CR623838,D10924,L06797,X71635 | CXCR4 | CCAAGTTCTTAGTTGCTGT |
| NM_001002009,NM_001002010,NM_016489,AF151067,AF312735,AK290118,AK314109,AL136716,AM393138,AM393316,AM393481,BC013292,BC015856,BC066914,BC071652,CR533518,CR597528,CR616013,DQ892843,EU176737 | NT5C3 | GTGCCTTGAGGAATACAGA |
| NM_030641,AK074645,AK095881,AL834420,AY014879,AY358210,BC038950,BC047864,CR456381,CU012939,CU013227 | APOL6 | CTGCTGGAAAGATTATCTA |
| NM_001077269,NM_003387,AK097901,AL832275,BC045584,BX640870 | WIPF1 | CTCTTCCCTTAATATCTGT |
| NM_019096,AB024574,AF168990,AJ420518,AL834331,BC020980,BC028347,BC032315,BC064968,CR604253,CR607185 | GTPBP2 | GGCCCAGAGTCTGAAGAAA |
| NM_006432,AK222474,AK298975,AK300879,AK311931,BC002532,CR595914,CR601885,CR605546,CR608935,CR609490,CR622486,CR624497,DQ891907,DQ895093,X67698 | NPC2 | CTGAATAAACTACCAGTGA |
| NM_001127713,NM_015915,NM_181598,AF131801,AF444143,AK223436,AK290185,AY032844,BC010708,CR599776 | ATL1 | CATGACCAATTGTCAATTA |
| NM_005465,NM_181690,AF085234,AF124141,AF135794,AJ245709,AK308052,AL117525,AM392791,AM392849,AY005799,BC020479,BC121154,AW960221 | AKT3 | CAAAGCCAAACACATTTAT |
| NM_004120,AK057968,AK226018,AK314325,BC073163,CR457062,CR599314,M55543 | GBP2 | GATTGAAGTGGAACGTATA |
| NM_002038,NM_022872,NM_022873,AK024814,AK314488,BC011601,BC015603,BN000257,BT006850,X02492 | IFI6 | CACCCACAAGTATCTCGAT |
| NM_020746,BC028962 | VISA | CAAACATTCTTCGTTCTTT |
| NM_016246,AF126781,AK307437,AY358430,BC006283,BC006294,BI602824 | HSD17B14 | CTGTCTTTATCCTCTGTGA |
| AF027866,BX641156 | SERPINB7 | GCCTTTCTTTGATAAGACA |
| NM_033064,AK027889,AK092309,AK125457,BC008736,BC026217 | ATCAY | CTCAGAAACTGGCTCTGAA |
| NM_014788,NM_033219,NM_033220,NM_033221,AF220130,AK055833,AK292825,CR594155,CR598301,D50919,DQ893293,DQ896623 | TRIM14 | GCTAATGCAGAGTCAAGTA |
| NM_198584,AK095314 | CA13 | GACACTAAGCATTAACTTA |
| NM_001037126,NM_021807,AB051486,AF380839,AK027688,AL831989,AM393428,BC026174,BC067263 | EXOC4 | CTGAACTTGTTGGATGAAA |
| NM_014873,D86960 | LPGAT1 | CTCTTAGCTGTAGTCTATA |
| NM_001130079,NM_020963,AB046851,AK023297,AK074174,AK226011,AK308874,AK310512,AL833353,AM393007,AM393678,BC002548,BC009312,BC025339,BX647787,DQ892188,DQ896845 | MOV10 | CTGAGTATCTTCATGGGAA |
| NM_021100,AK001470,AK056242 | NFS1 | CACTCCATCTGATTTGCAT |
| NM_001130079,NM_020963,AB046851,AK057353,AK074174,AK226011,AK308874,AL833353,AM393007,AM393678,BC002548,BC004499,BC009312,BC025339,BX647787,DQ892188,DQ896845 | MOV10 | CAGGAATACCGGGTCTTAA |
| NM_002443,NM_138634 | MSMB | CATTGATTTAATACACATT |
| NM_001571,AK292027,AK314421,BC071721,CR596241,CR608119,CR613061,CR615920,CR624964,Z56281,BM477213 | IRF3 | CCCTTCATTGTAGATCTGA |
| NM_001135651,NM_001135652,NM_002759,AK290655,AK313818,AY302136,BC007769,BC022314,BC040851,BC057805,BC093676,BC101475,CR600308,M35663,M85294 | EIF2AK2 | CTTCCTGACATGAAAGAAA |
| NM_002053,AB208912,CR591787,M55542,CD676970 | GBP1 | CTGTGTAACTAAACTGAAA |
| NM_005204,AK290320,AK313295,BC104833,BC113566,CR542284,CR619928,D14497,Z14138 | MAP3K8 | GACCATGTGTCAAGACAGT |
| NM_004159,NM_148919,BC001114,CR591946,CR595654,CR596244,CR606312,CR621259,CR621302 | PSMB8 | CGTGTTCTCTATTTGAACA |
| NM_001012509,NM_016180,AF172849,BC003597,BC064405,BQ677785 | SLC45A2 | GCCATAAGTGTCACCATGA |
| NM_020119,NM_024625,AK055851,AK292811,BC025308,BC033105,BC040956,BX571742,BX647974 | ZC3HAV1 | GAGCGGAATTTATGCAAAT |
| NM_006985,NM_178541,XM_001717235,XM_001720909,XM_001720910,XM_001723013,XM_001723016,XM_001732844,XM_001732845,AF132984,BC039707,BC094882,BC146422 | LOC23117, LOC339047, LOC729602, LOC729978, NPIP | CACTCTGCTTGGGTTATCA |
| NM_007315,NM_139266,AK096686,AK225853,AK292604,AK315002,BC002704,BT007241,CR749636,EU831980,EU832073,M97935,M97936,BG699050,CR998778 | STAT1 | CAAGCGTAATCTTCAGGAT |
| NM_002198,NM_008390,NM_012591,AB103081,AK314025,BC009483,BT019755,BT019756,CR541713,CR594837,DQ893269,DQ896599,X14454,AK152005,AK152104,AK152193,AK153514,AK155983,AK157347,BC003821,CT010234,BC076382 | IRF1, Irf1, Irf1 | CAGATTAATTCCAACCAAA |
| NM_002198,NM_008390,NM_012591,AB103081,AK314025,BC009483,BT019755,BT019756,CR541713,CR594837,DQ893269,DQ896599,X14454,AK152005,AK152104,AK152193,AK153514,AK155983,AK157347,BC003821,CT010234,BC076382 | IRF1, Irf1, Irf1 | CAGATTAATTCCAACCAAA |
| NM_080657,NM_021384,NM_138881,AF026941,AF442151,AK300085,AK312670,BC017969,CR598269,DQ891977,DQ895168,AF236064,AF442152,AK010906,AK036373,AK149900,AK150069,AK150347,AK150406,AK150425,AK150525,AK150536,AK150616,AK150684,AK150731,AK150767,AK150783,AK150788,AK150812,AK150835,AK151140,AK151185,AK151191,AK151206,AK151286,AK151308,AK151339,AK151389,AK151410,AK151412,AK151414,AK151476,AK151502,AK151538,AK151767,AK151770,AK151825,AK151833,AK151846,AK151906,AK151909,AK151940,AK151954,AK151971,AK152050,AK152065,AK152075,AK152235,AK152239,AK152241,AK152310,AK152354,AK152493,AK152600,AK152601,AK152631,AK152644,AK152664,AK152701,AK152796,AK152861,AK152880,AK152895,AK152934,AK152962,AK153003,AK153024,AK153062,AK153077,AK153078,AK153232,AK153234,AK153306,AK153318,AK153324,AK153350,AK153435,AK153453,AK153556,AK159147,BC057868,CT010378 | RSAD2, Rsad2, Rsad2 | ATGAAAGACTCCTACCTTA |
| NM_001729,AM392725,AM392995,AM393303,AM393550,BC011618,S55606 | BTC | CTATCAATGAAGATATTGA |
| NM_138287,AF484416,AK125086,AL833598,AY225123,BC042191,BC060509,BX648267,BX648645 | DTX3L | CTTACTTTGAATACTTTAA |

| | | |
|---|---|---|
| NM_001547,AK300431,AK312831,BC005987,BC032839,DA631671 | IFIT2 | CCCTGGAATGCTTACGTAA |
| NM_020414,AB209382,AF134475,AF145022,AF214731,AK025162,AK296726,AK297232,AK304504,AL136886,AM393538,BC008847,BC009406,BC096826,BX537533,CR596324,CR615921,CR619609,CR624535,DQ893523,DQ896510 | DDX24 | GGCTGTGGGAATTAATTAA |
| NM_003745,AB000676,AB005043,AK127621,U88326 | SOCS1 | ACCTCTTCATGTTTACATA |
| NM_001547,AK300431,BC032839 | IFIT2 | CCAAATCCTTCATGTAATA |
| NM_001003818,NM_001003819,AB039903,AK290172,BC047564,BC065575,BC136871,BG721109,CX760618 | TRIM6, TRIM6-TRIM34 | CGTGGTTGAGTTTAGATAA |
| NM_001007024,NM_001007025,NM_004871,AF047438,AF073926,AK000606,AK291381,AK298714,BC012620,BC040471,BE785872,BG654545 | GOSR1 | GAGACAATGGCGATTGAGA |
| NM_198282,AK095896,AK290661,FJ222241 | TMEM173 | CTGGCATGGTCATATTACA |
| NM_006877,BC008281,CR594139,CR610393,EU831974,EU832067,M24470 | GMPR | CACTCCATGTTTACAGCAA |
| NM_033109,AJ458465,AK307589,AY027528,AY290863,BC009057,BC021170,BC053660,CR625599 | PNPT1 | CAATAGGATTGGTCACCAA |
| NM_016118,AF155099,AF300717,AF459743,AY129295,BC046354,CR611732 | NUB1 | CTCTTTAAAGAGCTATATA |
| NM_006121,AK313986,BC063697,M10938 | KRT1 | GCCCTACTTTGAGTCATTC |
| NM_174908,NM_178335,AJ416916,AJ557013,BC065004,CR592179,CR601015,CR603155,CR610519,CR614299,CR618301,CR618316 | CCDC50 | GAGAAATCATCTTTGGACA |
| NM_018381,AK002148,AK096142,AK098062,AL137694,BC010847,BC026180,CR590846,CR592012,CR592313,CR594187 | C19orf66 | GAGTCTACCGTTCATTCCT |
| NM_001135651,NM_001135652,NM_002759,AK290655,AK313818,AY302136,BC007769,BC022314,BC040851,BC057805,BC093676,BC101475,CR600308,M35663,M85294 | EIF2AK2 | CCAGAAGGATTTCATTATA |
| NM_005858,AL050160,BC037270,Y11997 | AKAP8 | CCCAGAATATGCTGTAATC |
| NM_000389,NM_078467,AB209881,AK298901,AK309512,BC000275,BC000312,BC001935,BC013967,CR590203,CR612385,CR617250,CR621127,L25610,L26165,U03106,U09579 | CDKN1A | CAGCCTCTGGCATTAGAAT |
| NM_001165,NM_182962,AF070674,BC037420,L49432,U37546,U45878 | BIRC3 | CCTCTAGTGTTCTAGTTAA |
| NM_020651,AF300987,AF302505,AJ278859,AK027668,AK312441,AY364257,BC050019,BC050533,BC063611,BX649152 | PELI1 | GTTACAAGATGGCTCGTTA |
| NM_000389,NM_078467,AB209881,AB451290,AB451422,AK298901,AK309512,AM392711,AM392920,BC000275,BC000312,BC001935,BC013967,BT006719,CR536533,CR590203,CR612385,CR621127,L25610,L26165,L47232,L47233,S67388,U03106,U09579 | CDKN1A | CTGATCTTCTCCAAGAGGA |
| NM_138287,AF484416,AK125086,AL833598,AY225123,BC042191,BC060509,BX648267,BX648645 | DTX3L | GCACCATTGTGATTACTTA |
| NM_001644,BC096158,BC096159,BC096160,BC096161,BC101404,BC101405,BC101972,BC101973,L25877,L26234,U72891,U78720 | APOBEC1 | GGAATGCTCCCAGGCTATT |
| NM_018704,AB037854,AK022544,AK023340,AM393552,AM393771,BC016029 | CTTNBP2NL | CCTAATGAGCAATTGAAGA |
| NM_012420,AK293346,AK312358,BC025786,CR457031,CR624828,U34605 | IFIT5 | GAGCTGATTTCATCTGCTA |
| NM_005409,AF030514,BC005292,BC012532,BC110986,U66096 | CXCL11 | GCACATATTTCATAACCAA |
| NM_017878 | HRASLS2 | CAATTCTTACGGAGGAATA |
| NM_001135651,NM_001135652,NM_002759,AK290655,AK313818,AY302136,BC007769,BC022314,BC040851,BC057805,BC093676,BC101475,CR600308,M35663,M85294 | EIF2AK2 | CAGATACATCAGAGATAAA |
| NM_018704,AB037854,AK022544,AK023340,AM393552,AM393771,BC016029 | CTTNBP2NL | GCCATTGTGACCCAGAGAT |
| NM_007196,NM_144505,AB008390,AB008927,AB009849,AF095742,AY359036,BC040887 | KLK8 | GATGGCCCAGAGCAAGAAA |
| NM_005465,NM_181690,AF085234,AF124141,AF135794,AJ245709,AK308052,AL117525,AM392791,AM392849,AY005799,BC020479,BC121154 | AKT3 | CCAGAGGTGTTAGAAGATA |
| XM_001717586,XM_001718037,XM_001719484,XR_038342,AF035024,AF062180,BC020240,BX640874,Y13056 | EPC1, IGHM, IGHV3-30, LOC100132941, LOC100133739, LOC100134256, LOC652494, SCFV | GATTCACCTTCAGTAGCTA |
| NM_015900,AF035268,AF035269,AK222705,BC035347,U37591 | PLA1A | CATTCGTACTTAGGATTCA |
| NM_138456,AK092453,AK092761,AK096702,AK300917,BC012330,CR612726 | BATF2 | CCCAGGATTTCACAGTCGA |
| NM_018284,AK001823,AK309096,AL136680,BC017992,BC063819,BC140837,CR936755,CR936758,EU832447,EU832454,EU832530,EU832536 | GBP3 | CTGAAGCTAACGCAAGGTA |
| NM_017878,BC152862 | HRASLS2 | GAGACAACTACAGGGTCAA |
| NM_001165,NM_182962,AF070674,BC037420,L49432,U37546,U45878 | BIRC3 | CAGATTCGTTCAGAGTCTA |
| NM_002360,AF059194,AK092414,BC148265,BC152387 | MAFK | CAAACCGAATAAGGCATTA |
| NM_001815,BC106728,D90277,L00692,L00693,M76742 | CEACAM1, CEACAM3 | GACAGTTCCATGTATACCA |
| NM_152574,NM_027238,NM_001106665,AK093459,AK298581,AK298998,AK301185,AK301427,AK310271,AK310482,BC042417,AK037208,AK049108,AK132655,BC151021 | TTC39B, Ttc39b, Ttc39b | CCGAGTGTCTCCTACAGAA |
| NM_002360,AK056767,AL831913,BC012777,BC094871 | MAFK | CGTGGTAGGTAATCCATAT |
| NM_001040147,NM_003784,AF027866,AK290673,BX641156,D88575 | SERPINB7 | GTGTCTCACCCATTTCTAA |
| NM_005615,AK313580,BC020848 | RNASE6 | CTAGGATTCTCCTTCTGAA |
| NM_002818,NM_001029855,NM_011190,AK026580,AK225876,AY771595,BC004368,BC019885,BC072025,BX161498,CR541657,CR541743,CR594185,CR600073,CR601043,CR615548,CR618033,CR620148,D45248,DQ893404,DQ896722,AK012344,AK088654,AK143556,BC002301,BC005680,BC057859,D87910,U60329,AA216967,BY498670 | PSME2, Psme2, Psme2b-ps | GCCTTCTATGCTGAGCTTT |
| NM_000201,AF340038,AF340039,AK130659,AK298054,AK312636,BC015969,BT006854,CR617464,DQ890615,DQ893785,J03132,M24283,X06990 | ICAM1 | CAACCAATGTGCTATTCAA |
| NM_030641,AK074645,AK095881,AL834420,AY014879,AY358210,BC038950,BC047864 | APOL6 | CTTGGAGGTCCAAATAATA |
| NM_001560,AB209849,AK123515,AK296958,BC009960,BC015768,U81379,Y09328,Y10659 | IL13RA1 | CTCTCAGTGATGGAGATAA |
| NM_015900,AF035268,AF035269,AK222705,AK301880,BC035347,BC047703,BX647516,U37591 | PLA1A | CAGCTTATTGTAGACCATT |
| NM_001572,NM_004029,NM_004031,AF076494,AK303752,U53830,U53831,U53832,U73036 | IRF7 | GACATCGAGTGCTTCCTTA |
| NM_006820,AB000115,AK223087,AK225442,AK293385,AK303187,AK312272,AL832618,BC015932 | IFI44L | CCATTTATGTTGTGTGACA |
| NM_018381,AK002148,AK096142,AK098062,AL137694,BC010847,BC026180,BC035817,CR590846,CR592012,CR592313,CR594187 | C19orf66 | CATGTCCTCTCACCAGAAT |
| NM_005615,BC020848 | RNASE6 | CCTGTGTTCACAGCAATAA |
| NM_001037126,NM_021807,AB051486,AF132734,AF380839,AK027688,AL831989,AL834475,AM393428,BC020607,BC026174,BC067263 | EXOC4 | GAGTTACTGTTTGACAAGT |
| NM_005514,XR_037663,AB008102,AF130734,AF389378,AK124160,AK292226,AK300685,AY007140,BC007243,BC013187,BC013828,BC091497,BX647315,CR591388,CR591429,CR591706,CR592429,CR592731,CR594258,CR595307,CR597318,CR598082,CR598972,CR599061,CR599115,CR601192,CR601398,CR602366,CR606027,CR607623,CR607790,CR608195,CR608347,CR608524,CR609608,CR609850,CR611020,CR611400,CR611427,CR611658,CR612242,CR612477,CR612532,CR613710,CR618664,CR619303,CR619846,CR620408,CR621133,CR622087,CR622371,CR622879,D83043,D87665,L24373,L32862,L42345,M12678,M15470,M16102,M27540,M28203,M28204,M28205,M32317,M32319,M32320,M84382,U88407 | HLA-B, HNRPA1P5, MICA | GTGTCTCTCACAGCTTGAA |
| NM_014547,AF177171,AF237631,AK310344,AK312569,BX647277,CR611644 | TMOD3 | CAATTCTTGGGATGCACAA |

| | | |
|---|---|---|
| NM_006435,BC009696,BT009859,CR456894,CR541874,CR604902,DQ891667,DQ894615,X57351,BG164993 | IFITM2 | CAGCCTCCCAACTACGAGA |
| NM_001135651,NM_001135652,NM_002759,AK290655,AK313818,AY302136,BC007769,BC022314,BC040851,BC057805,BC093676,BC101475,CR600308,M35663,M85294 | EIF2AK2 | CAGGGAGTAGTACTTAAAT |
| NM_003113,XR_015185,XR_016060,XR_036905,XR_037908,XR_032216,XR_032636,XR_033528,XR_033584,XR_034140,XR_034184,XR_034482,XR_034492,XR_034517,XR_034623,XR_034778,XR_034873,XM_001053572,XR_005553,XR_005816,XR_006289,XR_007502,XR_008405,XR_008705,XR_008919,AF056322,CR616734,L79988,AK016539 | HMG1L6, LOC100132863, LOC729858, SP100, 100043354, 4932431P20Rik, EG666491, EG666786, LOC100046019, LOC100048350, OTTMUSG00000006164, OTTMUSG00000015039, OTTMUSG00000015121, LOC498043, LOC500210, LOC685764, LOC686375, RGD1562266, RGD1564581 | CATTATGAAAGAGAAATGA |
| NM_014426,NM_152227,XR_037381,XR_037574,XR_037582,AF121855,AK001793,AK054634,AK129505,AK223308,AK308017,BC000100,BC062638,BC093623,BC093980,CR590760,CR591673,CR600241,CR601419,CR610846 | LOC100131940, SNX5 | GGTCTATGACCAAAGAAGA |
| NM_004120,AK057968,AK226018,AK314325,BC073163,CR457062,CR599314,M55543 | GBP2 | CAAATCATTGGAGCCAATA |
| NM_020359,BC055415 | PLSCR2 | CATTGAATATTTACATGGA |
| NM_004613,NM_198951,AK058031,AK291714,AK295775,AK300292,AK314618,AY675221,BC003551,CR604340,M55153,M98478,S81734,BX471706 | TGM2 | CCAAGTTCATCAAGAACAT |
| NM_005409,AF030514,BC005292,BC012532,BC110986 | CXCL11 | GAAACATTCTTATGCATCA |
| NM_001032409,NM_002534,NM_016816,AJ629455,AK123528,AK223006,AK225384,AK291003,AY730627,AY730628,BC000562,BC061587,BC071981,BT006785,CR590221,CR603997,CR613331,CR621070,CR626748,D00068,DQ891892,DQ895079,X02661,X02874,X02875,X04371 | OAS1 | CTGAATTACCCATGCTTTA |
| NM_015668,AK097399,AK292559,AK302475,AK309194,AY009106,BC036665,BC047060,BC050053,BC092411 | RGS22 | CTGGATAGAAGATACGCTT |
| NM_002407,AF071219,AJ224173,BC062218 | SCGB2A1 | CAGTCACATAGAACTCTGA |
| NM_001035267,NM_021104,AF026844,AK023340,BC014383,BC015984,BC017820,BC032611,BC062462,BC064907,BC066328,BC070252,BC070253,BC070254,BC070255,BC070483,BC082759,BC105279,BC105286,BC105296,BC105794,BC105799,BC142666,BC142726,CR599786,CR605341,CR609127 | CTTNBP2NL, RPL41 | CATATAATAAATCACCTCT |
| NM_004509,NM_004510,NM_080424,AF280094,AF280095,AK026488,AK301097,AL832300,BC019059,CR625933,L22343 | SP110 | CAAAGAACTGGAAACGGAA |
| NM_004217,AB011446,AB011450,AF004022,AF015254,AK297976,BC000442,BC009751,BC013300,BC080581,CR591471,CR600836,CR603941,CR604968,CR605805,CR617702 | AURKB | GCCTGATGGTCCCTGTCAT |
| NM_004972,AF001362,AF005216,AF058925,AK292525,AK302618,AY973034,BC039695 | JAK2 | GTACAGATTTCGCAGATTT |
| NM_001560,NM_133990,AB209849,AK313467,BC009960,BC015768,CR603161,DQ893454,DQ896296,U81379,U81380,Y09328,Y10659,AK080147,AK082889,AK089984,AK143292,AK154675,BC021472,BC052425,BC059939,S80963 | IL13RA1, Il13ra1 | CAAATAATGGTCAAGGATA |
| NM_022168,AF095844,AK056293,AK292941,AK314611,AY017378,BC111750 | IFIH1 | GATATTAAAGAATGTAACA |
| NM_152542,AK314417,AL834271,AY157615,AY435431,AY994097,BC037552 | PPM1K | CTTGCAATGACAAGAAGTA |
| X98507 | MYO1C | GGCTGAATTATCGGTGATA |
| NM_020830,AB037856,AK022888,AK023415,BC040525,BC065934 | WDFY1 | GCAATATTCTGGCAATTTA |
| NM_022371,AJ299403,AK025998,AK130132,BC001085,BC007571,BC011746,BC018292,BC088368,CR612199,CR614363 | TOR3A | GTCTTGACATGTTTGTTGT |
| NM_004585,AB030815,AF060228,AF092922,AK293357,AK312110,BC009678,DQ891444,DQ894622,DA973023,R55646 | RARRES3 | CCCAAACCTGGAGACCTGA |
| NM_006074,AK290434,AK298715,AK298934,AK301192,AK310098,AL360134,AM040752,BC022281,BC035582,CR593076,DQ891201,DQ894384,X82200 | TRIM22 | CGCACCTGCACATTTAAGA |
| NM_014290,AB025254,AK294488,AK301954,AK310227,AK314853,BC028694,DQ894121 | TDRD7 | GATCGCACATGTTTATTTA |
| NM_005419,AK296939,AK308051,BC006092,BC051284,BX640607,CR621797,M97934 | STAT2 | CACCCTCCCTGTGGTGATT |
| NM_001080391,AF255565,AK090565,AK126529,AK160379,CR749288 | SP100 | GGAAACTCCACATCACAAT |
| NM_017912,AF336798,AK000644,AK097168,AK225506,AK295832,AL833664,AY653201,AY653202,AY653203,BC035775,BC042047,BX647121 | HERC6 | CATACATGCTTATGCATGA |
| NM_003810,AK296085,AK312742,BC009795,BC020220,BC032722,BT019564,CR456895,CR594331,CR598109,DQ848564,DQ890873,DQ894029,EU183231,U37518,U57059 | TNFSF10 | GGCATTGCTTGTTTCTTAA |
| NM_052966,AB050477,AF288391,AK022527,AK074139,AY550972,BC018532,BC030531 | FAM129A | CTGAGCAGGTGATTATTTC |
| NM_020414,AB209382,AF161446,AF214731,AK025162,AK294613,AK296726,AK297232,AL136886,AM393538,BC008847,BC009406,BC096826,BX537533,CR596324,CR615921,DQ893523,DQ896510 | DDX24 | GAACCAGTACCCAGAAAGA |
| NM_007196,NM_144505,NM_144506,NM_144507,AB008390,AB008927,AB009849,AF095742,AY359036,BC040887,CR601157,DQ267420 | KLK8, KLK9 | AGCACTAGATCTCCCTTAA |
| NM_022831,XM_001725626,XM_001726789,XM_930476,XR_015691,XR_017594,XR_038728,XR_038764,XR_039750,AK022868,AK289865,AL833718,BC015535,BC043142,BC067805 | AIDA, LOC646050, LOC646890, LOC653631 | CCAATTGTAATAGAACTAT |
| NM_001017369,NM_006745,AK292418,AK309206,BC010653,BC107879,CR605188,CR617900,CR623263,CR623543,DQ891060,DQ894237,U60205,U93162,BU940384,BX441001 | SC4MOL | ACCATTCGTTTATTAGAAA |
| NM_014426,NM_152227,NM_001106518,XR_006944,XR_008997,AF121855,AK001793,AK054634,AK129505,AK223308,AK308017,BC000100,BC062638,BC093623,BC093980,CR590760,CR591673,CR600241,CR601419,CR610846,BC160883 | SNX5, LOC293740, Snx5 | AGAAGAACTTCCTTATTAA |
| NM_001130020,NM_001130021,NM_005177,AK223554,BC017826,BC032398,BX648978,L78933,Z71460 | ATP6V0A1 | CTGGAACTGACCGAATTAA |
| NM_001141969,NM_001141970,NM_001350,NR_024517,AB209493,AF006041,AF015956,AF039136,AF050179,AF097742,AK223404,AK292187,AK303767,AK303854,BC000220,BC073776,BC109073,BC109074,CR457085,EU446949 | DAXX | CATCGTTACTGTCAGAAGA |
| NM_052941,NM_207398,NM_001039646,NM_001039647,NM_008620,NM_029509,NM_172777,NM_194336,XR_004768,XR_032536,XM_001064585,XM_223172,AF288814,AK096141,AK131094,AK312417,AL832576,BC008421,BC017889,BC050625,BC070055,BC156123,BC157015,AK088861,AK128993,AK152226,AK165231,AK171913,AK220567,BC007143,BC031475,BC057170,BC057969,BC112328,BC114209,BC115768,BK005759,DQ295175,DQ985742,EF494423,EF494424,EU304258 | GBP4, GBP7, 5830443L24Rik, 675363, BC057170, EG634650, Gbp10, Gbp4, Mpa2l, RGD1561523 | GAGTTTCTTTCCAGACTTT |
| NM_006820,AB000115,AK223087,AK225442,AK293385,AK303187,BC015932 | IFI44L | CAATGAGAGTCAATCTCTA |
| NM_017654,NM_152703,AB095925,AB095926,AF445355,AF474973,AK000080,AK097204,AK122951,AK125101,AK125131,AK225201,AK304307,AK304779,AL832264,BC029108,BC038974,BC127117,BC127118,BC132773,BC132775,BC150249,BX647072 | SAMD9, SAMD9L | GCAAGACATTTCTACATTA |
| NM_017554,AB033094,AK001770,AK026003,AK304269,AY134858,BX648758,DQ063584,DQ063585 | PARP14 | CACATCTTCACTCAAGATA |
| NM_020140,NM_152788,AF145204,AF164792,AK294994,BC142669 | ANKS1B | GTGGTCTAATCAACAGATA |
| NM_001032394,NM_001032395,NM_020455,NM_198569,AB183546,AB183547,AB183548,AB183549,AK075087,AK092519,AL080079,BC075798,BX648315 | GPR126 | CATTCACTTGCATTGTAAA |
| NM_016246,AF126781,AK307437,AY358430,BC006283,BC006294 | HSD17B14 | CCCTGGATGAAAGTCCATA |

| | | |
|---|---|---|
| NM_001136540,NM_001136541,NM_003661,NM_030882,NM_145343,NM_145637,AB208990,AF019225,AF305224,AF305225,AF305428,AF305429,AF323540,AF324223,AF324224,AK056938,AK296099,AK298053,AK300454,AK303919,AK303978,AK309143,AK313752,BC004395,BC017331,BC112943,BC113867,BC114475,BC127186,BC141823,BC142720,BC143038,BC143039,CR593100,CR626091,DQ893431,DQ896744,BG419709,DA375861,DC340792,DC367266 | APOL1, APOL2 | CAGTATCTTTATTGAGGAT |
| NM_003335,BC006378,BT007026,DQ893307,DQ893707,L13852 | UBA7 | CCCGGCATGAGTTTGAAGA |
| NM_016323,AB027289,AK093155,AK301756,AK302142,AY337518,BC140716 | HERC5 | CGTCATATGCCAGTTGGAT |
| NM_001142315,NM_001142316,NM_005574,NM_001142335,NM_001142336,NM_001142337,NM_008505,XM_001479884,AF257211,BC034041,BC035607,BC042426,BC073973,CR604507,CR614368,CR625714,DQ892333,DQ895538,X61118,AK013416,BC057880,M64360,BM690599 | LMO2, Lmo2, LOC100048263 | CCGCCTGTCAGAAGCATTT |
| NM_004223,NM_198183,AF061736,AK093462,AK129621,AK226165,BC032491,CR601863,CR609116 | UBE2L6 | CCAAGCCACTGATGGGAAT |
| NM_006074,AK290434,AK298715,AK298934,AK301192,AL360190,AM040752,BC022281,BC035582,DQ891201,DQ894384,X82200 | TRIM22 | CTGCTTATCCGTATTTCAA |
| NM_002462,AK093008,AK096355,AK225885,AK315465,BC014222,BC032602,CR592170,M30817,M33882 | MX1 | CTCATCACACATATCTGTA |
| NM_015459,NM_001044241,AK023383,AK097588,AK301910,AL117600,BC077727,CR936784 | ATL3, Atl3 | CCCTGACTTTGATGGGAAA |
| NM_024911,AK026744,AK074583,AK074984,BC137113,CR591378,CR602492,DQ323735 | GPR177 | CCCACTGAGTTAATATTTA |
| NM_001127397,NM_001127398,NM_015701,AF131849,AK001913,AK315477,AY358717,AY453410,BC013129,BC022228,CR597633,CR601764,CR604548,CR608194,CR613668,CR616366,CR623209,DQ894762 | C2orf30 | CATCTCCTGTGAATGACAT |
| NM_001008540,NM_003467,AF147204,AK129916,BC020968,CR594428,CR594588,CR596547,CR598681,CR598713,CR601301,CR604071,CR605131,CR610268,CR614199,CR614594,CR614663,CR619476,CR623838,D10924,L06797,M99293,X71635 | CXCR4 | CAGCTGTTTATGCATAGAT |
| NM_030641,AK074645,AK095881,AY014879,AY358210,BC038950,BC047864,CR456381,CU012939,CU013227 | APOL6 | GAAAGATTATCTATAATCT |
| NM_006432,AK298975,AK300879,AK311931,BC002532,CR601885,CR605546,CR608935,CR609490,CR622486,CR624497,DQ891907,DQ895093,X67698 | NPC2 | CCAGCAATATTCAGTCTAA |
| NM_001003819,NM_001003827,NM_021616,NM_130389,AB039902,AB039903,AF220143,AF220144,AK027876,AL583914,BC136871 | TRIM34, TRIM6-TRIM34 | GTTATAGGGTTACAGAATA |
| NM_017633,XM_135029,XM_918327,XM_925654,XM_994662,NM_001106844,AF350451,AJ420592,AK000044,AK056057,AK292109,AY740520,BC000683,BC007351,BX648876,AK135132,BC023892 | FAM46A, Fam46a, Fam46a | GCTAATGTCACTTGCTATT |
| NM_004458,NM_022977,NM_001033600,NM_019477,NM_207625,NM_053623,AB061713,AB061714,AF030555,AK292070,AK294915,AK307566,BC034959,DQ890835,DQ893990,Y12777,AB033885,AB033886,AB033887,AJ243502,AK054387,BC016416,BC058663,DR004263 | ACSL4, Acsl4, Acsl4 | CAGATACTCTGGATAAATT |
| NM_004458,NM_022977,NM_001033600,NM_019477,NM_207625,NM_053623,AB061713,AB061714,AF030555,AK292070,AK294915,AK307566,BC034959,DQ890835,DQ893990,Y12777,AB033885,AB033886,AB033887,AJ243502,AK054387,BC016416,BC058663,DR004263 | ACSL4, Acsl4, Acsl4 | CAGATACTCTGGATAAATT |
| NM_001040147,NM_003784,AF027866,BX641156,D88575 | SERPINB7 | CTGCTGCTTTCTAGAAATA |
| NM_001127213,NM_002346,AF085875,AK129615,AK223074,AK311983,BC003392,BC119708,BC119709,CR593133,CR593361,CR593965,CR594916,CR597237,CR597286,CR599019,CR599186,CR602572,CR603269,CR603365,CR606037,CR606634,CR606660,CR610259,CR620041,CR623247,U42376,U56145,Z68179,DB458093 | LY6E | CCAGAGCTTTCTGTGCAAT |
| NM_018295,AK001862,AK225055,AK301268,CR597549,CR598469,CR603717,CR608299,CR608312,CR613329,CR623230,CR625239 | TMEM140 | CAGCTGCTGTTCATGAGCA |
| NM_006074,AL360190,AM040752,BC035582,X82200 | TRIM22 | AAGAGATGCTTGTACATTA |
| NM_001024912,NM_001712,AL833584,J03858,X14831,X16354 | CEACAM1 | CAACCTTCTTGTCATTGAA |
| NM_016323,AB027289,AK093155,AK301756,AK302142,AY337518,BC140716 | HERC5 | GACACAAACTTAATTCCTA |
| NM_006432,AK222474,AK298975,BC002532,CR595914,CR601885,CR605546,CR608935,CR609490,CR622486,CR624497,X67698 | NPC2 | CAGCTCTGCTGCTTCAACA |
| NM_006877,BC008281,CR594139,CR610393,EU831974,EU832067,M24470 | GMPR | CTTCACGTTTCGAAATTCA |
| NM_018704,AB037854,AK022544,AM393771,BC016029 | CTTNBP2NL | GTAAGGATGTTGAGTTACT |
| NM_152542,AK054678,AK314417,AL834271,AY157615,AY435431,AY994097,BC037552 | PPM1K | GAATTAACTTCATGGTGAA |
| NM_005384,BC008197,CR599259,CR608714,CR614563,S79880,U26173 | NFIL3 | GCTGTATATATTGAACATT |
| NM_001223,NM_033292,NM_033293,NM_033294,AK223503,AK290114,AK290122,AK301037,AK310059,AK313516,AY660536,BC041689,BC062327,CR592117,M87507,U13697,U13698,U13699,X65019 | CASP1 | CAATCTTTAACATGTTGAA |
| NM_001003818,NM_001003819,NM_058166,AB039903,AF220030,AK027664,AK290172,AK293295,AK298301,BC065575,BC136871,CR624250,CR749260 | TRIM6, TRIM6-TRIM34 | AGGATCCAGACAGAGTTTA |
| NM_000600,BC015511,CR590965,CR626263,M14584,M18403,M29150,M54894,X04403,X04430,X04602 | IL6 | CTCAGATTGTTGTTGTTAA |
| NM_001003818,NM_001003819,NM_058166,AB039903,AF220030,AK027664,AK290172,AK293295,AK298301,BC047564,BC065575,BC136871,CR624250,CR749260 | TRIM6, TRIM6-TRIM34 | AGTTTAATCAGCTGCGAAA |
| NM_052957,AJ311392,AK097433,AK127607 | ACRC | CTGTGTATGTGCAGAAGTA |
| NM_020830,AB037856,AK022888,BC040525,BC065934 | WDFY1 | ATGTTAAGCTTCCTGTATT |
| NM_001511,BC011976,J03561,X12510 | CXCL1 | CAATCTGGATTCATATTTA |
| NM_000584,AK131067,BC013615,CR594973,CR595357,CR600500,CR601533,CR601902,CR603686,CR619554,CR623683,CR623827,M17017,M26383,Y00787 | IL8 | GGAATAATGAGTTAGAACT |
| NM_004696,AK129985,AK223307,AK290720,AK295946,AK298539,AK313735,AL833619,BC021664,BX647948 | SLC16A4 | GTTCTGCTTTCTTATACCA |
| NM_003528,AK093747,BC005827,BC069193,CR615793,M60756 | HIST2H2BE | CAAATGAGCTCTAGACATT |
| NM_020140,NM_152788,NM_181670,NM_001128086,XM_917405,XM_001076955,AF145204,AF164792,AK289768,AK294191,AK294994,AK295988,AM980945,AY281131,AY281132,AY283057,AY356353,AY620824,AY753193,BC026313,BC068451,BC091512,BC142669,BC150204,BC160005,AK032061,AK032211,AK034154,AK038609,AK049899,AK132228,AK163750,BC098373 | ANKS1B, Anks1b, Anks1b | GGCTAACTGTCAGAAGTCT |
| NM_007315,NM_139266,AK096686,AK292604,AK315002,BC002704,BT007241,CR618286,CR749636,EU831980,EU832073,M97935,M97936 | STAT1 | AGCTGTTACTCAAGAAGAT |
| NM_002308,NM_009587,NR_024043,AB003517,AB005894,AB006782,AB008492,AK126017,AK223232,AK296031,AK301624,BC105942,BC105944,BC110340,CR598297,CR602208,CR604285,CR608609,CR614567,CR616420,Z49107 | LGALS9 | GACTTCAGATCACTGTCAA |
| NM_020858,NM_024966,NM_153616,NM_153617,NM_153618,NM_153619,AB040912,AF389427,AF389428,AF389429,AF389430,AK290032,BC150253 | SEMA6D | GGATCAAGTTTATACAGTA |
| NM_021629,AF300648,AK001890,AK290571,AK297174,BC000873,CR591019 | GNB4 | GCTGGTTACGATGACTTTA |
| NM_002053,AB208912,CR591787,M55542,CB164272 | GBP1 | CACCCTAGCTTCTTAGTGA |
| NM_000577,NM_173841,NM_173842,NM_173843,AF043143,AK290898,AK290926,BC009745,BC068441,CR605915,CR616671,CR619093,CR625646,DQ892566,DQ895781,M55646,X52015,X53296,X84348,BI766516 | IL1RN | GACCAAATGTCAATTTAGA |
| NM_001511,BC011976,J03561,X12510 | CXCL1 | CTGCACACTGTCCTATTAT |
| NM_015668,AK097399,AK292559,AL117544,AY009106,BC036665,BC050053,BC092411 | RGS22 | CTGGTGCTACTCAAAGTAT |
| NM_004688,AB451305,AB451436,AK291548,BC001268,BC021987,CR604696,U32849 | NMI | GCATTCAAATGGATGAAGA |

| | | |
|---|---|---|
| NM_003895,NM_203446,AB020717,AF009039,AF009040,AK295706,AK304044,AK307762,BC098395,DQ421853 | SYNJ1 | CTGCCAGTCACCTACAATA |
| NM_003745,AB000676,AB005043,AK127621,U88326 | SOCS1 | CTCCTACCTCTTCATGTTT |
| NM_002162,AK226023,AK309643,BC034409,BC040123,BC046121,BC058903,CR623746,S50015,X69711,X69819 | ICAM3 | CTCAGATTCCGCACCAATA |
| NM_001710,AK130533,AK223400,AK304045,BC004143,BC007990,CR590153,CR590439,DQ892313,DQ895516,J00185,K01566,L15702,S67310,X72875 | C2, CFB | CTGATCAAGCTCAAGAATA |
| NM_001122665,NM_004660,XM_001713697,NM_010028,XR_032115,NM_001108246,XR_006413,AF000984,AF000985,AK302172,AK303176,AK303638,BC034942,AK140990,AK163250,L25126,Z38117,BC085914 | DDX3Y, LOC100130220, Ddx3x, LOC100045923, Ddx3x, LOC498906 | CACCTCATTCTTTAATGAA |
| NM_002164,AK313259,AY221100,BC027882,CR591703,CR613807,CR614485,CR616634,CR621546,M34455,X17668 | INDO | GAGGCACTGATTTAATGAA |
| NM_002198,AB209624,AK314025,BC009483,BT019755,BT019756,CR541713,CR594837,DQ893269,DQ896599,X14454 | IRF1 | CTCTGTCTATGGAGACTTT |
| NM_014426,NM_152227,AK001793,AK026227,AK129505,AK223308,AL832967,BC000100,BC062638,CR601419,CR608822,CR621801 | SNX5 | AGGTATATATGGAAACAAA |
| NM_001356,AB208983,AB451220,AB451343,AF000982,AF000983,AF061337,AK291153,AK296906,AK297159,AK304661,BC011819,U50553,BM833870 | DDX3X | GCGCTATATTCCTCCTCAT |
| NM_001775,AK297592,AK303169,CR606718,CR610722,D84276 | CD38 | GTAACATCCTTTCTATTGA |
| NM_007346,AF109134,AF172449,AF172450,AF172451,AF172452,AF172453,AK022234,AK024485,AK314696,BC008768,BC014137,BC032666,DQ892943,EU176723,BM551600 | OGFR | CCAAACCTGAGTTTCTACA |
| NM_152574,AK298581,AK298998,AK301185,AK310271,AK310482,BC070139,BC142985 | TTC39B | CCTTGGAAACCATCTCAAT |
| NM_000416,AF056979,AK222803,AK294252,AK297052,AK312674,BC005333,BT006814,CR601864,CR624388,CR625186,EF535103,J03143,DC349115 | IFNGR1 | CAAATCATGATTGACATAT |
| NM_001127397,NM_001127398,NM_015701,NM_001106023,AF131743,AF131849,AK001913,AK315477,AY358717,AY453410,BC013129,BC022228,CR597633,CR601764,CR604548,CR608194,CR613668,CR616366,CR623209,DQ894762,BC166988 | C2orf30, RGD1306508 | CTCATGCTGTTACTGTATA |
| NM_006985,NM_178541,XM_001132754,XM_001717235,XM_001717652,XM_001717676,XM_001720909,XM_001720910,XM_001723013,XM_001723016,XM_001723375,XM_001723497,XM_001724221,XM_001732844,XM_001732845,XR_016512,XR_016550,XR_041580,AF132984,AK128772,AK131084,AK131085,AK160377,AK302708,BC010188,BC039707,BC146422,CR619663 | LOC100132540, LOC100132620, LOC100133970, LOC23117, LOC339047, LOC399491, LOC642778, LOC642799, LOC728734, LOC728741, LOC729602, LOC729978, LOC730153, NPIP | GAAATTGGATATGAAAGAA |
| X67325 | IFI27 | CTGGGTGAAATATACCAAA |
| NM_006121,AK313986,BC063697 | KRT1 | CTCCTGGTGGCATACAAGA |
| NM_012420,U34605 | IFIT5 | AAGGTAAACTTATAACTTA |
| NM_001001330,AK289976,AY562241,BC010040,BC038724,BC057832 | REEP3 | CAAAGGAGCAAGTTTAATA |
| NM_004060,NM_199246,AK297190,AK312913,BC000196,BC007093,BT007134,BX538035,CR619041,D78341,DQ893765,EU176176,L49504,U47413,U53328,X77794,BM479568,BX644635 | CCNG1 | GGTTTCAGACTTGATGAGA |
| NR_000035,AF464191,AF464192,AF464193,AF464194,AF464195,AF486844,AF486845,AF486846,AF486847 | CASP12 | GATGTGAGATCTACTTGAA |
| BC001683,BC001847,BC019625,CR591065,CR592413,CR597001,CR598230,CR598799,CR601069,CR604510,CR608003,CR617904,CR620504,CR624884 | CHAC1 | CCATCCATAGCCCTGGGAA |
| NM_021181,AL713801,AL833025,AL834424,BC027867 | SLAMF7 | AGTCTGAAGTCACATTGTA |
| NM_017654,NM_152703,AB095925,AB095926,AF445355,AF474973,AK000080,AK097204,AK122951,AK125101,AK125131,AK225201,AK304307,AK304779,AL832264,BC029108,BC038974,BC127117,BC127118,BC132773,BC132775,BC150249,BX647072 | SAMD9, SAMD9L | CAAGACATTTCTACATTAA |
| NM_007237,BC038091,BC054890,BC105960,U36499,U36500,U63420 | SP140 | CTGTAAACTTGCTATACAA |
| AK311914,BC012472,DQ894624,Y12653 | UBD | CATCTTATGGCATTGACAA |
| NM_014314,AF038963,AK023661,AK125989,AK301678,AK315040,BC070029,BC132786,BC136610,BX647917 | DDX58 | GGATTAGCGACAAATTTAA |
| NM_014314,AF038963,AK023661,AK125989,AK301678,AK315040,BC070029,BC132786,BC136610,BX647917 | DDX58 | GGATTAGCGACAAATTTAA |
| NM_000566,BC032634 | FCGR1A | GAAGGAAATAGTACTTACA |
| NM_174975,AK131358,AK298079,AY158086,AY240872,BC069641,BC101001,BC101002,BC101003,BC101004,BU584962 | SEC14L3 | GCCCAACCCTGATGATTAT |
| NM_003528,AK093747,BC005827,BC069193,CR615793,M60756 | HIST2H2BE | CACATTATGGACACTTAAA |
| NM_001025107,NM_001111,NM_015840,NM_015841,AB209891,AK097241,BC038227,BX538232,BX640741,EU176459,EU176799,U10439,U18121,X79448,X79449,X98559 | ADAR | CACATCAAATGCCTCAAAT |
| NM_014788,NM_033219,NM_033220,NM_033221,AF220130,AK055833,AK292825,CR594155,CR598301,D50919,DQ893293,DQ896623 | TRIM14 | GAAATTCACTGAACTCAGA |
| NM_016246,AF126781,AK307437,AY358430,BC006283,BC006294 | HSD17B14 | CTCCCTGGAGCTGTCTTTA |
| NM_003733,NM_198213,AJ225089,BC117408,BC117410,CR609351 | OASL | CTCATCCTCTCGAAGAAGA |
| NM_004048,AB021288,AF072097,AK026463,AY007153,BC032589,BC064910,CR590254,CR591572,CR592576,CR592679,CR593405,CR593761,CR593874,CR594444,CR596238,CR596347,CR596444,CR596717,CR597808,CR599099,CR599510,CR599875,CR603043,CR603887,CR604110,CR604555,CR604705,CR604949,CR605353,CR606667,CR607236,CR608139,CR608197,CR610094,CR610959,CR612132,CR612519,CR613616,CR613782,CR613955,CR614402,CR614556,CR615574,CR615578,CR615729,CR616171,CR616437,CR618556,CR619071,CR619828,CR620994,CR621079,CR623073,CR624595,CR625152,CR626821,V00567 | B2M | GACATGATCTTCTTTATAA |
| NM_004696,AK129985,AK223307,AK290720,AK295946,AK298539,AK313735,AL833619,BC021664,BX647948 | SLC16A4 | CTGGTTATATGATTATACC |
| NM_019096,AB024574,AF168990,AJ420518,AK000430,AK290267,AL834331,BC020980,BC028347,BC064968,CR607185 | GTPBP2 | CTCAAAGTCTTTCTGAATA |
| NM_005746,AK292851,BC020691,BC072439,BC106046,CR593609,CR606862,U02020 | NAMPT | AAGATGATGTCTTTAATGA |
| NM_002163,AK313524,BC126247,CR599608,M91196,BG107396,BI835080 | IRF8 | GTTACGCTGTGCTTTGAAT |
| NM_001134486,NM_052942,AF288815,AF328727,AF430642,AF430643,AK090479,AK315064,AL832285,AY358953,BC031639,BC033761,DQ895650,EU176216 | GBP5 | CTGAAGAAGTTCTGCAGAA |
| NM_015474,AB013847,AB208944,AK027811,AK304187,AK304795,AK315169,BC036450,CR614403 | SAMHD1 | CCATCATCTTGGAATCCAA |
| NM_005204,AK290320,AK313295,BC104833,BC113566,CR542284,D14497,Z14138 | MAP3K8 | CGTGTAAACTGATCCCAGT |
| NM_003895,NM_203446,AB020717,AF009039,AF009040,AK307762,BC019630,BC098395 | SYNJ1 | GTGATACTATGTTACATTA |
| NM_001039916,NM_001039917,NM_001039918,NM_001039919,NM_001039920,NM_001135734,NM_133476,BC053361,U80738 | ZNF384 | GTCTCACTTTGCTTCCTAT |
| NM_000783,NM_057157,AF005418,AK027560,AK075374,CR598753,CR604765 | CYP26A1 | GAAATCTGATGAGCTTGAA |
| NM_000584,AK131067,BC013615,CR594973,CR600500,CR601533,CR601902,CR603686,CR619554,CR623683,CR623827,M17017,M26383,Y00787 | IL8 | GGAGAATATACAAATAGCA |
| NM_002800,NM_148954,AK303118,BC065513,BX641100,CR541656,CR621193,S75169,U01025,X62741,BI832878 | PSMB9 | GTGAGAAATATCAGCTATA |
| NM_000386,AF091082,AK223403,AK301948,AK312896,BC003616,BT007018,X92106 | BLMH | CAGCCCATTGACTTCCTGA |

| | | |
|---|---|---|
| NM_006330,NM_013006,AF052112,AF077198,AF077199,AF081281,AF291053,AK289400,AK296257,AK298206,AK307277,AK313739,BC008652,BC010397,CR457103,DQ891527,DQ894717,BC085750,U97146 | LYPLA1, Lypla1 | CATTCCTTCTAACAGAATT |
| NM_002985,BC008600 | CCL5 | CACACAGCAGCAGTTACAA |
| NM_000029,AK222798,AK222815,CR592799,CR601683,CR625369,K02215 | AGT | CGTGTAGTGTCTGTAATAC |
| NM_080657,AF442151,AK300085,AK312670,BC017969,CR598269,DQ891977,DQ895168 | RSAD2 | GAGTCGCTTTCAAGATAAA |
| XM_001723375,XM_001724221,AK294177,AK308629 | LOC100132540, LOC100132620, LOC23117, NPIP | GAGGTTCCGTATTTGTAAT |
| D86960 | LPGAT1 | GGTAGATAATCTGGTAATA |
| NM_002407,AF071219,AJ224173,BC062218 | SCGB2A1 | GACAGCATTTGGTGTAATA |
| NM_006877,BC008281,CR594139,CR610393,EU831974,EU832067,M24470 | GMPR | ATGTTTACAGCAATTCATA |
| NM_002463,AK122952,AY312582,BC035293,DQ892116,DQ895311,M33883 | MX2 | GACAAGATGTTCTTTCTAA |
| NM_022831,XM_001725626,XM_001726789,XM_930476,NM_181732,XM_001473773,XM_905339,NM_001127600,AK022868,AK289865,AL833718,BC015535,BC067805,AK076031,AK081494,AK143716,AK149788,AK150639,AK150702,AK151783,AK152961,AK153448,AK158103,AK159318,AK167520,AK172451,BC057183,BC066829,BC086763,DA483113 | AIDA, LOC653631, Aida, LOC631071, LOC682999 | GATAGACGAGTATCAGATA |
| NM_020746,BC028962 | VISA | GTTCTTTAAGGCCTAACTT |
| NM_032206,AF389420,AK074133,AK090439,BC040278,BC050527,BC063566,BC156512,EF452236 | NLRC5 | CTCAGGCAGTGTCACTGAA |
| NM_021629,AK001890,AK022599,CR591019 | GNB4 | GATTGTCCCTGCATCTGAA |
| NM_001136540,NM_001136541,NM_003661,NM_145343,AF019225,AF305224,AF323540,AK298053,AK300454,AK303919,AK309143,AK313752,BC017331,BC112943,BC113867,BC114475,BC127186,BC141823,BC142720,BC143038,BC143039,CR626091 | APOL1 | CTAAACATTCTCAACAATA |
| NM_004217,AB011446,AB011450,AF004022,AF008552,AF015254,AK297976,AK314889,AY677083,BC000442,BC009751,BC013300,BC080581,BT019534,CR600836,CR603941,CR604968,CR605805 | AURKB | GCCAGAAGGTGATGGAGAA |
| NM_138287,AF484416,AK125086,AL833598,AY225123,BC042191,BC060509,BX648267,BX648645 | DTX3L | CAAGTTTGCTGATGACTTT |
| NM_005746,AK292851,BC020691,BC072439,BC106046,CR593609,CR606862,U02020 | NAMPT | GGCATCTTCCAATAGAAAT |
| NM_024586,NM_148904,NM_148905,NM_148906,NM_148907,NM_148908,NM_148909,NM_001134791,NM_133885,NM_173350,AF392445,AK022554,AK027535,AK027707,AK056617,AK091703,AK124037,AK128043,AK315167,AY178997,BC025978,CR457288,CR607539,CR609523,AK035658,AK131984,AK172010,BC021507,BC023759,BC026927,BC089163 | OSBPL9, Osbpl9 | GTAGTTCTTCCAACGTTTA |
| NM_000304,NM_153321,NM_153322,AK300690,BC019040,BC091499,CR592810,CR593633,CR597127,CR597311,CR598725,CR602788,CR602976,CR603134,CR604376,CR604692,CR604834,CR608530,CR610349,CR611493,CR611595,CR612069,CR615033,CR615858,CR618437,CR618628,CR618649,CR622251,CR624540,D11428,L03203 | PMP22 | CAAACTGGAGCCCTTGCAA |
| NM_004871,AF073926,AK000606,AK291381,BC012620,BC040471 | GOSR1 | CAGGTATAGTTCTGATACA |
| NM_022347,AK289793,BC032790,BC096704,BC098170,BC098309,BC098348 | IFRG15 | CAGGTATTCTTAGACCGTA |
| NM_152784,AK128088,BC043005 | TMEM146 | GAGCATGTCTTTAGGTATA |
| NM_001080391,AF255565,AK090565,AK126529,AK160379,CR749288 | SP100 | GTGCCTGTGGTCCCACTAA |
| NM_005082,AB208994,AK310720,BC016924,BC042541,CR617665,D21205,CF121587 | TRIM25 | GGAACAGTTAGTGGATTTA |
| NM_005531,AB208989,AF208043,AK296228,AK307957,AY138863,BC017059,DQ891062,DQ894242,M63838 | IFI16 | CATATCAGATTATTTGGAA |
| NM_014788,NM_033219,NM_033220,NM_033221,AF220130,AK055833,AK097480,AK292825,CR594155,CR598301,D50919,DQ893293,DQ896623 | TRIM14 | CCAAGAAATTCATTGATAA |
| NM_021964,AF039019,AF432210,AJ236885,AK314203,AL833616,BC050260,BC128582,L04282 | ZNF148 | CCTTAACTTTGTGACTGAT |
| NM_001003818,NM_001003819,AB039903,AK290172,BC047564,BC065575,BC136871,BG721109,CX760618 | TRIM6, TRIM6-TRIM34 | CGCGCTCTGTTCCTTAAGA |
| NM_022750,AL136766,AL137255,BC044660,CR602813 | PARP12 | CTGCCTACCATTTATAGAA |
| NM_001007245,NM_001550,AK308066,AK313211,BC001272,BT019889,BT019890,CR602983,CR614885,CR616001,Y10313 | IFRD1 | CAATCAATGAAGTGAAGAA |
| NM_005217,BC027917,BC119706,DQ890545,DQ890546,DQ896798,EU176174,M21131,M23281,X13621 | DEFA1, DEFA3 | GACTGCTATTGCAGAATAC |
| NM_005746,NM_021524,NM_177928,AK292851,BC020691,BC072439,BC106046,CR593609,CR606862,U02020,AK086415,AK153926,AK166270,AY679720,BC004059,BC018358,AB081730,AY831728 | NAMPT, Nampt, Nampt | AGTACATTCTTAATAAGTA |
| NM_020858,NM_024966,NM_153616,NM_153617,NM_153618,NM_153619,AB040912,AF389427,AF389428,AF389429,AF389430,AK021660,AK290032,BC150253 | SEMA6D | CACACTTTCTTCATGCCAT |
| NM_001135651,NM_001135652,NM_002759,NM_019335,XM_579433,AK290655,AK313818,AY302136,BC007769,BC022314,BC040851,BC057805,BC093676,BC101475,CR600308,M35663,M85294 | EIF2AK2, Eif2ak2 | GTTTACATTTCAAGTTATA |
| NM_022091,AK023189,AK315197,BC050681,BC125211,BC125212,BC130038 | ASCC3 | AGCAGAGCTTCTGACAGAT |
| NM_021105,AB006746,AF098642,AK313377,BC017901,BC021100,BC032718,CR593697,CR607459,CR623341,DQ890879,DQ894033,DR005513 | PLSCR1 | CAGTGTATAATCAGCCAGT |
| NM_006074,AK298715,AK298934,AK301192,AL360190,AM040752,BC022281,BC035582,DQ891201,DQ894384,X82200 | TRIM22 | GTTATAGGATTACAGAATA |
| NM_012429,AB033012,AK056163,AK091402,AL096881,AY203959,BC048337 | SEC14L2 | CTGCCTCTTTGATTACTAA |
| NM_002462,AK093008,AK096355,AK225885,AK315465,BC014222,BC032602,CR592170,M30817,M33882 | MX1 | CTGCCAGGCTTTGTGAATT |
| NM_004223,NM_198183,AF061736,AK093462,AK129621,AK226165,BC032491,CR601863,CR609116 | UBE2L6 | GCCTCAGACTGTGAAGTAT |
| NM_001080391,NM_003113,AF056322,AF255565,AK091898,AK293373,AK304205,BC011562,CR601757,CR613678,M60618,U36501,DA562355 | SP100 | GCTGCTCTATGACATTGTA |
| NM_006084,BC035716,CR596057,CR602478,CR603020,CR609755,CR610511,CR619285,M87503,BI497082 | IRF9 | GTGATAATCGTGTCCTGAA |
| NM_004060,NM_199246,BC000196,BC007093,CR619041,U53328,X77794 | CCNG1 | GGTTAACATTCTAGGCAGT |
| NM_006401,XR_017650,XR_018651,NM_130889,XM_001479982,XR_032654,AK223454,AK313733,BC013003,BC019658,CR595726,CR616535,U70439,Y07569,Y07570,Y07969,AB025582,AK011134,AK043419,AK136648,AK165807,BC003489,BC005628,BC093506,AW296974 | ANP32B, LOC646791, Anp32b, OTTMUSG00000010553 | GAGAACTTGTCTTGGACAA |
| NM_001098478,NM_018950,AK225185,BC062991 | HLA-F | CTCACGAACATACATAAAT |
| NM_174908,NM_178335,NM_001025615,NM_026202,AJ416916,AJ557013,BC065004,CR592179,CR601015,CR603155,CR610519,CR614299,CR618301,CR618316,AJ534985,AK012190,AK016827,AK077557,AK147820,AK151672,AK162434,AK165214,BC145933,BC145935,BY716501 | CCDC50, Ccdc50 | GACTTCTAATGGCTGAAGA |
| NM_001031683,NM_001549,AF026939,AF083470,AK290427,AK297137,AK312675,BC001383,BC004977,BT007284,CR618361,CR626380,U52513 | IFIT3 | CAAATTATTGGTATCTTCA |
| NM_000029,AB209769,AK222798,AK222815,AK293507,AK303755,AK307978,AK312291,BC011519,BT006851,CR591984,CR592799,CR595628,CR601683,CR606672,CR607896,CR607898,CR615493,CR615499,CR618295,CR618811,CR618903,CR619790,CR625369,CR625415,DQ891016,DQ894196,K02215,W01494 | AGT | CCCTCAACTGGATGAAGAA |

| | | |
|---|---|---|
| NM_004335,AK223124,AK291099,AK303593,BC033873,CR591045,CR609450,D28137 | BST2 | ATCACTACATTAAACCATA |
| NM_031458,AF307339,AK292959,AK313494,BC017463,BC039580,BX648869,CR596753 | PARP9 | GAGCAGCAGCTTACAATGA |
| NM_003641,AK290480,BC000897,BT007173,DQ891094,DQ894276,J04164,X84958 | IFITM1 | CATATTATGTTACAGATAA |
| NM_002198,NM_008390,NM_012591,AB103081,AK314025,BC009483,BT019755,BT019756,CR541713,CR594837,DQ893269,DQ896599,X14454,AK152005,AK152104,AK152193,AK153514,AK155983,AK157347,BC003821,CT010234,BC076382 | IRF1, Irf1, Irf1 | CTGGCTAGAGATGCAGATT |
| NM_002198,NM_008390,NM_012591,AB103081,AK314025,BC009483,BT019755,BT019756,CR541713,CR594837,DQ893269,DQ896599,X14454,AK152005,AK152104,AK152193,AK153514,AK155983,AK157347,BC003821,CT010234,BC076382 | IRF1, Irf1, Irf1 | CTGGCTAGAGATGCAGATT |
| NM_001003818,NM_001003819,AK290172,BC047564,BC136871,BG721109,CX760618 | TRIM6, TRIM6-TRIM34 | CTGCCTTTCTCGGAACGGA |
| NM_004116,NM_054033,BC002614,BC050998,CR614690,CR616275,CR626535,D38037,L37086,S69800,S69815 | FKBP1B | CACTTTCTCTCTTATAAAT |
| NM_001003818,NM_001003819,AB039903,AK290172,BC047564,BC065575,BC136871,BG721109,CX760618 | TRIM6, TRIM6-TRIM34 | CGCTCTGTTCCTTAAGATT |
| NM_006121,AK313986,BC063697,M10938 | KRT1 | GATAGTGTGAGAAATTCAA |
| NM_001012509,NM_016180,NM_053077,NM_001107653,AF172849,BC003597,BC064405,AF360357,AK029155,AK089932,AK148207,AY034377,BC125376,BC132429,BQ677785 | SLC45A2, Slc45a2, Slc45a2 | CCCATCAAAGCCTACTTAT |
| NM_003265,BC043204,BC068487,BC094737,BC096333,BC096334,BC096335,DQ445682,U88879 | TLR3 | GAGAAACTTTCTCAATTTA |
| X67325 | IFI27 | CAAATATACCTGGGTGAAA |
| NM_024119,AK021416,AK097669,AK225549,BC014949,BM723672 | DHX58 | CTTGTACGCACATAGACAT |
| NM_001077500,NM_002776,NM_145888,AF024605,AK292365,AK309456,AY561635,BC002710,CR450327,DQ890750,DQ893917 | KLK10 | AAATACATGTCCTGGATCA |
| AY562241 | REEP3 | CAGTAAAGGTAATTGTTCA |
| NM_000304,NM_153321,NM_153322,NM_008885,NM_017037,BC019040,BC091499,CR592810,CR593633,CR597127,CR597311,CR598725,CR602788,CR602976,CR603134,CR604376,CR604692,CR604834,CR605458,CR608530,CR610349,CR611493,CR611595,CR612069,CR615033,CR615858,CR618437,CR618628,CR618649,CR622251,CR624540,D11428,L03203,M94048,AK076005,AK171745,BC010765,M32240,M69139,S55427 | PMP22, Pmp22, Pmp22 | CCACCAACTGTAGATGTAT |
| NM_002122,XM_001719804,XM_001722107,XM_001722240,XM_001723439,AB209628,AF533895,AF533896,AF533897,AF533898,AF533899,AF533900,AF533902,AF533903,AF533904,AF533905,AF533906,AF533907,AF533909,AF533910,AF533911,AF533912,AF533913,AF533915,AF533916,AF533917,AF533918,AF533921,AF533922,AF533923,AF533924,AF533925,AF533926,AF533927,AF533928,AF533936,AF533937,AF533938,AF533939,AF533940,AK130598,AK130811,AK130838,BC008585,BC051832,BC125044,BC125045,M17846,M20431,M20506,M26041,M33906,M34996,X00033,X00370,X00452 | HLA-DQA1, LOC100133678, LOC731682 | GCTACCTAATTCCTCAGTA |
| NM_000389,NM_078467,AB209881,AK298901,AK309512,BC000275,BC000312,BC001935,BC013967,CR590203,CR612385,CR617250,CR621127,L25610,L26165,U03106,U09579 | CDKN1A | CCTCTGGCATTAGAATTAT |
| NM_001734,NM_201442,AK025309,AK290295,AM886411,BC056903,J04080,M18767 | C1S | CAGAGAACCCTTTGAAGAA |
| NM_001048205,NM_005132,AK001435,AK291191,AL832973,BC004159,BC010887,BC019326,CR596261,CR597987,CR609880 | REC8 | CTGCTTACCTCATTTCTGA |
| NM_020359,BC055415,BC069785,BC120969,BC141969 | PLSCR2 | CTAGAGACCTTGATGTTAA |
| NM_002116,NM_005514,NR_001434,XM_001713644,XM_001717045,AB008102,AB209117,AF008933,AF012766,AF015930,AF016641,AF036921,AF116214,AF130734,AF140506,AF165065,AF168611,AF170577,AF189017,AF196489,AF217561,AF287958,AF287959,AF298582,AF305699,AF405691,AF436090,AF436091,AF436092,AF436093,AF436094,AF436095,AF436096,AF436098,AJ001269,AJ001274,AJ002675,AJ003063,AJ005199,AJ010748,AJ131118,AJ223602,AJ249241,AJ290394,AJ537578,AJ580771,AK125608,AK130165,AK299998,AK300685,AK301014,AK301019,AK301282,AK304472,AK308374,AK310586,AK313119,AK313911,AY007140,AY102648,AY191309,AY191310,AY365426,AY569160,AY786585,AY786586,AY786587,AY856830,BC002463,BC003069,BC004489,BC007243,BC008457,BC008611,BC010542,BC013187,BC019236,BC041078,BX647315,CR590089,CR590229,CR590761,CR591238,CR592438,CR593473,CR593532,CR593771,CR596197,CR596736,CR597883,CR598559,CR598869,CR600122,CR600696,CR601516,CR603931,CR604944,CR605111,CR605534,CR605752,CR605841,CR608593,CR609975,CR610110,CR611177,CR611213,CR611534,CR612012,CR616547,CR619034,CR619049,CR619724,CR621304,CR622028,CR622132,CR624725,CR625317,CR625955,D14350,D14351,D14354,D16841,D16842,D16843,D26550,D32129,D32130,D32131,D38525,D50068,D50290,D50291,D50292,D50293,D50294,D50295,D50296,D50853,D50854,D64146,D64148,D64150,D64151,D64152,D83029,D83030,D83515,D83741,D87665,DQ327720,DQ327721,DQ327722,DQ336693,DQ336694,DQ336695,DQ336696,DQ336697,DQ336698,DQ891760,DQ891896,DQ895082,EU176622,L07950,L11603,L11604,L11666,L15005,L18898,L19937,L22027,L32862,L33922,L36591,L37880,L37881,L38251,L41925,L42345,L42545,L47338,L47984,L47985,L47986,L48400,L49341,L49342,L49343,L63544,L76252,L76639,L76930,L76932,L76933,L76936,L77114,L79935,L79936,M15470,M16102,M24031,M24033,M24042,M24095,M24097,M26430,M26712,M28203,M28205,M28206,M32317,M32318,M32319,M32321,M64740,M75138,M77774,M77778,M83191,M83192,M83193,M83194,M83195,M84172,M84173,M84174,M84377,M84378,M84379,M84382,M84386,M86404,M98453,M99389,M99390,U01848,U03027,U03697,U03754,U03859,U03907,U04245,U06695,U06696,U06697,U06862,U07161,U09864,U09912,U11262,U11264,U11265,U11266,U11268,U11269,U16298,U16309,U17107,U18660,U18930,U21052,U21053,U25971,U29057,U29880,U30904,U30936,U31334,U32184,U32661,U32678,U34618,U34619,U34810,U35431,U36392,U36914,U39088,U40498,U41057,U41386,U41420,U43334,U43335,U43336,U45480,U50574,U50710,U52429,U56825,U57028,U57966,U58110,U62824,U64801,U70528,U70529,U80945,X13111,X13112,X57954,X58536,X60764,X61700,X61703,X61704,X61708,X61709,X61711,X64366,X64454,X70856,X75953,X76189,X82122,X90390,X90391,X91399,X91749,X94566,X94567,X94568,X94570,X94572,X94573,X96638,X96724,X97802,X98742,X99704,Y08275,Y08994,Y10520,Y11441,Y11843,Y13028,Y13029,Y13267,Y13567,Y17224,Z15143,Z15144,Z23071,Z27120,Z46633,Z46808,Z83247,DC345094 | HLA-A, HLA-A29.1, HLA-B, HLA-C, HLA-H, LOC100133382 | CACTCCATGAGGTATTTCT |
| NM_004060,NM_199246,AK297190,BC000196,BC007093,BX538035,CR619041,D78341,L49504,U47413,U53328,X77794 | CCNG1 | CTTGTTCTATGGATTCCAT |
| NM_004048,AB021288,AF072097,AK303560,AK315776,AY007153,AY187687,BC032589,BC064910,CR457066,CR590254,CR592576,CR592679,CR593405,CR593761,CR593874,CR594444,CR596238,CR596347,CR596444,CR596717,CR597808,CR599099,CR599510,CR603043,CR603887,CR604555,CR604705,CR604949,CR605353,CR606667,CR607236,CR608139,CR608197,CR610094,CR612132,CR613616,CR613782,CR614402,CR614556,CR615574,CR615578,CR615729,CR616171,CR616437,CR619071,CR619828,CR620994,CR621079,CR623073,CR624595,CR625152,CR626821,DQ839493,DQ890551,DQ893711,S54761,S82297,S82300,V00567,X07621,BU658737 | B2M | CAGAGAATGGAAAGTCAAA |
| NM_001002009,NM_001002010,NM_016489,NM_026004,NM_001107862,AF151067,AF312735,AK290118,AK314109,AL136716,AM393138,AM393316,AM393481,BC013292,BC015856,BC066914,BC071652,CR533518,CR597528,DQ892843,EU176737,AK011525,AK011894,AK160139,AK208033,BC038029,BC166444,BF790086,CB127255,CB127877 | NT5C3, Nt5c3, Nt5c3 | CTGCCAAACTTCAGATAAT |
| NM_001048205,NM_005132,AF006264,AK001435,AK291191,AL832973,BC004159,BC010887,BC019326,CR457263,CR596261,CR597987,CR609880 | REC8 | AAGAAGCTATCTTGTTAGA |
| NM_001040147,NM_003784,AF027866,AK290673,AK290876,AK300828,BC069417,BC069442,BC069547,BC106743,BC106744,D88575 | SERPINB7 | CATAAATTGCCATTTCAAA |
| NM_001624,BC156646,U83115 | AIM1 | CTGGAAAGGTAGTGATATA |

| | | |
|---|---|---|
| NM_032505,AB058745,AK096640,BC117487 | KBTBD8 | CTAAATAAATGGACTCGTA |
| NM_005858,AL050160 | AKAP8 | GATTCCAGGTGAATATTTA |
| NM_004972,AF001362,AF005216,AF058925,AK292525,AY973034 | JAK2 | CCAATTCTATGAAGCAAAT |
| NM_001128166,NM_001128167,NM_001128168,NM_001128172,NM_001128173,NM_002578,NM_008778,NM_019210,AB102659,AF068864,AF155651,AK128022,AK290504,AK315284,AM943850,AM943851,AM943852,BC117353,BC117355,BC152761,AF082297,AJ496262,AJ496263,AK031853,AK158893,AM943853,AM943854,BC053403,U39738 | PAK3, Pak3, Pak3 | CAAAGTAAACGAAGCACTA |
| NM_006573,AF116456,AF132600,AF134715,AF136293,AF186114,AK309629,AY129225,AY302751,AY358881,BC020674,CR541818,DQ891413,EU176634,BI818459 | TNFSF13B | CGCCTTACTTCTTGCCTTA |
| NM_004079,AK024855 | CTSS | CTAAATTAACCTAAACGTA |
| NM_001032409,NM_002534,NM_016816,AJ629455,AK123528,AK223006,AK225384,AK291003,AK300346,AK301498,AY730627,AY730628,BC000562,BC061587,BC071981,BT006785,CR590221,CR603997,CR613331,CR621070,CR626748,D00068,DQ891892,DQ895079,X02661,X02874,X04371 | OAS1 | CTGAGAAGGCAGCTCACGA |
| NM_052941,AF288814,AK131094,AK312417,AL832576,BC008421,BC017889,BC050625,BC070055 | GBP4 | GCCTACAAATGACAAGCAA |
| NM_003641,AK290480,BC000897,BT007173,DQ891094,DQ894276,J04164 | IFITM1 | CTGTGACAGTCTACCATAT |
| NM_003745,AB005043,AK127621,U88326 | SOCS1 | CAGCTTAACTGTATCTGGA |
| NM_001135651,NM_001135652,NM_002759,AK290655,AK313818,AY302136,BC007769,BC022314,BC040851,BC057805,BC093676,BC101475,CR600308,M35663,M85294 | EIF2AK2 | GCAGTTAGTCCTTTATTAT |
| NM_006820,AB000115,AK223087,AK225442,AK312272,BC015932 | IFI44L | GGCTTACATTGATTACAAT |
| NM_000270,AK098544,AK126154,AK313490,BC104206,BC104207,CR407607,CR592961,CR594642,CR595160,CR598701,CR598851,CR598982,CR599285,CR600826,CR601251,CR601413,CR604278,CR604317,CR606974,CR608191,CR608198,CR608316,CR612303,CR612939,CR614792,CR624654,CR625788,X00737 | NP | GCAAACAAGCTGCACAGAA |
| NM_006019,NM_006053,AF025374,AY548968,BC018133,BC032465,CR594613,CR602058,EU176593,U45285 | TCIRG1 | CCGCTACCTGCTCCTGCTT |
| NM_006674,AK093953,AK290875,BC106759,BC114462,CR626077,L06175 | HCP5 | CAGATTACAATTACAATCA |
| NM_002800,NM_148954,BC065513,CR541656,CR621193,S75169,U01025,X62741,BI832878 | PSMB9 | CGCTTCACCACAGACGCTA |
| NM_005465,NM_181690,NM_011785,AF085234,AF124141,AF135794,AJ245709,AK308052,AL117525,AM392791,AM392849,AY005799,BC020479,BC121154,AF124142,BC066861 | AKT3, Akt3 | GTCATTATTGCAAAGGATG |
| NM_015900,AF035268,AF035269,AK222705,AK301880,AK313519,BC035347,BC047703,BX647516,U37591 | PLA1A | CTGAAGATAGCCTGTGTGT |
| NM_005101,M13755,BM712238,BU161226 | ISG15 | AGCATCCTGGTGAGGAATA |
| NM_022347,AK289793,BC096704,BC098170,BC098309,BC098348 | IFRG15 | CCTTGTTCTTCCTTGGTCT |
| NM_001001438,NM_002340,AK226141,AK296313,AK312489,BC035638,D63807,DQ891234,DQ894418,S81221,U22526,X87809 | LSS | GCGCTTAAGTATTTCCACA |
| NM_017912,AF336798,AK000644,AK097168,AK225506,AK295832,AL833664,AY653201,AY653202,AY653203,BC035775,BC042047,BX647121 | HERC6 | CATGCAAGAGGCATACAGA |
| NM_002053,AB208912,CR591787,M55542,CD676970 | GBP1 | CCAAATACACAATGTATTT |
| NM_021100,NM_053462,AF097025,AJ010952,AK001265,AK001470,AK056242,AK223239,AK297969,AK302023,BC065560,CR594451,CR610756,AF336041,BC072482,BC089205 | NFS1, Nfs1 | CACTTGATGTCAATGACAT |
| NM_012429,AB033012,AK056163,AK091402,AK223587,AL096881,BC048337,CR456571,CR607299 | SEC14L2 | GAAATAACACCTTCTCCTA |
| NM_003150,NM_139276,NM_213662,AB451232,AJ012463,AK291933,AK297994,AK301200,AM393108,BC000627,BC014482,EU446657,EU831439,EU831531,L29277 | STAT3 | GGCGTCCAGTTCACTACTA |
| NM_031212,AF267854,AF327402,AF327403,AJ303077,AJ303078,AK056782,AL831943,BC047312,BC058937,BC064541,BC076399,BC094821,CR590395,CR591608,CR601812,CR601821,CR605062 | SLC25A28 | CACCTCAAGTGGAGTTAGA |
| NM_004688,AB451305,AB451436,AK291548,AK298754,BC001268,BC021987,CR604696,U32849 | NMI | GCCAAGCCAGTTCCATTAA |
| NM_020746,AB033097,AB097003,AK023799,AK123956,BX649084 | VISA | GCAGGTCAGTTAACAATTT |
| NM_004696,AK129985,AK223307,AK290720,AK298539,AL833619,BC021664,BX647948 | SLC16A4 | GCAATCAAGTGAGAGCTAA |
| NM_004060,NM_199246,AK297190,AK312913,BC000196,BC007093,BT007134,BX538035,CR619041,D78341,DQ893765,EU176176,L49504,U47413,U53328,X77794,BM479568,BX644635 | CCNG1 | CAACTGACTTGATCCGAAT |
| NM_031458,AF307339,AK292959,AK313494,BC017463,BC039580,CR596753 | PARP9 | CTTTAAAGCTGCTTCAGAA |
| NM_016323,AB027289,AK093155,AK301756,AK302142,AY337518,BC140716 | HERC5 | GGTTTCATTTAGTGGAGAA |
| NM_001007245,NM_001550,AK225222,AK308066,AK313211,BC001272,BT019889,BT019890,CR602983,CR614885,CR616001,Y10313 | IFRD1 | CAGCTCTTGAAGGTATTAA |
| NM_031212,NM_001109515,XM_001056340,AF267854,AF327402,AF327403,AJ303077,AJ303078,AK056782,AL831943,BC047312,BC058937,BC064541,BC076399,BC094821,CR590395,CR591608,CR601812,CR601821,CR605062,BC166775 | SLC25A28, LOC679437, Slc25a28 | GCCAGAGTAATTTACCAGA |
| NM_000161,NM_001024024,NM_001024070,NM_001024071,BC025415,U19523,U66095,U66097,Z29433 | GCH1 | GCTGTTGTCTTATTAGTAA |
| NM_017554,AK098816,AY134858,BX648758,DQ063584,DQ063585 | PARP14 | CTCAGTGCCTTAAATTATA |
| NM_000201,AK130659,AK298054,AK298983,AK301412,BC015969,CR617464,J03132,M24283,X06990 | ICAM1 | CCACGCATCTGATCTGTAG |
| NM_022168,AF095844,AK056293,AK292941,AK314611,AY017378,BC078180,BC111750 | IFIH1 | CCTACAAATTAATGACACA |
| NM_006828,AJ223948,AL834463,AM393382,AY013288,BC014358,BC039857,BC150215 | ASCC3 | GAAGGTATATCTACACATT |
| NM_000270,AK098544,AK126154,CR592961,CR594642,CR595160,CR598701,CR598851,CR598982,CR599285,CR600826,CR601251,CR601413,CR604278,CR604317,CR606974,CR608191,CR608198,CR608316,CR612303,CR612939,CR614792,CR624654,CR625788,X00737 | NP | CTACTAGCTCTTTGAGATA |
| NM_152703,AB095926,AF474973,AK097204,AK304307,AK304779,AL832264,BC029108,BC038974,BC127117,BC127118 | SAMD9L | GAGACAAATTTCAACATGA |
| NM_006417,BC022870,CR621445 | IFI44 | GAATCTAAGGGAGGAAATT |
| NM_024911,AK026744,AK074583,AK074984,BC137113,CR591378,CR602492,DQ323735 | GPR177 | CAGGTAGCCCACTGAGTTA |
| NM_001356,NM_001108246,AB208983,AB451220,AB451343,AF000982,AF000983,AF061337,AK291153,AK297159,AK304689,AK310891,BC007668,BC011819,U50553,BC085914,CB107740 | DDX3X, Ddx3x | CCATCTTGAGTCAGATTTA |
| NM_001356,AB208983,AB451220,AB451343,AF000982,AF000983,AF061337,AK291153,AK297159,AK304689,AY776161,BC007668,BC011819,U50553 | DDX3X | CTGGCAACCTCATTCTTTA |
| NM_003810,AK296085,BC020220,BC032722,CR594331,CR598109,U37518,U57059 | TNFSF10 | GCAATAACCTCAAAGTGAC |
| NM_000030,AB292648,AF191687,BC132819,CR614071,CR620771,X53414,X56092 | AGXT | CTCCCGGGAATGTTTAATA |
| NM_032505,AB058745,AK096640,BC117487 | KBTBD8 | CCTACACTTCCAGAGTTAT |
| NM_001136540,NM_001136541,NM_003661,NM_145343,AF019225,AF305224,AF305428,AF323540,AK298053,AK300454,AK303919,AK309143,AK313752,BC017331,BC112943,BC113867,BC114475,BC127186,BC141823,BC142720,BC143038,BC143039,CR626091 | APOL1 | CAGAGCCAATCTTCAGTCA |
| NM_005419,AK094039,AK296939,BC051284,BX640607,CR621797,M97934 | STAT2 | CCCACACTATGCATGGTAT |
| NM_001562,AF380360,AY044641,AY266351,BC007007,BC007461,BC015863,CR541973,CR542001,D49950,U90434 | IL18 | GAAATCGGCCTCTATTTGA |

| | | |
|---|---|---|
| NM_001048205,NM_005132,AK001435,AK291191,AL832973,BC004159,BC010887,BC019326,CR609880 | REC8 | GAGCTATTGTTCAAGCAGA |
| NM_006573,NM_033622,AF116456,AF132600,AF134715,AF136293,AF186114,AK309629,AY129225,AY129226,AY129227,AY129228,AY302751,AY358881,BC020674,CR541818,DQ891413,EU176634,AF352245,AK034121,AK079180,AK155288,AY290823,BC106840,BC106841 | TNFSF13B, Tnfsf13b | CATGGCTTCTCAGCTTTAA |
| NM_000379,BC166696,CR614711,D11456,U06117,U39487 | XDH | CTCTTCCTGGCTGCTTCTA |
| NM_020414,AB209382,AF134475,AF145022,AF214731,AK025162,AK296726,AK297232,AK304504,AL136886,AM393538,BC008847,BC009406,BC096826,BX537533,CR596324,CR615921,CR619609,CR624535,DQ893523,DQ896510 | DDX24 | CCGGCTGTGGGAATTAATT |
| NM_032855,AK027792,AK131222,AK307563,BC016826,BC025237 | HSH2D | CCACACTCCTGAATGCCTT |
| NM_021035,AB037825,AK023836,BC156357 | ZNFX1 | GGAATGTACTGCATCGGAA |
| NM_000783,NM_057157,AK027560,AK075374 | CYP26A1 | CAGCTTATCTAACATGTCA |
| NM_002038,NM_022872,NM_022873,AK024814,AK314488,BC011601,BC015603,BN000257,BT006850,X02492 | IFI6 | CCACCCACAAGTATCTCGA |
| NM_021035,AB037825,AK000573,AK023836,BC062347,BC156357 | ZNFX1 | CAACCAGCTTGCTTCTGAA |
| NM_001134486,NM_052942,AF288815,AF328727,AF430642,AF430643,AK090479,AK315064,AL832285,AY358953,BC031639,BC033761,DQ895650,EU176216 | GBP5 | CTGGAAGCATCCTCGGATT |
| BC037552 | PPM1K | GCAGCTATGGGTTTCTTCT |
| NM_001511,BC011976,J03561,X12510 | CXCL1 | GTCCTATTATATTCATTCT |
| NM_000304,NM_153321,NM_153322,BC019040,BC091499,CR592810,CR593633,CR597127,CR597311,CR598725,CR602788,CR602976,CR603134,CR604376,CR604692,CR604834,CR605458,CR608530,CR610349,CR611493,CR611595,CR612069,CR615033,CR615858,CR618437,CR618628,CR618649,CR622251,CR624540,D11428,L03203,M94048 | PMP22 | CTGAATAATTCTGTGTAAT |
| NM_001040020,NM_014888,BC024200,BC068526,D87120 | FAM3C | CTGTGTTTATCTAACTTCA |
| NM_138287,AK125086,AL833598,AY225123,BC040372,BC042191,BC060509,BX648267,BX648645 | DTX3L | CTGATTTAATGCCAGTCTA |
| NM_021100,AF097025,AJ010952,AK001265,AK001470,AK056242,AK091049,BC065560,CR594451,CR610756 | NFS1 | GACTCCACCAGTTATTCTA |
| NM_007115,AF086484,AJ419936,AJ421518,BC030205,M31165 | TNFAIP6 | AGTACTACTTCTACTGGAA |
| NM_000062,NM_001032295,AB209826,AK293054,AK303809,AK303840,AK312626,AY732485,BC011171,BT006966,CR590689,CR590941,CR591439,CR594035,CR600380,CR603289,CR604184,CR606235,CR607756,CR608695,CR610016,CR612050,CR612675,CR612737,CR613065,CR614264,CR614379,CR614595,CR615596,CR619147,CR619152,CR620172,CR620983,CR621088,CR621653,CR626630,CR626631,DQ891799,DQ894982,M13656,M13690,M14036,X07577 | SERPING1 | ATGGAACCCTTTCACTTCA |
| AF105151,AK225657,AK299603,M74447 | TAP2 | CGCCTTGTACCTGCTCATA |
| NM_002717,AK095153,AK303981,BC041071,M64929 | PPP2R2A | CTCCATGTCTGCTAGCCAT |
| NM_004458,NM_022977,NM_001033600,NM_019477,NM_207625,NM_053623,AB061713,AB061714,AF030555,AK292070,AK294915,AK307566,BC034959,DQ890835,DQ893990,Y12777,AB033885,AB033886,AB033887,AJ243502,AK054387,BC016416,BC058663,DR004263 | ACSL4, Acsl4, Acsl4 | GAGCAGATACTCTGGATAA |
| NM_004458,NM_022977,NM_001033600,NM_019477,NM_207625,NM_053623,AB061713,AB061714,AF030555,AK292070,AK294915,AK307566,BC034959,DQ890835,DQ893990,Y12777,AB033885,AB033886,AB033887,AJ243502,AK054387,BC016416,BC058663,DR004263 | ACSL4, Acsl4, Acsl4 | GAGCAGATACTCTGGATAA |
| NM_006828,AK299117,AL834463,AM393382,BC026066,BC150215 | ASCC3 | GACTTGAACTTATTGAGAA |
| NM_022168,NM_001109199,AF095844,AK056293,AK292941,AK314611,AY017378,BC111750,AF374384,AK018602,AK153018,AK153344,AY075132,BC004031,BC025508,BC080200,BC168680 | IFIH1, Ifih1, Ifih1 | GAGAGAAGATGATGTATAA |
| NM_001130020,NM_001130021,NM_005177,AK223554,BC032398,BX648978,CR627443,L78933,Z71460 | ATP6V0A1 | CTGAGTCTGTTCAACCATA |
| NM_006820,AB000115,AK223087,AK225442,AK293385,AK312272,AL832618,BC015932 | IFI44L | GACATAAAGAGGATAATTA |
| NM_002535,NM_016817,AK292796,AK292906,BC049215,M87284,M87434 | OAS2 | GGCTCATTGATCTGTATAA |
| NM_001134486,NM_052942,AF288815,AF430642,AF430643,AK315064,AL832285,AY358953,BC031639,BC033761,DQ895650,EU176216 | GBP5 | CACAACATTCCAAGCTCAA |
| NM_001039916,NM_001039917,NM_001039918,NM_001039919,NM_001039920,NM_001135734,NM_133476,BC053361,U80738 | ZNF384 | CTCAGTCCCTGAGAGCCAT |
| NM_000544,NM_018833,AB073779,AB208953,AF078671,AF105151,AF176984,AK222823,AK223300,AK225657,AK225693,AK292963,AK299603,BC002751,BC152839,BT009906,EU832685,FM246456,M74447,M84748,U07844,Z22935,Z22936 | TAP2 | CATGAAGTCTGTCGCTATA |
| NM_031458,AF307339,AK292959,AK313494,BC017463,BC039580,CR596753 | PARP9 | CATTGTAAGTATTCTGAAT |
| NM_001511,BC011976,J03561,X12510 | CXCL1 | CACACTGTCCTATTATATT |
| BC001683,BC001847,BC019625,CR591065,CR592413,CR597001,CR598230,CR598799,CR601069,CR604510,CR608003,CR617904,CR620504,CR624884 | CHAC1 | CACTGAAGGCATTGGCCTA |
| NM_004060,NM_199246,AK297190,AK312913,BC000196,BC007093,BT007134,BX538035,CR619041,D78341,DQ893765,EU176176,L49504,U47413,U53328,X77794 | CCNG1 | GCAAGAGCTTGTATCCAAA |
| NM_002801,AK225658,AK298012,AK312208,BC017198,BC052369,BT019723,BT019724,CR456982,CR597889,CR607369,CR619398,Y13640,BM845639 | PSMB10 | GACAAGAGCTGCGAGAAGA |
| NM_003265,AB445631,AK314208,BC043204,BC059372,BC068487,BC094737,BC096333,BC096334,BC096335,DQ445682,U88879 | TLR3 | CATATATAATTCATGCCTA |
| NM_006401,BC019658,CR595726,U70439,Y07570,Y07969,BP395780 | ANP32B | CAGTTACACTGAGATTGTA |
| NM_001729,BC011618,S55606 | BTC | GACAGAATGTGTCTCAGGA |
| NM_016376,AB033081,AK025483,AK057047,AK074324,AK292930,BC060812,BC148355,BC152991,CR933717 | ANKFY1 | GAGCCAAAGTGAACGAATT |
| NM_014705,AB018259,AY233380,BC117688,BC117689 | DOCK4 | CATTCTTTGTCAAAGAATA |
| NM_001729,AM392725,AM392995,AM393303,AM393550,BC011618,S55606 | BTC | CACCAGAAGTCCTGAAACT |
| NM_024119,AK021416,AK097669,AK225549,BC014949,BM723672 | DHX58 | CTACCAAGGCCACTTCTAT |
| NM_002717,AK095153,AK303981,BC041071,M64929 | PPP2R2A | GTCATGCTATGGGATTTAA |
| NM_000864,BC007720 | HTR1D | GAGCTTAAAGGAGGGTGAA |
| NM_006121,AK313986,BC063697,M10938 | KRT1 | GCAGGAAATTGATTTCCTT |
| NM_000600,BC015511,BT019748,BT019749,CR450296,CR590965,CR626263,DQ891463,DQ894639,M14584,M18403,M29150,M54894,S56892,X04403,X04430,X04602 | IL6 | GACATGACAACTCATCTCA |
| NM_021105,AB006746,AF098642,BC021100,BC032718,BC070251,CR607459 | PLSCR1 | GTTAATATTTCTACATGAA |
| NM_033109,AJ458465,AK307589,AY027528,BC009057,BC053660,CR625599,DA998523 | PNPT1 | GGCAACAGGAAATTAGAAA |
| NM_003895,NM_203446,AB020717,AF009039,AF009040,AK307762,BC019630,BC098395 | SYNJ1 | GTGAACATATGCTAAGTAA |
| NM_000463,NM_001072,NM_007120,NM_019075,NM_019076,NM_019077,NM_019078,NM_019093,NM_021027,NM_205862,AF030310,AF056188,AF462267,AF462268,AK025403,AK290834,AK313488,AK313510,AK313623,AY435136,AY435137,AY435139,AY435141,AY435143,AY435144,BC011409,BC019861,BC020971,BC043491,BC053576,BC058844,BC069210,BC121036,BC128414,BC128415,BC131623,BC139784,BC141470,BC146418,BC153172,BC156848,BC166641,J04093,M57899,M57951,S55985,U89507,U89508,BM924331 | UGT1A1, UGT1A10, UGT1A3, UGT1A4, UGT1A5, UGT1A6, UGT1A7, UGT1A8, UGT1A9 | GTTCCCATGGTGTTTATGA |

| | | |
|---|---|---|
| NM_000584,AK131067,BC013615,CR594973,CR600500,CR601533,CR601902,CR603686,CR619554,CR623683,CR623827,M17017,M26383,Y00787 | IL8 | CAGTGAAACTTCAAGCAAA |
| NM_006745,AK292418,AK309206,BC010653,BC107879,CR623543,DQ891060,DQ894237,U60205,U93162,BX441001 | SC4MOL | GGAACTATATGTTGAATAA |
| NM_006985,NM_178541,XM_001723375,XM_001724221,XR_016512,XR_016550,AF132984,AF229069,AK124516,AK160377,BC008178,BC010188,BC039707,BC146422 | LOC100132540, LOC100132620, LOC339047, LOC642778, LOC642799, NPIP | CGGATGATAATCTCAAGAA |
| NM_006828,AK299117,AL834463,AM393382,AY013288,BC026066,BC150215 | ASCC3 | CAGTTTATATCCAAGACTT |
| NM_001003819,NM_001003827,NM_021616,NM_130389,AB039902,AB039903,AF220143,AF220144,AK027876,AL583914 | TRIM34, TRIM6-TRIM34 | GATTTAGTGTCTGGAACAT |
| NM_000584,AK131067,BC013615,CR594973,CR595357,CR600500,CR601533,CR601902,CR603686,CR619554,CR623683,CR623827,M17017,M26383,Y00787 | IL8 | CACAGTCAATATTAGTAAT |
| NR_000035,AF464191,AF464192,AF464193,AF464194,AF464195,AF486844,AF486845,AF486846,AF486847,AY358222 | CASP12 | CTTGGATACTCAGTGGTTA |
| NM_003141,AK225564,BC010861,M34551,M62800,BU594670 | TRIM21 | CAGTGAAGCAGCCTCCTTA |
| NM_022750,NM_172893,AL136766,AL137255,BC044660,AK036886,AK053205,AK156623,AK162200,BC120733,BC137645 | PARP12, Parp12 | CGGGAAGAACTGTAGGAAT |
| NM_020414,AB209382,AF134475,AF145022,AF214731,AK025162,AK296726,AK297232,AK304504,AL136886,AM393538,BC008847,BC009406,BC096826,BX537533,CR596324,CR615921,CR619609,CR624535,DQ893523,DQ896510 | DDX24 | CGGCTGTGGGAATTAATTA |
| NM_001135651,NM_001135652,NM_002759,AK290655,AK313818,AY302136,BC007769,BC022314,BC040851,BC057805,BC093676,BC101475,CR600308,M35663,M85294 | EIF2AK2 | GCCAGAAGGATTTCATTAT |
| NM_000168,AK299299,AK308429,BC113616,BC117168,M57609 | GLI3 | CAGCTTGTGCACCATATAA |
| NM_001560,AB209849,AK313467,BC009960,BC015768,CR603161,DQ893454,DQ896296,U81379,U81380,Y09328,Y10659 | IL13RA1 | CTTCCACAATGATGACCTA |
| NM_015668,AK292559,AK302475,AK309194,AY009106,BC036665,BC047060,BC050053,BC092411 | RGS22 | CGATGAAGATGAGACCATT |
| BC001683,BC001847,BC019625,CR591065,CR592413,CR597001,CR598230,CR598799,CR601069,CR604510,CR608003,CR617904,CR620504,CR624884 | CHAC1 | CTCTTACCCACTTGGTTGT |
| XM_001722453,XM_001723441,XR_037985,XR_038167,XR_040965,XR_040966,AK021862,BC098263,BC098294,BC098363 | FAM106A, LOC100128973, LOC100129396, LOC100132568 | CTCTGAGGAACTTATTAGA |
| NM_006985,NM_178541,NR_003610,XR_016512,XR_016550,XR_041580,AF132984,AK124516,AK131084,AK131085,AK160377,AK302708,BC039707,BC063633,BC146422,CR619663,BE253379 | LOC339047, LOC399491, LOC642778, LOC642799, NPIP, PDXDC2 | CCGTAGACAGGAAGGAATT |
| NM_001647,BC007402,J02611 | APOD | GCTGCACCCACTCCATGTT |
| NM_001040147,NM_003784,AF027866,AK290673,AK290876,AK300828,BC069417,BC069442,BC069547,BC106743,BC106744,D88575 | SERPINB7 | CAGATACAATGGTGGCATA |
| NM_004688,AB451305,AB451436,AK291548,BC001268,BC021987,CR604696,U32849 | NMI | GAGTCAGATTCCAGGTTTA |
| NM_014398,AB013924,AJ005766,AK302894,AK313172,BC032940,DQ892118 | LAMP3 | GGAAGCAGACTCTGTATAA |
| NM_000544,NM_018833,AB073779,AB208953,AF078671,AF105151,AF176984,AK222823,AK223300,AK225657,AK225693,AK292963,AK299603,BC002751,BC152839,BT009906,EU832685,FM246456,M74447,M84748,U07844,Z22935,Z22936 | TAP2 | CAGACCCTGGTATACATAT |
| NM_003190,NM_172208,NM_172209,AB010639,AF009510,AF029750,AF067286,AF314222,BC080574,CR600344,CR614167,CR618722,EU693375,Y13582 | TAPBP | GAGTGAGACTGGGACAAGA |
| NM_152899,NM_172374,AJ880386,AJ880387,AJ880388,AJ880389,AY358933,BC026103,BC064378,BC090852,BC131625,CR592016,CR622916,CR625444,DQ079587,DQ079588,DQ079589 | IL4I1 | CTCTCAACCAGGCCCTCAA |
| NM_000146,XR_017149,XR_019548,XR_037197,NM_008049,NM_010240,XM_001471608,XM_001478411,XM_357312,XM_486478,XM_904376,XM_909091,XM_973894,XR_032412,XR_032597,NM_001014009,NM_022500,XM_001056161,XM_001061623,XM_001070733,XM_001077872,XM_001078366,XM_574537,XM_576192,XM_577041,XR_005676,XR_006207,XR_006269,XR_006880,XR_007199,XR_007698,XR_007930,XR_008262,XR_008570,XR_009006,XR_009181,AK026534,AK130191,AK130205,AK131048,AK131050,AK131053,AK307065,AK311773,AY207005,AY466472,BC002991,BC004245,BC008439,BC008441,BC013928,BC016346,BC016354,BC016715,BC018990,BC021670,BC050625,BC058820,BC062708,BC067772,BC105971,BX571748,CR456715,CR596451,CR610876,DQ892980,DQ896228,EU831925,EU832020,M10119,M11147,Y09188,AK002242,AK002253,AK002547,AK011009,AK011029,AK011244,AK075620,AK088647,AK145265,AK145658,AK150480,AK150481,AK150960,AK151255,AK152030,AK152385,AK152393,AK152564,AK152612,AK159574,AK167410,AK168310,AK168735,AK168862,AK168866,AK169100,AK169159,AK213945,BC019840,BC020403,BC081462,BC083350,BC085309,BC092259,BC106145,BC106146,BC150761,J04716,S89400,BC086583,BC088756,K01930,L01122,U75408,BY134899 | FTL, GBP4, LOC392437, SEC62, 382986, EG383891, EG665937, ENSMUSG00000062382, Ftl1, Ftl2, LOC434624, LOC545679, LOC630762, Ftl, LOC364392, LOC364842, LOC367190, LOC680217, LOC682465, LOC686673, LOC688591, LOC690464, RGD1306939, RGD1560687, RGD1561055, RGD1562192, RGD1566189 | CAGATTCGTCAGAATTATT |
| NM_017633,NM_001106844,AF350451,AJ420592,AK000044,AK056057,AK292109,BC000683,BC007351,BX648876 | FAM46A, Fam46a | CTGCAATTAAGAATCATTT |
| NM_002818,NM_017257,XR_005793,XR_007517,AK026580,AK225876,AK311569,AY771595,BC004368,BC019885,BC072025,BX161498,CR541657,CR541743,CR594185,CR600073,CR601043,CR615548,CR618033,CR620148,D45248,DQ893404,DQ896722,BC058486,D45250 | PSME2, LOC304754, Psme2 | CAAGGATGATGAGATGGAA |
| NM_004458,NM_022977,NM_001033600,NM_019477,NM_207625,NM_053623,AB061713,AB061714,AF030555,AK292070,AK294915,AK307566,BC034959,DQ890835,DQ893990,Y12777,AB033885,AB033886,AB033887,AJ243502,AK054387,BC016416,BC058663,DR004263 | ACSL4, Acsl4, Acsl4 | AGCAGATACTCTGGATAAA |
| NM_004458,NM_022977,NM_001033600,NM_019477,NM_207625,NM_053623,AB061713,AB061714,AF030555,AK292070,AK294915,AK307566,BC034959,DQ890835,DQ893990,Y12777,AB033885,AB033886,AB033887,AJ243502,AK054387,BC016416,BC058663,DR004263 | ACSL4, Acsl4, Acsl4 | AGCAGATACTCTGGATAAA |
| NM_005567,XM_001726123,AK055977,AK057776,AK293183,AK295442,AK301040,AK301444,AK307665,BC002403,BC002998,BC015761,CR592307,CR594407,CR595010,CR603304,CR608142,CR610951,CR619022,DQ892224,DQ895424,L13210,X79089 | LGALS3BP, LOC100133842 | GTCAAATATTCTTCTGATT |
| NM_015474,AB013847,AB208944,AF228421,AK027811,AK304187,AK304795,AK315169,AL050267,BC036450,CR614403 | SAMHD1 | CTTCCTTTATGAGATAGTA |
| NM_001562,AF380360,AY044641,AY266351,BC007007,BC007461,BC015863,CR541973,CR542001,D49950,U90434 | IL18 | GACCATATTTATTATAAGT |
| NM_016118,AF155099,AF300717,AF459743,AY129295,BC046354,CR593223,CR611732 | NUB1 | CCGTCAATGAGATACTGGA |
| NM_001007245,NM_001550,AK308066,AK313211,BC001272,BT019889,BT019890,CR602983,CR614885,CR616001,Y10313 | IFRD1 | GTTAGCATCTGTTCTTTGT |
| NM_002675,NM_033238,NM_033239,NM_033240,NM_033244,NM_033246,NM_033247,NM_033249,NM_033250,AB208950,AB209051,AB209411,AF230401,AF230402,AF230403,AF230404,AF230405,AF230406,AF230407,AF230408,AF230409,AF230410,AF230411,BC000080,BC020994,BT009911,M73778,M79462,M79463,M79464,M80185,S50913,X63131 | PML | GCACACCAGTGGTTCCTCA |
| NM_003265,AB445631,AK314208,BC017954,BC059372,BC068487,BC094737,BC096333,BC096334,BC096335,DQ445682,U88879,DA374150 | TLR3 | CTATCTCAACTTTCTGATA |
| NM_014547,AF177171,AF237631,AK310344,AK312569,BX647277,CR611644 | TMOD3 | CTCATCTTGTTGAAGTTAA |
| NM_006435,BC009696,CR604902,X02490,X57351,BG164993,BG257141 | IFITM2 | CCCACGTACTCTATCTTCC |
| AK090822,AK094227,BX648468,CR936784 | ATL3 | CCTTAAAGGAGTTAATACT |
| NM_017912,AF336798,AK000644,AK097168,AK225506,AK295832,AL833664,AY653201,AY653202,AY653203,BC035775,BC042047,BX647121 | HERC6 | GAATGTTCATGTATCCTGA |

| | | |
|---|---|---|
| NM_001040078,NM_002308,NM_009587,NR_024043,AB005894,AB006782,AK097892,AK126017,AK223232,AK290263,AK301624,AK315818,BC034392,BC073889,BC110340,CR597107,CR598297,CR602208,CR604285,CR608609,CR614567,CR616420,CR622120,CR626506,Z49107 | LGALS9, LGALS9B, LGALS9C | CCTGGGATCTGGGCTTTAA |
| NM_006573,AF116456,AF132600,AF134715,AF136293,AF186114,AK309629,AY129225,AY129226,AY129227,AY129228,AY302751,AY358881,BC020674,CR541818,DQ891413,EU176634 | TNFSF13B | GTGACTTTGTTTCGATGTA |
| NM_014314,AF038963,AK023661,AK125989,AK301678,AK315040,BC070029,BC132786,BC136610,BX647917 | DDX58 | GAATTTGGAACACAGAAAT |
| NM_014314,AF038963,AK023661,AK125989,AK301678,AK315040,BC070029,BC132786,BC136610,BX647917 | DDX58 | GAATTTGGAACACAGAAAT |
| NM_001032394,NM_001032395,NM_020455,NM_198569,AB183546,AB183547,AB183548,AB183549,AK075087,AK092519,AL080079,BC075798,BX648315 | GPR126 | GGAGGTTACATATGGATGA |
| NM_024119,AK021416,AK097669,AK225549,AK300900,BC014949 | DHX58 | GTTATGGGCTCTTGACCAA |
| NM_021035,AB037825,AK023836,BC030789,BC041630,BC156357,AU140771 | ZNFX1 | CGAGGATTGTCATAGTGGA |
| NM_000161,NM_001024024,NM_001024070,NM_001024071,BC025415,U19523,U66095,U66097,Z29433 | GCH1 | CACAGGCTGTTGTCTTATT |
| NM_020830,AB037856,AK022888,AK023415,BC040525,BC065934 | WDFY1 | CAGTATCTTTGCTAATCTT |
| NM_002800,NM_148954,AK303118,BC065513,BX641100,CR541656,CR621193,S75169,U01025,X62741,BI832878 | PSMB9 | CTGCAAATGTGGTGAGAAA |
| NM_000379,BC166696,D11456,U06117,U39487 | XDH | CAGGATCTCTCTCAGAGTA |
| NM_000201,AK130659,AK298054,AK301412,BC015969,CR617464,J03132,M24283,M55091 | ICAM1 | GACATGATTGATGGATGTT |
| NM_031458,AF307339,AK292959,AK313494,BC017463,BC039580,CR596753 | PARP9 | GATTTAACTTGTTCTGTAA |
| NM_021629,AK001890,AK022599 | GNB4 | GAGTTAGAGAGCTATATTA |
| NM_002984,AY766446,AY766448,BC027961,BC107433,J04130,M23502,M25316,M57503,X16166,X53683 | CCL4 | GTCATTTCCATTATTTATA |
| NM_033064,AB058775,AK092309,AK125457,AK299522,AY220297,BC008736,BC026217 | ATCAY | CTGAAGAAGTGCTACCAGA |
| NM_001001438,NM_002340,AK226141,AK296313,AK312489,BC035638,D63807,DQ891234,DQ894418,S81221,U22526,X87809 | LSS | CCGTGCAGAAGCTGTATGA |
| NM_007346,AF109134,AF172449,AF172450,AF172451,AF172452,AF172453,AK022234,AK024485,AK314696,BC008768,BC014137,BC032666,DQ892943,EU176723,BM551600 | OGFR | CTGTTTCATTGAGGACATT |
| NM_020119,NM_024625,AK055851,AK292811,BC025308,BC033105,BC040956,BX571742,BX647974 | ZC3HAV1 | CAAATATTCTCATGAGGTT |
| NM_019096,NM_019581,AB024574,AF168990,AJ420518,AK000430,AK290267,AL834331,BC020980,BC028347,BC064968,CR607185,AB024573,AF168991,AK087511,AK146415,AK148020,BC049089 | GTPBP2, Gtpbp2 | GAGTTCCAGGTGGATGAAA |
| NM_004116,NM_054033,BC002614,BC050998,CR614690,CR616275,CR626535,D38037,L37086,S69800,S69815 | FKBP1B | CTCAGACATGAAATGTACA |
| NM_001710,AK130533,AK223400,AK304045,BC004143,BC007990,CR590153,CR590439,DQ892313,DQ895516,J00185,K01566,L15702,S67310,X72875 | C2, CFB | CCTGATCAAGCTCAAGAAT |
| NM_003141,AK225564,AY742713,BC010861,EU446706,M34551,M62800 | TRIM21 | CCAGCACTTTCACTCTGGA |
| NM_001030287,NM_001040619,NM_001674,NM_004024,NM_007498,AB078026,AB078027,AB078028,AB209032,AK125255,AK312998,BC006322,BT006996,CR450334,CR600751,CR602765,CR612905,CR614684,CR614862,CR620886,CR626784,DQ892723,DQ895972,L19871,AB291912,AK133965,AK156786,AY329367,BC019946,BC064799,CT010291,D50524,U19118,AU135799 | ATF3, Atf3 | CTCTTTATCCAACAGATAA |
| NM_031212,NM_145156,NM_001109515,XM_001056340,AF267854,AF327402,AF327403,AJ303077,AJ303078,AK056782,AL831943,BC047312,BC058937,BC064541,BC076399,BC094821,CR590395,CR591608,CR601812,CR601821,CR605062,AF377993,AF377994,AK034173,AK155044,BC023172,BC166775 | SLC25A28, Slc25a28, LOC679437, Slc25a28 | CCATTCACTTCATGACCTA |
| NM_031458,NM_030253,AB209742,AF307339,AK123412,AK130147,AK292959,AK313494,AL713679,BC017463,BC039580,BX648869,AK037903,AK050032,BC003281,BC010312 | PARP9, Parp9 | CACCACATCATTGAGAATA |
| NM_001142315,NM_001142316,NM_005574,NM_001142335,NM_001142336,NM_001142337,NM_008505,XM_001479884,NM_001037358,AF257211,BC034041,BC035607,BC042426,BC073973,CR604507,CR614368,CR625714,X61118,AK013416,BC057880,AV361906,BX523983 | LMO2, Lmo2, LOC100048263, Lmo2 | CACAGATTGTTTCTATACA |
| NM_015459,NM_146091,AK023383,AK097588,AK301910,AL117600,BC077727,CR936784,AK028842,AK030660,AK077752,AK147720,AK169476,AK170550,BC002149,BC017138 | ATL3, Atl3 | GAGTTTCCCTTATGAATAT |
| NM_002818,NM_001029855,NM_011190,AK026580,AK225876,AY771595,BC004368,BC019885,BC072025,BX161498,CR541657,CR541743,CR594185,CR600073,CR601043,CR615548,CR618033,CR620148,D45248,DQ893404,DQ896722,AK012344,AK088654,AK143556,BC002301,BC005680,BC057859,D87910,U60329,AA216967,BY498670 | PSME2, Psme2, Psme2b-ps | CCTTCTATGCTGAGCTTTA |
| NM_020858,NM_024966,NM_153616,NM_153617,NM_153618,NM_153619,NM_172537,NM_199238,NM_199239,NM_199240,NM_199241,AB040912,AF389427,AF389428,AF389429,AF389430,AK021660,AK290032,BC150253,AK031307,AK052232,AK084922,AK122515,BC060680 | SEMA6D, Sema6d | CTATGAAAGGCAAGCATAA |
| NM_001017369,NM_006745,NM_025436,NM_080886,AK292418,AK309206,BC010653,BC107879,CR617900,CR623263,CR623543,DQ891060,DQ894237,U60205,U93162,AK005090,AK005441,AK045059,AK146684,AK151135,AK154682,AK169017,BC006802,BC063155,BU940384,BX441001 | SC4MOL, Sc4mol, Sc4mol | GAAAGAATGCCAAGATGGT |
| NM_014873,NM_001134829,NM_172266,NM_001047894,NM_001109376,XM_001067105,XM_001067154,AY561706,BC034621,D86960,AK133814,AK172914,BC013667,BC024750 | LPGAT1, Lpgat1, LOC317456, LOC679692, LOC683760 | GCCTCCAGTGGATAATAGA |
| NM_014873,NM_001134829,NM_172266,AY561706,BC034621,D86960,AK133814,AK172914,BC013667 | LPGAT1, Lpgat1 | CTACATCTGCTATGTAATT |
| NM_020140,NM_152788,NM_181670,NM_001128086,XM_917405,AF145204,AF164792,AK289768,AK294191,AY281131,AY281132,AY283057,AY620824,AY753193,BC026313,BC068451,BC091512,BC142669,BC150204,BC160005,CR602042,AK019361,AK032211,AK034154,AK049899,AK132228,AK163750,BC098373 | ANKS1B, Anks1b, Ipw | CACCTGATGGCTAATGGAT |
| NM_000298,NM_181871,AB015983,AK225947,AK298399,BC025737,EU176228,M15465 | PKLR | GCATCAAGATCATCAGCAA |
| NM_001134486,NM_052942,AF288815,AF328727,AF430642,AF430643,AK090479,AK315064,AL832285,AY358953,BC031639,BC033761,DQ895650,EU176216 | GBP5 | GAGACTCTACTAGATGCAA |
| AK056767 | MAFK | CGGCCACTACCTAATTTAT |
| NM_016376,NM_020740,AB033081,AB037360,AK025483,AK057047,AK225170,AK292930,BC052308,BC060812,BC148355,BC152991,CR933717 | ANKFY1 | CGTGGTAAATGGGACTTCA |
| NM_002407,AF071219,AJ224173,BC062218 | SCGB2A1 | GTGTAATATGAAGAGTAAT |
| NM_015668,AK097399,AK292559,AL117544,AY009106,BC036665,BC050053,BC092411 | RGS22 | GAGTTAGGACCATATGTAT |
| NM_014873,D86960 | LPGAT1 | CAATAGATATGTGAGTTAA |
| NM_014398,AB013924,AJ005766,AK302894,AK313172,BC032940,DQ892118 | LAMP3 | GATGAAGAATCATATTATA |
| NM_002818,BX161498,CR541657,CR594185,CR600073,CR601043,CR615548,CR620148 | PSME2 | GAGAAGAAAGAAGTCCATA |
| NM_020971,AB046862,AF082075,AF311855,AF324063,AF324064,AY004226,AY004227 | SPTBN4 | CTCGAACTTGGCTGGCATA |
| NM_000566,AK291451,AK291502,BC032634,BC152383,CR591922,CR608315,CR622515,DQ786309,DQ890842,DQ893997,L03418,X14355,X14356 | FCGR1A | GAATATCTGTCACTGTGAA |
| XR_017251,XR_019519,XR_037483,AF061738,AK022055,AK225511,AK298613,AK298790,AK307723,BC006199,BC065564,CR457128 | LAP3, LOC389386 | CCGAAATTATTGAGAAGAA |
| NM_004972,AF001362,AF005216,AF058925,AK292525,AK302618,AY973034,BC039695 | JAK2 | CTTTGTCTTTCGTGTCATT |
| NM_001007245,NM_001550,AK308066,AK313211,BC001272,BT019889,BT019890,BX648799,CR602983,CR614885,CR616001,Y10313,BI915700 | IFRD1 | GAACCAAAGCTAGAAGCAA |

| | | |
|---|---|---|
| NM_001003818,NM_001003819,NM_058166,AB039903,AF220030,AK027664,AK290172,BC047564,BC136871,CR749260,BG721109 | TRIM6, TRIM6-TRIM34 | GCAGGAAACATCTTAGAAA |
| EF535103,J03143 | IFNGR1 | CTTGGTATAGCATATGTGT |
| NM_004116,NM_054033,BC002614,BC050998,CR614690,CR616275,CR626535,D38037,L37086,S69800,S69815 | FKBP1B | CTGTCACTTTCTCTCTTAT |
| NM_000566,BC032634,CR591922,CR622515,DQ786309,X14356 | FCGR1A | GTATGTAACTCTTAAAGCA |
| NM_002343,AF332168,AK093852,AK290859,AK292813,AK303889,AY137470,AY165046,AY178998,AY360320,AY493417,AY875691,BC015822,BC015823,BC022347,DQ522304,DQ892855,DQ896102,EU779935,M83202,M83205,M93150,U07643,X52941,X53961,CD722125 | LTF | CCCTCCTGTCAGCTGCATA |
| NM_174908,NM_178335,AJ416916,AJ557013,BC065004,CR592179,CR601015,CR603155,CR610519,CR614299,CR618301,CR618316 | CCDC50 | CTGTGAAATTGCTCAGGAA |
| NM_000389,NM_078467,AB209881,AK298901,AK309512,BC000275,BC000312,BC001935,BC013967,CR590203,CR612385,CR617250,CR621127,L25610,L26165,U03106,U09579 | CDKN1A | CATTAGAATTATTTAAACA |
| NM_005465,NM_181690,NM_011785,NM_031575,AF085234,AF124141,AF135794,AJ245709,AK308052,AL117525,AM392791,AM392849,AY005799,BC020479,BC121154,AF124142,AK052953,BC066861 | AKT3, Akt3, Akt3 | CAGCTCAGACTATTACAAT |
| NM_004972,AF001362,AF005216,AF058925,AK292525,AK302618,AY973034,BC039695 | JAK2 | GGCAGAATTAGCAAACCTT |
| NM_001008540,NM_003467,NM_009911,NM_022205,AF025375,AF147204,AF348491,AK129916,AK296674,AK312244,AY242129,BC020968,BT006660,CR594428,CR594588,CR596547,CR598713,CR601301,CR605131,CR610268,CR614199,CR614594,CR614663,CR619476,CR623838,D10924,EU831811,EU831888,L01639,L06797,M99293,X71635,AB000803,BC031665,BC098322,U59760,X99582,Z80111,Z80112,AF452185,BC089804,U54791,U90610,BB872868,CO557829 | CXCR4, Cxcr4, Cxcr4 | CTGGTCATGGGTTACCAGA |
| NM_003745,NM_009896,NM_145879,AB000676,AB005043,AK127621,BC029307,BC148542,BC153088,U88326,AB000677,AB000710,AF120490,AF180302,AK028632,AK154706,BC132366,BC132368,U88325 | SOCS1, Socs1, Socs1 | GTGGCGGCGCCGCGCCGCA |
| NM_003745,NM_009896,NM_145879,AB000676,AB005043,AK127621,BC029307,BC148542,BC153088,U88326,AB000677,AB000710,AF120490,AF180302,AK028632,AK154706,BC132366,BC132368,U88325 | SOCS1, Socs1, Socs1 | GTGGCGGCGCCGCGCCGCA |
| NM_000304,NM_153321,NM_153322,BC019040,BC091499,CR592810,CR593633,CR597127,CR597311,CR598725,CR602788,CR602976,CR603134,CR604376,CR604692,CR604834,CR605458,CR608530,CR610349,CR611493,CR611595,CR612069,CR615033,CR615858,CR618437,CR618628,CR618649,CR622251,CR624540,D11428,L03203,M94048 | PMP22 | CATTCCTTATTATCACTAA |
| NM_001165,NM_182962,BC037420,L49432,U37546,U45878 | BIRC3 | CAACACGTTTGAACTGAAA |
| NM_004048,AB021288,AF072097,AK026463,AY007153,BC032589,BC064910,CR590254,CR591572,CR592576,CR592679,CR593405,CR593761,CR593874,CR594444,CR596238,CR596347,CR596444,CR596717,CR597808,CR599099,CR599510,CR603043,CR603887,CR604110,CR604555,CR604705,CR604949,CR605353,CR606667,CR607236,CR608139,CR608197,CR610094,CR610959,CR612132,CR612519,CR613616,CR613782,CR613955,CR614402,CR614556,CR615574,CR615578,CR615729,CR616171,CR616437,CR619071,CR619828,CR620994,CR621079,CR623073,CR624595,CR625152,CR626821,V00567 | B2M | CTTATACACTTACACTTTA |
| NM_052966,AB050477,AF288391,AK074139,AL137572,BC030531 | FAM129A | AGTGCTTGTTCTAGCATAA |
| NM_052941,AF288814,AK131094,AK312417,AL832576,BC008421,BC017889,BC050625,BC070055 | GBP4 | CTACAAATGACAAGCAATA |
| NM_000161,NM_001024024,NM_001024070,NM_001024071,BC025415,S44053,U19523,U66095,U66097,Z29433 | GCH1 | CCTTATTAATGATTATCTT |
| NM_005531,AB208989,AF208043,AK094968,AK296228,AK307957,AY138863,BC017059,CR603557,CR620182,CR621058,DQ891062,DQ894242,M63838 | IFI16 | AGATCATTGCCATAGCAAA |
| NM_014426,NM_152227,XR_015542,XR_039554,AF121855,AK001793,AK026227,AK054634,AK129505,AK223308,AK308017,AL832967,BC000100,BC062638,BC093623,BC093980,CR590760,CR591673,CR600241,CR601419,CR610846 | LOC728467, SNX5 | AGCATTTAGAAAGAATCTA |
| NM_006398,BC012472,Y12653 | UBD | GTATTACTGAGACTAGTAA |
| NM_001356,AB208983,AB451220,AB451343,AF000982,AF000983,AF061337,AK291153,AK296906,AK297159,AK304661,BC007668,BC011819,U50553,BM833870 | DDX3X | CTATATTCCTCCTCATTTA |
| NM_000201,AK130659,AK298054,AK298983,AK301412,BC015969,CR617464,J03132,M24283,M55091,X06990 | ICAM1 | CATCTGATCTGTAGTCACA |
| NM_152703,AB095926,AF474973,AK097204,AK304307,AK304779,AL832264,BC029108,BC038974,BC127117,BC127118 | SAMD9L | CTGGGAACCTGAAAGCTTA |
| NM_000304,NM_153321,NM_153322,NM_008885,NM_017037,BC019040,BC091499,CR592810,CR593633,CR597127,CR597311,CR598725,CR602788,CR602976,CR603134,CR604376,CR604692,CR604834,CR605458,CR608530,CR610349,CR611493,CR611595,CR612069,CR615033,CR615858,CR618437,CR618628,CR618649,CR622251,CR624540,D11428,L03203,M94048,AK076005,AK171745,BC010765,M32240,M69139,S55427 | PMP22, Pmp22, Pmp22 | CACCAACTGTAGATGTATA |
| NM_003141,AK225564,AY742713,BC010861,EU446706,M34551,M62800 | TRIM21 | CCAATCCGTGGCTGATACT |
| NM_016118,AF155099,AF300717,AF459743,AY129295,BC046354,CR611732 | NUB1 | GATCATGCGGCCACTCATA |
| NM_001136540,NM_001136541,NM_003661,NM_145343,AF019225,AF305224,AF305428,AF323540,AK296099,AK300454,AK303919,AK309143,AK313752,BC017331,BC112943,BC113867,BC114475,BC127186,BC141823,BC142720,BC143038,BC143039,CR626091,BG419709,DA375861,DC340792,DC367266 | APOL1 | CAGATACTGGAGATCCTCA |
| NM_017414,XM_001126794,XM_001128283,XM_001128502,XM_001129362,AF176642,AJ243526,AL136690,BC014896,CR604420 | LOC727996, LOC728216, LOC728253, LOC728438, USP18 | CAACATTAATTCCATATGA |
| NM_001127713,NM_015915,NM_181598,AF131801,AF444143,AK223436,AK290185,AK312518,AL833591,AY032844,BC010708,CR457153,CR599776 | ATL1 | CCATTTCAGAGTCTGATAT |
| NM_001135651,NM_001135652,NM_002759,AF086566,AK290655,AK313818,AY302136,BC007769,BC022314,BC040851,BC057805,BC093676,BC101475,CR600308,M35663,M85294 | EIF2AK2 | CTTGGAACAATGGATTGAA |
| NM_001565,BC010954,X02530 | CXCL10 | CTCTGCATGTTACATAAGA |
| NM_001223,NM_033292,NM_033293,NM_033294,AK223503,AK290114,AK290122,AK296646,AK301037,AK310059,AK313516,AY660536,BC041689,BC062327,CR592117,M87507,U13697,U13698,U13699,X65019 | CASP1 | CTCTCATTATCTGCAATGA |
| NM_004079,AK226095,BC002642,CR592748,CR612707,CR613968,CR623699,CR626534,M90696 | CTSS | GATAAAGTTTGCTAAGTAA |
| NM_004223,NM_198183,AF061736,AK093462,AK129621,AK226165,BC032491,CR601863,CR609116 | UBE2L6 | CTCAGACTGTGAAGTATAT |
| NM_001040020,NM_014888,BC024200,BC068526,D87120 | FAM3C | CCGAAGCTATAGTGGCATA |
| NM_004509,NM_004510,NM_080424,AF280094,AF280095,AK026488,AK301097,AL832300,BC012447,BC019059,CR625933,DA958469,DB474530 | SP110 | CTCTGGAAGCCTGTAGAAA |
| NM_005746,AK292851,BC020691,BC072439,BC106046,CR593609,CR606862,U02020 | NAMPT | GAAATATGAGGAAACAGTA |
| NM_014314,AF038963,AK125989,AK301678,AK315040,BC070029,BC107731,BC132786,BC136610,DN998145 | DDX58 | CAGGTTATTCTGGACTTTA |
| NM_014314,AF038963,AK125989,AK301678,AK315040,BC070029,BC107731,BC132786,BC136610,DN998145 | DDX58 | CAGGTTATTCTGGACTTTA |
| NM_001032394,NM_001032395,NM_020455,NM_198569,AB183546,AB183547,AB183548,AB183549,AK075087,AK092519,AL080079,BC075798,BX648315 | GPR126 | GACTGTGAGTGCAAATAGA |
| NM_080745,NM_182985,AF302088,AF302089,AK226115,AK292252,AL360161,AY305385,BC024199,BC033314,BC047945,DQ232883,EU446573,EU832108,EU832202 | TRIM69 | GTAACACTTTCATGGAGAA |

| | | |
|---|---|---|
| NM_002462,AK093008,AK096355,AK225885,AK315465,BC014222,BC032602,CR592170,M30817,M33882 | MX1 | CACAAATGGAGTACAATAA |
| NM_014873,D86960 | LPGAT1 | CAGTTATACTTGACAATTT |
| NM_001003819,NM_001003827,NM_021616,NM_130389,NM_130390,AB039902,AB039903,AB039904,AF220143,AF220144,AK027876,AL583914,BC136871,BC156770,AK048625 | TRIM34, TRIM6-TRIM34, C130089K02Rik | TGCAAAGATTGGAAGAAGA |
| NM_003150,NM_139276,NM_213659,NM_213660,NM_012747,AB451232,AJ012463,AK291933,AK297994,AK301200,AM393108,BC000627,BC014482,EU446657,EU831439,EU831531,L29277,AK145486,AK153005,AK153472,AK192932,AY299489,AY299490,BC003806,BC037688,L29278,U06922,U08378,X91810 | STAT3, Stat3, Stat3 | ACCAACGACCTGCAGCAAT |
| NM_006398,AK311914,BC012472,DQ894624,Y12653 | UBD | GGGAAGATGATGGCAGATT |
| NM_001032394,NM_001032395,NM_020455,NM_198569,AB183546,AB183547,AB183548,AB183549,AF216967,AK027843,AK075087,AK299785,AY181244,AY426673,BC036008,BC075798,BX640873,BX640971,BX648315 | GPR126 | AGAAAAGTTCTGATAATCT |
| NM_032505,AB058745,AK096640,BC117487 | KBTBD8 | AGGACAACTATGCACTCTT |
| NM_080745,NM_182985,AF302088,AF302089,AK226115,AK292252,BC033314,BC047945,DQ232883 | TRIM69 | AAATTCAGAGTGTTATTAA |
| NM_001025107,NM_001111,NM_015840,NM_015841,NM_001038587,NM_019655,NM_031006,AB209891,AK097241,BC038227,BX538232,BX640741,EU176459,EU176799,U10439,U18121,X79448,X79449,X98559,AF052506,AF291050,AF291876,AK076413,AK089451,AK147614,CT010323 | ADAR, Adar, Adar | ACGGTGTTCACCCTACAAG |
| NM_001974,AK131562,AK290401,AK291518,AK313495,BC059395,X81479 | EMR1 | AGGAGTGGAATGTAGAGAT |
| NM_005567,AK055977,AK057776,AK293183,AK295442,AK301040,AK301444,AK307665,BC002403,BC002998,BC015761,CR595010,CR603304,CR610951,DQ892224,DQ895424,L13210,X79089 | LGALS3BP | TGCTGGTTGCAGGAACCCA |
| NM_001130079,NM_020963,AB046851,AK057353,AK074174,AK226011,AL832805,AL833353,AM393007,AM393678,BC002548,BC004499,BC009312,BC025339,BX647787,DQ892188,DQ896845 | MOV10 | AGGTGGGTTCAGTAGAAGA |
| NM_001128166,NM_001128167,NM_001128168,NM_001128172,NM_001128173,NM_002578,NM_008778,NM_019210,AB102659,AF068864,AF155651,AK128022,AK290504,AK315284,AM943850,AM943851,AM943852,BC117353,BC117355,BC152761,AF082297,AJ496262,AJ496263,AK031853,AK158893,AM943853,AM943854,BC053403,U39738 | PAK3, Pak3, Pak3 | ACCAAGACCAGAGCATACA |
| NM_001128166,NM_001128167,NM_001128168,NM_001128172,NM_001128173,NM_002578,AB102659,AF068864,AF155651,AK128022,AK290504,AK315284,AM943850,AM943851,AM943852,BC117353,BC117355,BC152761 | PAK3 | AGGTGATCCATAGAGATAT |
| NM_001974,AK131562,AK290401,AK291518,AK313495,BC059395 | EMR1 | ACCAATACAGTGGACAGTT |
| NM_001130079,NM_020963,AB046851,AK023297,AK074174,AK226011,AK308874,AK310512,AL833353,AM393007,AM393678,BC002548,BC009312,BC025339,BX647787,DQ892188,DQ896845 | MOV10 | AAGGAGATCGCAGAGATCA |
| NM_001025107,NM_001111,NM_015840,NM_015841,AB209891,AK097241,BC038227,BX538232,BX640741,EU176459,EU176799,U10439,U18121,X79448,X79449,X98559 | ADAR | AGGAACCTGTCATAAAGTT |
| NM_080745,NM_182985,AF302088,AF302089,AK226115,AK292252,AL360161,AY305385,BC024199,BC033314,BC047945,DQ232883,EU446573,EU832108,EU832202 | TRIM69 | GGACAGTTGTCCTTCTACA |
| NM_001025107,NM_001111,NM_015840,NM_015841,AB209891,AK097241,BC017853,BC038227,BX538232,BX640741,EU176459,EU176799,U10439,U18121,X79448,X79449,X98559 | ADAR | AGGGGGATGTCTATAGACA |
| NM_005858,AK308251,AK309529,BC037270,Y11997 | AKAP8 | CGGAAGCAGTTCCAACTTT |
| NM_001003819,NM_001003827,NM_021616,NM_130389,NM_130390,AB039902,AB039903,AB039904,AF220143,AF220144,AK027876,BC136871 | TRIM34, TRIM6-TRIM34 | AGCAGTGCAATGGCTTCAA |
| NM_001130079,NM_020963,AB046851,AK057353,AK074174,AK226011,AK308874,AL833353,AM393007,AM393678,BC002548,BC004499,BC009312,BC025339,BX647787,DQ892188,DQ896845 | MOV10 | AAGAAGAAGCTGCAGGAAT |
| NM_152899,NM_172374,AJ880386,AJ880387,AJ880388,AJ880389,AY358933,BC026103,BC064378,BC090852,BC131625,CR592016,CR622916,CR625444,DQ079587,DQ079588,DQ079589 | IL4I1 | AAGTTCACCCAGTACGACA |
| NM_001008540,NM_003467,AF025375,AF147204,AF348491,AK129916,AK312244,AY242129,BC020968,BT006660,CR594428,CR594588,CR596547,CR598713,CR601301,CR605131,CR610268,CR614199,CR614594,CR614663,CR619476,CR623838,D10924,EU831811,EU831888,L01639,L06797,M99293,X71635 | CXCR4 | AGGAAGCTGTTGGCTGAAA |
| NM_001032394,NM_001032395,NM_020455,NM_198569,AB183546,AB183547,AB183548,AB183549,AK027843,AY181244,BC075798,BX640873,BX640971,BX648315 | GPR126 | AGGATCCTGTTCAAATAAA |
| NM_005567,XM_001726123,AK055977,AK057776,AK293183,AK295442,AK301040,AK301444,AK307665,BC002403,BC002998,BC015761,CR592307,CR594407,CR595010,CR603304,CR608142,CR610951,CR619022,DQ892224,DQ895424,L13210,X79089 | LGALS3BP, LOC100133842 | CCCTCTGACTACAGATACT |
| NM_005082,AB208994,AK310720,BC016924,BC042541,CR617665,D21205 | TRIM25 | AGGTGGAGCAGCTACAACA |
| NM_032505,AB058745,AK096640,BC117487 | KBTBD8 | AGACCAATGTGCTAAGTAT |
| NM_003150,NM_139276,NM_213662,NM_012747,AB451232,AJ012463,AK291933,AK297994,AK301200,AM393108,BC000627,BC014482,EU446657,EU831439,EU831531,L29277,X91810 | STAT3, Stat3 | AGAGTCAAGGAGACATGCA |
| NM_000864,AF498979,BC007720,BT007027,DQ893875,EU176172,M81589 | HTR1D | TCGGCAAGCTTTTCAGAAA |
| NM_001128166,NM_001128167,NM_001128168,NM_001128172,NM_001128173,NM_002578,AB102659,AF068864,AF155651,AK128022,AK290504,AK315284,AM943850,AM943851,AM943852,BC117353,BC117355,BC152761 | PAK3 | GGGATGGATGGCTCTGTTA |
| NM_017414,XM_001126794,XM_001128283,XM_001128502,XM_001129362,AF176642,AJ243526,AL136690,BC014896,CR604420 | LOC727996, LOC728216, LOC728253, LOC728438, USP18 | TGGATCTACGGAGTCTTCT |
| NM_001003819,NM_001003827,NM_021616,NM_130389 | TRIM34, TRIM6-TRIM34 | CACTTTATCACTGATATGA |
| NM_001130079,NM_020963,AB046851,AK057353,AK074174,AK226011,AK308874,AK310512,AL833353,AM393007,AM393678,BC002548,BC004499,BC009312,BC025339,BX647787,DQ892188,DQ896845 | MOV10 | ACGTTAGTGGAGGCAATTA |
| NM_002675,NM_033238,NM_033239,NM_033240,NM_033244,NM_033249,NM_033250,XM_001717388,AB208950,AB209411,AF230401,AF230402,AF230403,AF230404,AF230405,AF230406,AF230410,AF230411,AK300256,BC000080,BC020994,BC034251,BC139795,BT009911,BX647287,M73778,M79462,M79463,M79464,M80185,S50913,X63131 | LOC652671, PML | AGGAAGGTCATCAAGATGG |
| NM_001003818,NM_058166,AF220030,AK023210,AK027664,AK290172,AK293295,AK298301,BC047564,BC065575,CR624250,CR749260,CX760618 | TRIM6, TRIM6-TRIM34 | TGGACCTTCCTGTCTGGAA |
| NM_001356,AB208983,AB451220,AB451343,AF000982,AF000983,AF061337,AK291153,AK296906,AK297159,AK304661,AK304689,AK310891,BC007668,BC011819,U50553,BM833870 | DDX3X | TGGAGTTCTAGCAAAGATA |
| NM_005858,AK308251,BC037270,Y11997 | AKAP8 | CCCAGCTACAGCTACGACT |
| NM_001025107,NM_001111,NM_015840,NM_015841,AB209891,BC038227,BX538232,BX640741,EU176459,EU176799,U10439,U18121,X79448,X79449,X98559,BE886208 | ADAR | AGTGAGTTAATGAAATACA |
| NM_003467,AF025375,AF348491,AK129916,AK296674,AK312244,AY242129,BC020968,BT006660,CR594428,CR594588,CR596547,CR598713,CR601301,CR605131,CR610268,CR614199,CR614594,CR614663,CR619476,CR623838,D10924,EU831811,EU831888,L01639,L06797,M99293,X71635 | CXCR4 | AGTATATACACTTCAGATA |
| NM_001130079,NM_020963,AB046851,AK074174,AK226011,AK308874,AK310512,AL833353,AM393007,AM393678,BC002548,BC004499,BC009312,BC025339,BX647787,DQ892188,DQ896845 | MOV10 | CCGGCAAGACTGTCACGTT |

| | | |
|---|---|---|
| NM_001032394,NM_001032395,NM_020455,NM_198569,AB183546,AB183547,AB183548,AB183549,AF216967,AK027843,AK075087,AK299785,AY181244,AY426673,BC036008,BC075798,BX640873,BX640971,BX648315 | GPR126 | AGGAAAATCTTTGTCTTCA |
| NM_005858,AL050160,BC037270,Y11997 | AKAP8 | CAGAATATGCTGTAATCTA |
| NM_001356,AB208983,AB451220,AB451343,AF000982,AF000983,AF061337,AK291153,AK297159,AK304661,AK304689,AK310891,BC007668,BC011819,U50553 | DDX3X | AGATGAAGATGATTGGTCA |
| NM_032505,AB058745,AK096640,BC117487 | KBTBD8 | AAGAGAAGATCTATGTTTT |
| NM_152899,NM_172374,AJ880386,AJ880387,AJ880388,AJ880389,AY358933,BC026103,BC064378,BC090852,BC131625,CR592016,CR622916,CR625444,DQ079587,DQ079588,DQ079589 | IL4I1 | AGGCGATGAAGAAGTTTGA |
| NM_032505,AB058745,AK096640,BC117487 | KBTBD8 | TCGGAGATCGATCAAAAGA |
| NM_001974,AK131562,AK290401,AK313495,X81479 | EMR1 | ACGATGGACTTTTCCTTGT |
| NM_005082,AB208994,AK310720,BC016924,BC042541,CR617665,D21205,CF121587 | TRIM25 | CAGAGCACCATAGACCTCA |
| NM_005567,XM_001726123,AK055977,AK057776,AK293183,AK295442,AK301040,AK301444,AK307665,BC002403,BC002998,BC015761,CR592307,CR594407,CR595010,CR603304,CR608142,CR610951,CR619022,DQ892224,DQ895424,L13210,X79089 | LGALS3BP, LOC100133842 | CGGAAGTCACAACTGGTCT |
| NM_003141,AY742713,BC010861,EU446706,M34551,M62800 | TRIM21 | GCAGAGTTTGTGCAGCAAA |
| NM_017414,XM_001126794,XM_001128283,XM_001128502,AF176642,AJ243526,AL136690,BC014896,CR604420 | LOC727996, LOC728216, LOC728253, USP18 | AGCATTGTTTTCAAACTAT |
| NM_005858,AK308251,BC037270,Y11997 | AKAP8 | AGCTACGACTATGAGTTCG |
| NM_003150,NM_139276,NM_213662,NM_011486,NM_213659,NM_213660,XM_001474017,NM_012747,AB451232,AJ012463,AK291933,AK297994,AK301200,AM393108,BC000627,BC014482,EU446657,EU831439,EU831531,L29277,AK145486,AK153005,AK153472,AY299489,AY299490,BC003806,BC019168,L29278,U06922,U08378,U30709,X91810 | STAT3, LOC100045296, Stat3, Stat3 | TGAGTTGAATTATCAGCTT |
| NM_152899,NM_172374,AJ880386,AJ880387,AJ880388,AJ880389,AY358933,BC026103,BC064378,BC090852,BC131625,CR592016,CR622916,CR625444,DQ079587,DQ079588,DQ079589 | IL4I1 | TGCAGAAAGGCGATGAAGA |
| NM_003141,AY742713,BC010861,EU446706,M34551,M62800 | TRIM21 | AGGAGCTCATCTCAGAGCT |
| NM_001128166,NM_001128167,NM_001128168,NM_001128172,NM_001128173,NM_002578,AB102659,AF068864,AF155651,AK128022,AK290504,AK315284,AM943850,AM943851,AM943852,BC117353,BC117355,BC152761 | PAK3 | AGATGAACCTTCAACAGCA |
| NM_003141,AK225564,AY742713,BC010861,EU446706,M34551,M62800,BP280435,BP288941 | TRIM21 | TGGTGAACAACCTTAAAGA |
| NM_001974,AK131562,BC059395,X81479 | EMR1 | ACAGAGAACCTCTCAATAA |
| NM_001356,NM_010028,XR_032115,NM_001108246,AF061337,AK291153,AK296906,AK297159,BC007668,BC011819,U50553,AK140990,AK163250,AK182672,DQ858458,L25126,Z38117,BC085914 | DDX3X, Ddx3x, LOC100045923, Ddx3x | AGCTTCAAGAACTCGCAGT |
| NM_001128166,NM_001128167,NM_001128168,NM_001128172,NM_001128173,NM_002578,NM_019210,AB102659,AF068864,AK128022,AK290504,AK315284,AM943850,AM943851,AM943852,BC117353,BC117355,BC152761 | PAK3, Pak3 | TCCAAAGAAACAGTCAACA |
| NM_001008540,NM_003467,AF025375,AF147204,AF348491,AK129916,AK296674,AK312244,AY242129,BC020968,BT006660,CR594428,CR594588,CR596547,CR598681,CR598713,CR601301,CR604071,CR605131,CR610268,CR614199,CR614594,CR614663,CR619476,CR623838,D10924,EU831811,EU831888,L01639,L06797,M99293,X71635 | CXCR4 | TGAGTCTGAGTCTTCAAGT |
| NM_001130079,NM_020963,NM_008619,NM_001107711,AB046851,AK057353,AK074174,AK226011,AL832805,AL833353,AM393007,AM393678,BC002548,BC004499,BC009312,BC025339,BX647787,DQ892188,DQ896845,AK004542,AK150204,AK159295,AK169206,AK173224,BC053743 | MOV10, Mov10, Mov10 | CCTGGAGTTCTGTAAAGAA |
| NM_012346,NM_016553,NM_153718,NM_153719,AK000829,AK125857,AK225817,AL162061,AM393785,BC003663,BC014842,BC050717,BC095410,BC101104,BC101105,BC101106,BC101107,CR541721,DQ892591,DQ896857,X58521,BQ672052,CB961185 | IL4I1, NUP62 | TGGCACTGCAAAGACGGCA |
| NM_003141,AK225564,AY742713,BC010861,EU446706,M34551,M62800 | TRIM21 | AGAGCATACCTGGAAATGA |
| NM_152899,NM_172374,AJ880386,AJ880387,AJ880388,AJ880389,AY358933,BC026103,BC064378,BC090852,BC131625,CR592016,CR622916,CR625444,DQ079587,DQ079588,DQ079589 | IL4I1 | AAGGCGATGAAGAAGTTTG |
| NM_001032394,NM_001032395,NM_020455,NM_198569,AB183546,AB183547,AB183548,AB183549,AF216967,AK027843,AK075087,AK299785,AY181244,BC036008,BC075798,BX640873,BX640971,BX648315 | GPR126 | TGATGGAGATCAAACATCA |
| NM_001003819,NM_001003827,NM_021616,NM_130389,AB039902,AB039903,AF220143,AF220144,AK027876,AL583914,BC136871 | TRIM34, TRIM6-TRIM34 | CATATTTTCAGAGAAAGTT |
| NM_001008540,NM_003467,AF025375,AF147204,AF348491,AK129916,AK296674,AK312244,AY242129,BC020968,BT006660,CR594428,CR594588,CR596547,CR598681,CR598713,CR601301,CR604071,CR605131,CR610268,CR614199,CR614594,CR614663,CR619476,CR623838,D10924,EU831811,EU831888,L01639,L06797,M99293,X71635 | CXCR4 | CACTGAGTCTGAGTCTTCA |
| NM_000864,AF498979,BC007720,BT007027,DQ893875,EU176172,M81589 | HTR1D | AGGTACTGGGCAATCACAG |
| NM_002675,NM_033238,NM_033239,NM_033240,NM_033244,AF230401,AF230402,AF230403,AF230404,AF230405,AF230406,AF230410,AK300256,M73778,M79462,M79463,M79464,M80185,S50913,X63131 | PML | TGGAGAGGATGTCTCCAAT |
| NM_006398,AK311914,BC012472,DQ894624,Y12653 | UBD | TGGCAGATTACGGCATCAG |
| NM_032505,AB058745,AK096640,BC117487 | KBTBD8 | AGGACTCTATCTACTACAT |
| NM_001025107,NM_001111,NM_015840,NM_015841,AB209891,AK097241,BC038227,BX538232,BX640741,EF190449,EF190450,EU176459,EU176799,U10439,U18121,X79448,X79449,X98559,CR982397 | ADAR | AGGAACTGAGTATCTACCA |
| NM_003141,AY742713,BC010861,EU446706,M34551,M62800 | TRIM21 | AGTGGAAACACAGAAATCT |
| NM_001130079,NM_020963,NM_008619,NM_001107711,AB046851,AK023297,AK074174,AK226011,AK308874,AK310512,AL833353,AM393007,AM393678,BC002548,BC004499,BC009312,BC025339,BX647787,DQ892188,DQ896845,AK004542,AK150204,AK159295,AK169206,AK173224,AK177752,BC053743 | MOV10, Mov10, Mov10 | AAGGTGAACTTTACCTTCA |
| NM_001003819,NM_001003827,NM_021616,NM_130389,AB039902,AB039903,AF220143,AF220144,AK027876,AL583914,BC136871 | TRIM34, TRIM6-TRIM34 | TGCAACTGACTCATCTGCA |
| NM_001003819,NM_001003827,NM_021616,NM_130389,AB039902,AB039903,AF220143,AF220144,AK027876,AL583914,BC136871 | TRIM34, TRIM6-TRIM34 | TGAGTTATGAGAGATGCTT |
| NM_001003819,NM_001003827,NM_021616,NM_130389,AB039902,AB039903,AF220143,AF220144,AK027876,AL583914,BC136871 | TRIM34, TRIM6-TRIM34 | GAGTTATGAGAGATGCTTA |
| NM_001003818,NM_058166,AF220030,AK023210,AK027664,AK290172,AK293295,AK298301,BC047564,BC065575,CR624250,CR749260,CX760618 | TRIM6, TRIM6-TRIM34 | TGGGTTGACGTGACCCTGA |
| NM_005858,AK308251,BC037270,Y11997 | AKAP8 | AGCTACAGCTACGACTATG |
| NM_005082,AB208994,AK310720,BC016924,BC042541,CR617665,D21205,CF121587 | TRIM25 | CCGAACTCAACATCTCTCA |
| NM_003467,AF025375,AF348491,AK129916,AK296674,AK312244,AY242129,BC020968,BT006660,CR594428,CR594588,CR596547,CR598713,CR601301,CR605131,CR610268,CR614199,CR614594,CR614663,CR619476,CR623838,D10924,EU831811,EU831888,L01639,L06797,M99293,X71635 | CXCR4 | CAGTATATACACTTCAGAT |
| NM_000864,AF498979,BC007720,BT007027,DQ893875,EU176172,M81589 | HTR1D | CCCTCGGTGTTGCTCATCA |
| NM_001974,AK131562,AK290401,AK291518,AK313495,BC059395,X81479 | EMR1 | CGCTAAAAGACACCAGGTT |
| NM_001130079,NM_020963,AB046851,AK023297,AK057353,AK074174,AK226011,AL832805,AL833353,BC002548,BC004499,BC009312,BC025339,BX647787 | MOV10 | TGGGGAGAATGACACATCA |

| | | |
|---|---|---|
| NM_001008540,NM_003467,NM_022205,AF025375,AF147204,AF348491,AK129916,AK296674,AK312244,AY242129,BC020968,BT006660,CR594428,CR594588,CR596547,CR598713,CR601301,CR605131,CR610268,CR614199,CR614594,CR614663,CR619476,CR623838,D10924,EU831811,EU831888,L01639,L06797,M99293,X71635,U59760,AF452185,BC089804,U54791,U90610 | CXCR4, Cxcr4, Cxcr4 | TCCAAGCTGTCACACTCCA |
| NM_001003819,NM_001003827,NM_021616,NM_130389,NM_130390,AB039902,AB039903,AB039904,AF220143,AF220144,AK027876,BC136871 | TRIM34, TRIM6-TRIM34 | AGGAGTTAGGACCAGAAGA |
| NM_005858,AK308251,AK309529,AL050160,BC037270,Y11997 | AKAP8 | AGCACCAAGCTGCCCGACA |
| NM_001130079,NM_020963,AB046851,AK023297,AK057353,AK074174,AK226011,AL832805,AL833353,BC002548,BC004499,BC009312,BC025339,BX647787 | MOV10 | TGCCTGACCCTGAACCAGA |
| NM_001008540,NM_003467,AF025375,AF147204,AF348491,AK129916,AK296674,AK312244,AY242129,BC020968,BT006660,CR594428,CR594588,CR596547,CR598713,CR601301,CR605131,CR610268,CR614199,CR614594,CR614663,CR619476,CR623838,D10924,EU831811,EU831888,L01639,L06797,M99293,X71635 | CXCR4 | CAGATAACTACACCGAGGA |
| NM_001003819,NM_001003827,NM_021616,NM_130389,AB039902,AB039903,AF220143,AF220144,AK027876,AL583914 | TRIM34, TRIM6-TRIM34 | CAGGTACATCAAGGAATCA |
| NM_003150,NM_139276,NM_213662,AK092965,AK291933,AK297994,AK301200,BC000627,BC014482,L29277 | STAT3 | ACGGAAGCTGCAGAAAGAT |
| NM_001032394,NM_001032395,NM_020455,NM_198569,AB183546,AB183547,AB183548,AB183549,AF216967,AK027843,AK075087,AK299785,BC036008,BC075798,BX640873,BX640971,BX648315 | GPR126 | TGGAAGAGTTGTCATCACT |
| NM_006398,BC012472,Y12653 | UBD | AGTGACATGATTTTTACTA |
| NM_001003819,NM_001003827,NM_021616,NM_130389,NM_130390,AB039902,AB039903,AB039904,AF220143,AF220144,AK027876,BC136871,BC156770 | TRIM34, TRIM6-TRIM34 | TGCATCACTGTGAGCAACA |
| NM_001003819,NM_001003827,NM_021616,NM_130389,AB039902,AB039903,AF220143,AF220144,AK027876,AL583914,BC136871 | TRIM34, TRIM6-TRIM34 | GAGAGATGCTTATTTATTC |
| NM_005567,XM_001726123,AK055977,AK057776,AK293183,AK301444,AK307665,BC002403,BC002998,BC015761,CR606873,CR613600,L13210,X79089 | LGALS3BP, LOC100133842 | TGGTCTGAGGTCATTAAAA |
| NM_002675,NM_033238,NM_033246,NM_033249,NM_008884,NM_178087,AB209411,AF230401,AF230406,AF230407,AF230411,AK096663,AK300256,BC139795,M73778,M79462,X63131,AK028044,AK032799,AK052305,AK137538,AK156279,BC020990,U33626 | PML, Pml | ACCTCAAGATTGACAATGA |
| NM_005858,AL050160,BC037270,Y11997 | AKAP8 | CCAGAATATGCTGTAATCT |
| NM_017414,XM_001126794,XM_001128283,XM_001128502,XM_001129362,AF176642,AJ243526,AK313385,AL136690,BC014896,BT006835,CR604420,CT841507,CU013072 | LOC727996, LOC728216, LOC728253, LOC728438, USP18 | ACATGAAGATGGAGTGCTA |
| NM_001128166,NM_001128167,NM_001128168,NM_001128172,NM_001128173,NM_002578,AB102659,AF068864,AF155651,AK128022,AK290504,AK315284,AM943850,AM943851,AM943852,BC117353,BC117355,BC152761,BQ268758 | PAK3 | CCACTGAGGATGAATAGTA |
| NM_017414,XM_001126794,XM_001128283,XM_001128502,XM_001129362,AF176642,AJ243526,AK313385,AL136690,BC014896,BT006835,CR604420,CT841507,CU013072 | LOC727996, LOC728216, LOC728253, LOC728438, USP18 | TACATGAAGATGGAGTGCT |
| NM_000864,AF498979,BC007720,BT007027,DQ893875,EU176172,M81589 | HTR1D | AGAGGATTTCTGCTGCTCG |
| NM_001008215,AK131034,CR590322 | C2orf64 | AGAAGAAAAACCTAATTGA |
| NM_016582,AF135600,AK130141,BC025710,BC037974,CR607140,CR615238,CR623241 | SLC15A3 | CAAGTGTCCTCTATGCTTA |
| NM_002198,NM_008390,AB209624,BC009483,CR594837,CR626580,X14454,AK152005,AK152104,AK155983,AK157347,BC003821 | IRF1, Irf1 | TGATCAGAGGTGTACACTA |
| NM_006993,AF081280,AY049737,BC041067,BC054868,CR601718,CR603429,CR621461,BM739536,BM765409 | NPM3 | CCGGTCACTATGGACAGTT |
| NM_001008215,AK131034,CR590322 | C2orf64 | AAAAGATCAGTGTTGGATA |
| NM_004172,AF070609,AK057823,BC037310,CR601776,D26443 | SLC1A3 | AACTTAATTCTGTTACACA |
| NM_006993,AF081280,AY049737,BC041067,BC054868,CR601718,CR603429,CR621461,BM739536,BM765409 | NPM3 | CGGTCACTATGGACAGTTT |
| NM_016582,AF135600,AK127216,AK130141,BC025710,BC037974,CR607140,CR615238,CR623241 | SLC15A3 | CGGATGGACCTCTACTTCT |
| NM_016582,AF135600,BC037974,CR607140,CR615238,CR623241 | SLC15A3 | TGGTTTTACTGGAGCATCA |
| NM_006993,AF081280,AY049737,BC041067,BC054868,CR601718,CR603429,CR621461 | NPM3 | TGGATAAAGGTTGAAATAA |
| NM_016582,AF135600,BC025710,BC037974,CR607140,CR623241 | SLC15A3 | AACCTCGTGCTGTACCTCA |
| NM_016582,AF135600,BC037974,CR607140,CR615238,CR623241 | SLC15A3 | CGCTTCTTCAACTGGTTTT |
| NM_001008215,AK131034,CR590322 | C2orf64 | GAGTTGTTTTCGAATATGT |
| NM_004172,AK293886,AK295032,AK312304,AY954110,BC037310,D26443,L19158,U03504 | SLC1A3 | AGGACAATGAAACTGAGAA |
| NM_016582,AF135600,BC025710,BC037974,CR607140,CR615238,CR623241 | SLC15A3 | GCGCTCAGCCTGCTGCTCT |
