## Supplementary material for "Sequence diversity in the 3’ untranslated region of alphavirus modulates IFIT2-dependent restriction in a cell type-dependent manner": Table S6

| 3'UTR mutant | Lineage | Epizootic clade | Strain accession number(s) |
|---|---|---|---|
| IC | Colombia, western Venezuela, northern Peru | Colombia/Venezuela 1962-64, 1995 | U55342 |
| | | | KC344528 |
| | | | L04653 |
| | | | U55345 |
| | | | AF375051 |
| | | | KC344512 |
| | | | U55350 |
| | | | KF985959 |
| | | | U55347 |
| | | | KP282671 |
| | | | AY986475 |
| | | | AY973944 |
| ID | Colombia, western Venezuela, northern Peru | - | KC344519 |
| | | | KC344477 |
| | | | KC344484 |
| | | | KC344524 |
| | | | KC344522 |
| | | | KC344486 |
| | | | KC344525 |
| | | | KC344520 |
| | | | KC344514 |
| | | | KC344487 |
| | | | KC344502 |
| | | | KC344521 |
| | | | KC344429 |
| | | | KC344462 |
| | | | KC344459 |
| | | | KC344460 |
| | | | KC344509 |
| | | | KC344508 |
| IE | Central America/Mexico | - | KC344481 |
| | | | KC344463 |
| | | | KC344451 |
| | | | KC344478 |
| | | | KC344480 |
| | | | KC344450 |
| | | | KC344493 |
| | | | KC344446 |
| | | | KC344440 |
| | | | KC344448 |
| | | | KC344447 |
| ID mut1 (U11354C) | Colombia, western Venezuela, northern Peru | - | AF004458 |
| | | | AF004472 |
| ID mut2 (U11354C+A11360U) | Colombia, western Venezuela, northern Peru | Venezuela, 1992-93 | AF004459 |
| | | | U55360 |
| | | - | MF459684 |
| | | | KC344513 |
| | | | AF100566 |
| ID mut3 (U11355C+U11357C+C 11364U) | Panama, Peru | - | KC344510 |
| | | | KC344511 |
| | | | KC344474 |
| | | | KC344475 |
| ID mut4 (U11355C+C11364U+C 11366U) | Panama, Peru | - | KC344490 |
| | | | KC344526 |
| | | | KC344506 |
| | | | KC344518 |
